# A novel fragmented mitochondrial genome in the protist pathogen *Toxoplasma gondii* and related tissue coccidia

**DOI:** 10.1101/2020.05.16.099366

**Authors:** Sivaranjani Namasivayam, Rodrigo P. Baptista, Wenyuan Xiao, Erica M. Hall, Joseph S. Doggett, Karin Troell, Jessica C. Kissinger

**Author notes:** Corresponding Author: Jessica C. Kissinger, University of Georgia, 500 D.W. Brooks Drive, Coverdell Center, Athens, GA, 30602, USA.

## Abstract

Mitochondrial genome content and structure vary widely across the eukaryotic tree of life with protists displaying extreme examples. Apicomplexan and dinoflagellate protists have evolved highly-reduced mitochondrial genome sequences, mtDNA, consisting of only 3 cytochrome genes and fragmented rRNA genes. Here we report the independent evolution of fragmented cytochrome genes in *Toxoplasma* and related tissue coccidia and evolution of a novel genome architecture consisting minimally of 21 sequence blocks (SBs) that exist as non-random concatemers. Single-molecule Nanopore reads consisting entirely of SB concatemers ranging from 1-23 kb reveal both whole and fragmented cytochrome genes. Full-length cytochrome transcripts including a divergent *coxIII* are detected. The topology of the mitochondrial genome remains an enigma. Analysis of a *cob* point mutation reveals that homoplasmy of SB’s is maintained. Tissue coccidia are important pathogens of man and animals and the mitochondrion represents an important therapeutic target. Their mtDNA sequence has remained elusive until now.

## Introduction

Mitochondria are notable for their incredible diversity in genome content and genome topology (Burger et al. 2003; Flegontov and Lukes 2012; Kolesnikov and Gerasimov 2012). The endosymbiont genome sequence that gave rise to the mitochondrial genome sequence, mtDNA, was likely circular, characteristic of its prokaryotic origins (Lang et al. 1997). However, increasing bodies of literature indicate that this structure is not universal and that considerable evolution of both genome structure and gene content has occurred in both protists and multicellular eukaryotes (Burger et al. 2003; Gissi et al. 2008). At the time of its discovery, the mtDNA of the apicomplexan *Plasmodium falciparum* was the smallest mtDNA known at 5,967 bp and among the most unusual with only 3 protein-encoding genes (cytochrome oxidase subunit I, *coxI*: cytochrome oxidase subunit III, *coxIII*; and cytochrome b, *cob*) and fragmented rRNA genes with many rRNA portions missing (Vaidya and Arasu 1987; Feagin 1992; Feagin et al. 1997; Wilson and Williamson 1997; Feagin et al. 2012). At the other end of the size spectrum, the mtDNA of angiosperms ranges from 200 kb to 11 Mb in size and shows a huge variation in structure, content, gene/DNA transfers and processes such as RNA editing (Gualberto and Newton 2017; Petersen et al. 2017). Mitochondrial genomes consisting of multiple divergent circular molecules have been noted in the fungus *Spizellomyces* (Burger and Lang 2003), *Columbicola* feather lice (Sweet et al. 2019) and certain cnidarian parasites (Yahalomi et al. 2017). Among the protists, the kinetoplastid mtDNA (called kDNA) consists of multiple gene-encoding maxi-circles and guide RNA encoding minicircles that form a tight physical network, the kinetoplast (Morris et al. 2001). The mtDNA of the symbiotic protist *Amoebidium parasiticum* is comprised of hundreds of distinct linear molecules with a common pattern of terminal repeats (Burger et al. 2003).

The alveolates have their own non-canonical mtDNA sequences and structures (Flegontov and Lukes 2012). The ciliates, for example, contain a 47 kb linear mtDNA flanked by telomere-like repeats with a number of ciliate-specific open reading frames (ORFs) of unknown function (Slamovits et al. 2007; Nash et al. 2008; Waller and Jackson 2009; Flegontov and Lukes 2012). The mtDNA of dinoflagellates (sister group to the Apicomplexa) share the reduced gene content and fragmented rRNA genes with the Apicomplexa but their protein-encoding genes are highly-fragmented, highly-repetitive, contain multiple non-identical mtDNA molecules, utilize non-canonical start codons and have evolved trans-splicing and RNA editing (Slamovits et al. 2007; Nash et al. 2008; Waller and Jackson 2009; Flegontov and Lukes 2012) (Fig. 1).

**Fig. 1.**
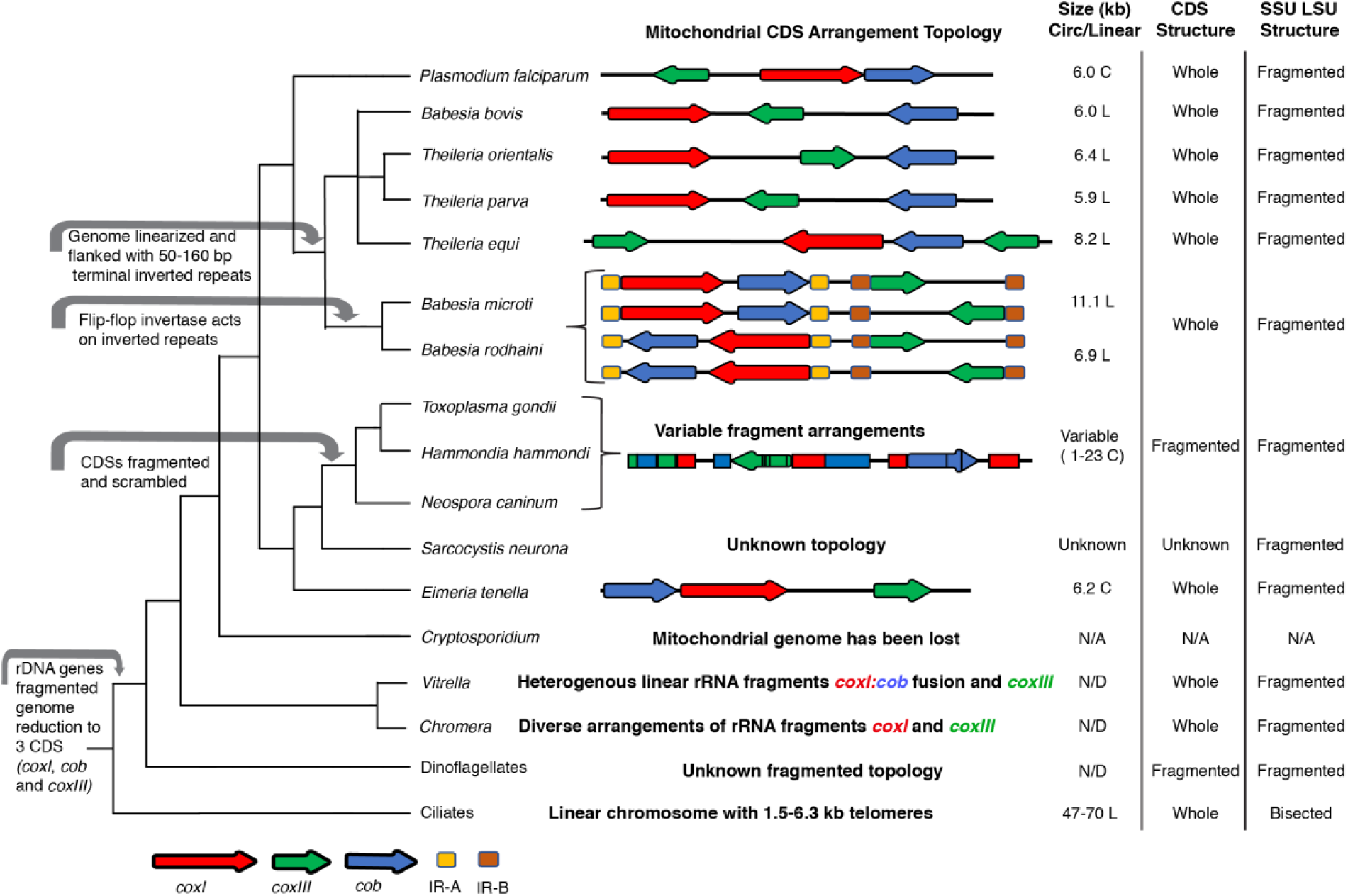
Alveolate mtDNA evolution and characteristics. Left – Cladogram of alveolate relationships with major mitochondrial genome events indicated with grey arrows. Center – Schematic of mtDNA CDS (red, green and blue arrows) or inverted repeats (gold and brown) indicated. Spacing is approximate with CDS lengths exaggerated for ease of viewing. Ribosomal RNA and other RNA genes or fragments thereof are not represented. Right – mtDNA size and topology (L - Linear; C – Concatemer, presumably from circular progenitors); status of CDS sequences and status of SSU and LSU rRNA are indicated.

While all sequenced apicomplexan mtDNAs contain only three protein-coding genes (*coxI, coxIII* & *cob*) and show a high level of rRNA gene fragmentation like the dinoflagellates, the orientation and arrangement of these genes varies across the phylum, as does the genome architecture (Creasey et al. 1993; Hikosaka et al. 2010; Hikosaka et al. 2011; Hikosaka et al. 2012; Hikosaka et al. 2013) (Fig. 1). The *Theileria* mtDNA is a 7.1 kb linear monomer with telomere-like termini (Kairo et al. 1994). Members of the genus *Babesia* have linear mtDNA monomers with a dual flip-flop inversion system that results in four different sequences that are present in equal ratios in the mitochondrion (Hikosaka et al. 2011). The coccidian *Eimeria*, the species closest to *Toxoplasma gondii* with a published genome sequence and structure, is detected as a linear concatemer of a repeating 6.2 kb sequence similar to that of *Plasmodium* albeit with the genes in a different order.

The mitochondrial genome of the coccidian protist pathogen *Toxoplasma gondii*, has been enigmatic and elusive. *Toxoplasma gondii* is an incredibly successful zoonotic generalist pathogen which infects 30-50% of the human population (Flegr et al. 2014) but primarily causes illness in the immunocompromised, or fetuses as a result of transplacental infections (Montoya and Liesenfeld 2004; Dubey 2010). Attempts to assemble the mitochondrial genome from the various organismal genome projects failed and efforts to isolate the single mitochondrion organelle and identify the complete mtDNA sequence and structure have faced numerous challenges. Attempts to elucidate the mtDNA sequence using sequenced-based approaches, hybridization probes, PCR amplification or assembly from Sanger or next-generation short read genome sequence projects were unsuccessful due to the large number of nuclear-encoded sequence fragments of mitochondrial origin (NUMTs) present in the nuclear genome (Minot et al. 2012; Lau et al. 2016; Lorenzi et al. 2016). The NUMT sequences, which display a range of degeneration relative to the organellar mtDNA as they decay with evolutionary time following insertion, cross hybridize and interfere with signals from molecular-based mtDNA isolation and amplification methods. They also interfere with assembly algorithms for short-read sequences (Sanger and Illumina) that were used to generate the numerous existing *Toxoplasma* genome sequences (Minot et al. 2012; Lau et al. 2016; Lorenzi et al. 2016). Physical attempts to isolate the mitochondrion have also failed thus far. *In vitro*, the single mitochondrion of *T. gondii* is physically polymorphic and occurs as an elongated S, lasso-shaped or boomerang-shaped organelle when the parasite is located inside of a host cell and polymorphic topologies are observed in the extracellular matrix, making physical isolation of the mitochondrion from cellular fractions nearly impossible (Nishi et al. 2008), but mitochondria-enriched fractions can be obtained.

Despite the community’s inability to isolate and determine a mitochondrial genome sequence for this important pathogen multiple lines of evidence suggest the *T. gondii* mitochondrion is functional. First, rhodamine 123 and MitoTracker dyes which are markers of a transmembrane potential across the mitochondrial membrane indicate the mitochondrion is active (Tanabe 1985; Melo et al. 2000; Weiss and Kim 2007). Second, a complete set of tRNAs are imported into the *T. gondii* mitochondrion (Esseiva et al. 2004). Third, the mitochondrial protein *cob* has been characterized and is an excellent drug target (Vaidya et al. 1993). Mutations in *cob* were found to be associated with atovaquone (McFadden et al. 2000) and Endochin-like quinolone (ELQ-316) (Alday et al. 2017) drug resistance. Fourth, NUMTs identified in the current *T. gondii* ME49 nuclear genome sequence, even if combined together cannot functionally encode any of the cytochrome genes in their entirety. Thus, the genes must be encoded in the mitochondrion (Namasivayam 2015).

In the present work we employ Sanger, Illumina and single-molecule Oxford Nanopore sequencing of DNA and RNA together with molecular and bioinformatics strategies to better understand the sequence and structure of the mtDNA of *T. gondii*. We show how long-read single-molecule Nanopore sequences elucidated a novel mtDNA architecture for *T*. gondii based on concatenated sequence blocks (SBs) that do not appear to be the direct result of rolling-circle replication. The SBs are evolutionarily conserved with a few minor variations among the examined Toxoplasmatinae genera (*Toxoplasma, Neospora* and *Hammondia*). We also show that this lineage has independently evolved fragmentation of their cytochrome genes, a feature previously detected only in dinoflagellates (Waller and Jackson 2009; Jackson et al. 2012).

## Results

### 21 Discrete Sequence Blocks Constitute the *Toxoplasma gondii* Mitochondrial Genome

The *T. gondii* ME49 genome sequence project yielded numerous contigs containing portions of, or in some cases, complete cytochrome genes and mtDNA rRNA gene fragments (Gajria et al. 2008; Lorenzi et al. 2016). These contigs did not assemble into a single mtDNA sequence. However, the sequence information contained in these contigs as well as individual unassembled *T. gondii* genomic and EST reads (Sanger chemistry) that matched mtDNA from related species, served as a template for PCR primer design (Table S1). The primer design carefully avoided NUMTs which are located in the nuclear genome and degenerate relative to the putative mtDNA. The initial strategy was to use a PCR-based approach on DNA and cDNA obtained from mitochondria-enriched cell fractions as template. We performed PCR under very stringent conditions using several combinations of primer pairs. The results were puzzling. No amplicon was larger than ∼3.5 kb. Primer pairs often generated multiple amplicons (Fig. S1A, lanes 6-11), that when sequenced revealed a high level of sequence identity to each other in some regions (blocks) but differed in others (Figs. 2A, S2 & Dataset S1). This variety of amplicon sequences prompted the testing of PCR reactions with only a single primer. They too, generated multiple amplicons (Fig. S1B, lanes 1-2). With difficulty, we did identify primers that avoided NUMTs and would amplify a single region (100-350 bp) within each one of the three-cytochrome genes that are typically encoded by apicomplexans, *coxI, coxIII* and *cob* (Hikosaka et al. 2013) (Fig. S1A lanes 2-4).

**Fig. 2.**
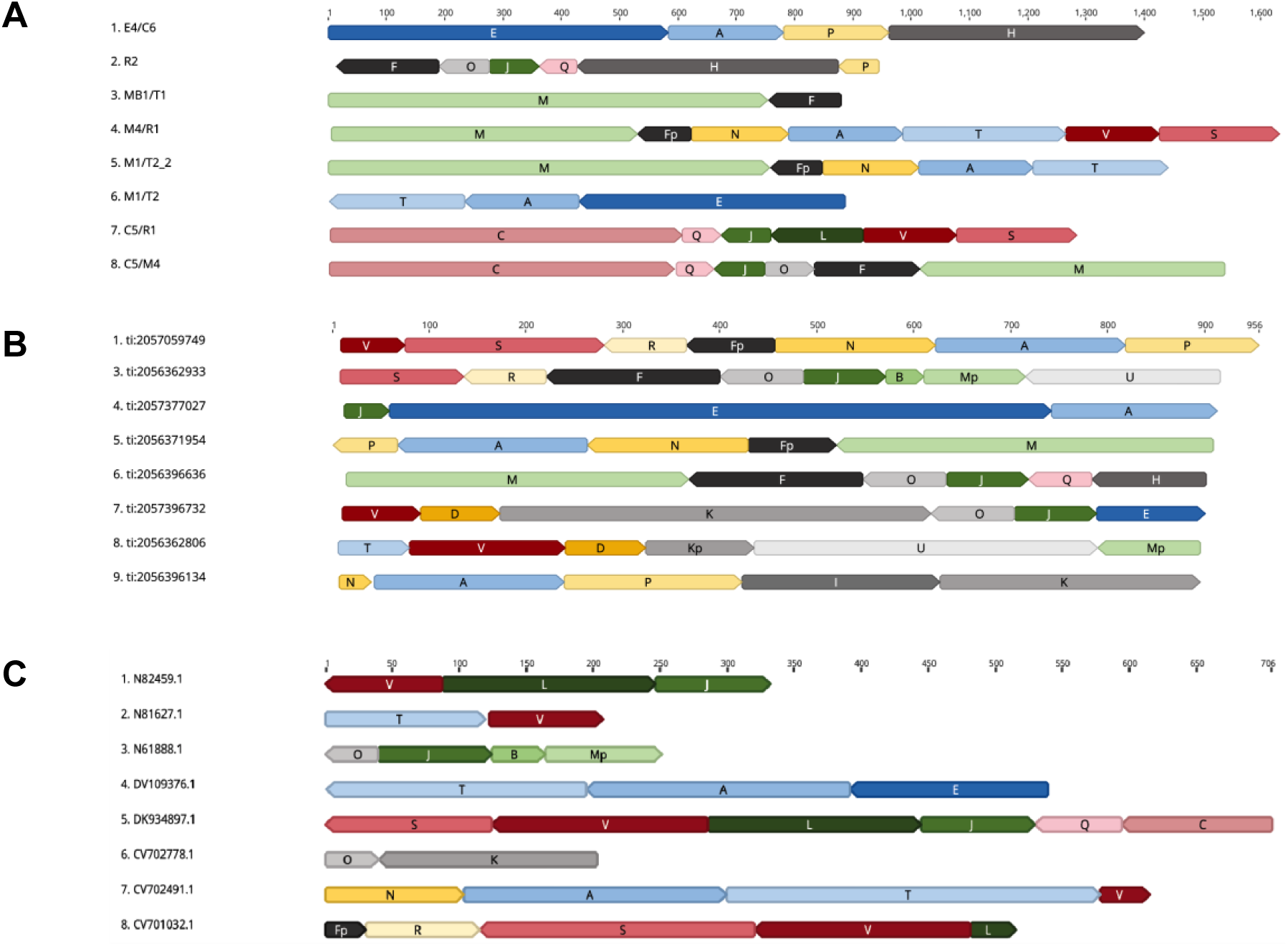
Sequence evidence of fragmented mtDNA. (A-C) Each row of colored blocks represents individual, annotated, (A) mtDNA-specific PCR amplicons; (B) Sanger genomic reads; or (C) EST reads. Sections of reads containing identical sequence to other mtDNA reads are called ‘sequence blocks’, SBs, and are colored and labeled with a unique color and letter corresponding to Fig 3 and Table 1. Shades of blue represent different SB’s found in *cob;* shades of red are SBs found in *coxI;* and shades of green are SBs found in *coxIII*. Orientation of a block is indicated by the point on each block. Multiple sequences are shown for each type. SBs located on the ends of reads may be incomplete. GenBank accession numbers are as indicated in panels B-C. Sequences for the reads in panel A are located in dataset S1. Scale (in bp) is as indicated in each panel.

As PCR is extremely sensitive and capable of amplifying rare artifacts or, in some cases, a NUMT capable of annealing to the primers, we abandoned approaches that relied on amplification or existing assembled sequences and turned our attention to the analysis of existing individual genomic and EST Sanger sequence reads of up to 1 kb present in the NCBI GenBank (Benson et al. 2017). The goal was to identify reads that only contained putative mtDNA or mitochondrial cDNA. Each putative mtDNA read or transcript consisted only of discrete, reproducible, blocks of sequence with similarity to various regions of known apicomplexan mtDNA (Figs. 2B-C; S3-S4). In total, 21 sequence blocks (SB’s) that reproducibly shared 100% sequence identity with parts of the amplicon sequences derived from mitochondrial-enriched cell fractions and portions of unassembled Sanger sequence reads generated during the *Toxoplasma gondii* ME49 genome project were identified. The 21 SBs were named from A to V in the order of their discovery (Fig. 3, Table 1, Dataset S2). Three SBs (F, K & M) were observed to also exist in precise, reproducible, truncated forms, indicated with a “p” for partial, *e*.*g*. Fp (Table 1). Fifteen of the 21 SBs share significant BLASTN or TBLASTX similarity to regions of *Eimeria tenella* and/or *Plasmodium falciparum* mtDNA sequences (Table S2). The SBs range in size from 40 bp to 1050 bp and none encodes a cytochrome or rRNA gene in its entirety (Fig. 3). The SBs were compared to each other with a nucleotide dot plot analysis. The SBs are non-redundant except for the last 50 bp of SB’s C and H, which are identical. Small 10-14 bp regions of microhomology are detected between many SBs (Fig. S5A-B). With distinct, named, SBs identified, all PCR, genomic and EST Sanger reads could be annotated. Annotation revealed that amplicons and Sanger reads contained one or more SBs, one directly after the other, with no intervening sequence, in distinct, non-random permutations (Figs. 2A & S2). No full-length cytochrome genes were observed, but this was not unexpected given the size of the reads. However, many fragments of cytochrome genes were observed in a variety of contexts and orientations. Some SBs were always observed in specific combinations with other blocks. For example, we found block E to always occur between blocks J and A. However, these blocks couldn’t be assembled into a larger SB because J and A each occur in other contexts. We identified similar such permutations of the SBs from amplicons and unassembled genomic and EST Sanger reads (Figs. 2; S3-S4).

**Table 1.** Characterization and conservation of the 21 mtDNA sequence blocks in *T. gondii, N. caninum* and *Hammondia*.

| Block | *T. gondii* ME49 |  | *N. caninum* LIV |  | *Hammondia*<sup>^</sup> |  |
| --- | --- | --- | --- | --- | --- | --- |
|  | Block Length | Entire block found at 100% identity in nuclear genome? | Match to *T. gondii* block |  | Match to *T. gondii* block |  |
|  |  |  | Start | Stop | Start | Stop |
| **A** | 196 | N | 1 | 196 | 1 | 196 |
| **B** | 40 | Y | 1 | 40 | 1 | 40 |
| **C** | 1050 | N | 1 | 1050 | 1 | 1050 |
| **D** | 82 | N | 1 | 82 | 2 | 82 |
| **E** | 683 | N | 1 | 683 | 1 | 683 |
| **F<sup>#</sup>** | 179 | Y | 1 | 179 | 1 | 179 |
| **H** | 447 | N | 1 | 447 | 1 | 438 |
| **I** | 204 | N | 1 | 204 | 1 | 204 |
| **J** | 85 | Y | 1 | 85 | 1 | 85 |
| **K<sup>#</sup>** | 445 | N | 1 | 445 | 1 | 445 |
| **L** | 159 | Y | 1 | 159 | 1 | 159 |
| **M<sup>#</sup>** | 754 | N | 1 | 754 | 104 | 560 |
| **N** | 166 | N | 1 | 166 | 1 | 166 |
| **O** | 86 | Y | 1 | 86 | 1 | 86 |
| **P** | 184 | N | 1 | 184 | 1 | 184 |
| **Q** | 65 | Y | 1 | 65 | 1 | 65 |
| **R** | 85 | Y | 1 | 85 | 1 | 85 |
| **S<sup>\$</sup>** | 205 | Y | 1 | 205 | 1 | 205 |
| **T** | 279 | N | 1 | 278 | 1 | 276 |
| **U** | 354 | N | 1 | 354 | 26 | 354 |
| **V** | 161 | Y | 1 | 161 | 1 | 161 |

**Fig. 3.**
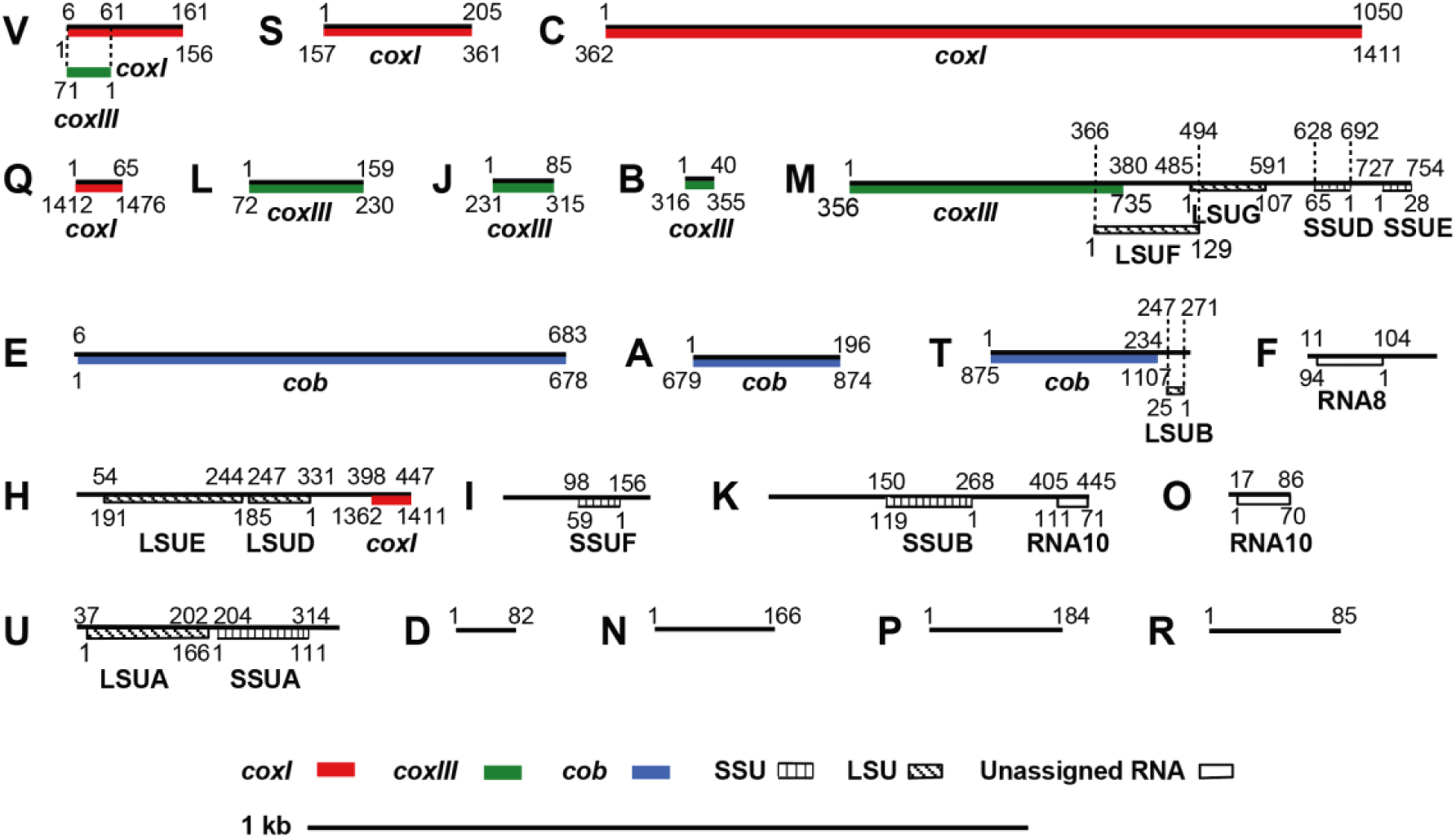
The 21 sequence blocks of the *T. gondii* mtDNA. The identified 21 SBs are represented by a black line, drawn to scale and named with 21 alphabet characters, A to V (there is no sequence block “G”). The blocks are annotated using their nucleotide correspondence to sequences of assembled cytochrome genes and rRNA gene fragments. These annotations are shown on the blocks as defined in the key. The coordinates of a block that encodes a cytochrome or rRNA gene fragment are indicated above the black bar and the corresponding coordinates of the assembled gene or rRNA fragment are indicated below the gene fragment. Portions of sequence block V contribute to *coxI* in the forward orientation and *coxIII* in the reverse orientation. Blocks D, N, P and R could not be annotated as part of a cytochrome gene or rRNA fragment.

All SBs including the partial forms of F, K and M when not located at the beginning or end of a read have the same length and sequence irrespective of the blocks they are flanked by, whether they appear in a PCR product, genomic or EST sequence read (Figs. 2, S2-S4; Table 1). A full-length block C was not be found in the Sanger genomic or EST reads nor in the PCR products, perhaps due to its large size, but it was observed in Oxford Nanopore data described below. Exhaustive searches of Sanger and next-generation sequence read data generated by us, or others, repeatedly identified only these 21 blocks. For example, if a Sanger read was identified that contained a 100% match to a SB, the rest of the sequence in the read was examined. These analyses usually revealed that the entire read contained a subset of the 21 mtDNA SBs, often in different permutations (Figs. S3-4). Occasionally, examination of the rest of the read revealed that the remaining sequence was nuclear and the 100% similarity to the sequence block was due to a recent mtDNA transfer (*e*.*g*. a NUMT) rather than mtDNA. NUMT-containing reads were not analyzed further in this study. The mtDNA SBs are not to be confused with the degenerate mtDNA fragments inserted in the *T. gondii* nuclear genome referred to as NUMTs (nuclear sequences of mitochondrial origin) (Ossorio et al. 1991; Lopez et al. 1994; Namasivayam 2015). NUMTs are insertions of fragments of mtDNA at the location of a nuclear double-strand DNA break (Lopez et al. 1994; Mishmar et al. 2004; Richly and Leister 2004). A NUMT can be derived from any mtDNA region and the boundaries of a NUMT usually do not match the boundaries of the identified mtDNA sequence block(s), but a few exceptions have been noted. NUMTs can begin and end at any mtDNA sequence point. Perhaps the best visualization is one of randomly-sheared mtDNA. mtDNA SBs can be distinguished from NUMTs by their distinct sizes, boundaries and non-varying sequence. NUMTs vary greatly in size, they demonstrate varying levels of sequence decay with respect to the 21 mtDNA SBs (Table 1)(Namasivayam 2015) and they are flanked by non-mitochondrial (nuclear) DNA sequence.

Finally, we generated Oxford Nanopore long-read single-molecule sequences for *T. gondii* ME49 (Table S3) and mined these unassembled and uncorrected data for reads that contained mtDNA SBs. We used Nanopore sequencing precisely because each read is generated from an individual DNA molecule with the addition of any amplification steps. We also searched uncorrected reads to avoid the possibility of correction confounding our detection and disambiguation from NUMTs. We identified dozens of single-molecule reads ranging in length from 1-23 kb that when annotated consisted exclusively of non-random concatemers of the 21 SBs and many had the ability to encode one or more full-length cytochrome genes *e*.*g*. the SBs EAT encode *cob* and can be seen in the top panel of Fig 4. Following identification of the Nanopore mtDNA reads, the reads were corrected with *T. gondii* ME49 Illumina data. It is the corrected Nanopore mtDNA reads that are analyzed and presented in this work (Fig. S6, dataset S3). As an extensive analysis of the Nanopore sequence data did not reveal any new SBs, we believe we have identified all sequences that minimally constitute the mitochondrial genome of *T. gondii*. We refer to the mtDNA sequences as SBs due to their fragmented nature. The 21 blocks total 5,909 bp.

**Fig. 4.**
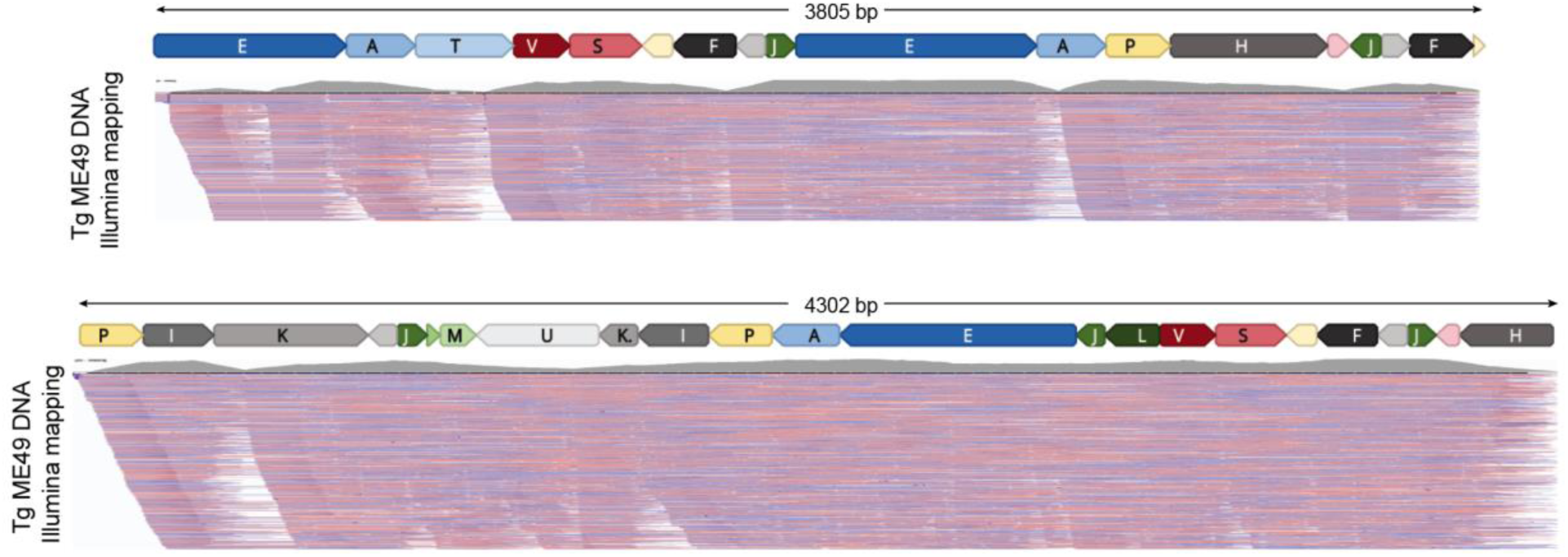
Annotated *Toxoplasma gondii* ME49 Nanopore mtDNA reads. Each panel represents the annotation of a single Nanopore read with SBs. There are no intervening nucleotides between SBs. The length of the read is indicated above the annotation. MtDNA-specific paired-end Illumina reads generated from *T. gondii* ME49 DNA (SRR9200762) were mapped to the Nanopore sequences and the alignment was visualized using the Integrated Genomics Viewer (IGV). Both Illumina read ends were required to map at 100% identity. A histogram of read abundance is shown in grey just below each annotated Nanopore read. Red and blue lines below each histogram indicate the mapped paired-end reads. Not all mapped reads are shown.

### The 21 Sequence Blocks Encode Expected mtDNA Features

The SBs were subjected to extensive analysis as detailed in the materials and methods. They were used to search the GenBank with BLASTN and BLASTX. All ORFs > 50 amino acids from both the sequence blocks and several long Nanopore reads were determined and searched with InterProScan and the SBs were used to search RFAM to detect and annotate rRNA and other ncRNAs. No proteins other than *coxI, coxIII* & *cob* were identified. Twelve of the 21 SBs encode portions of the three expected cytochrome genes and eleven, when present in a particular order, encode: *coxI* (blocks V, S, C, Q); *coxIII* (blocks <u>V</u> [reverse orientation indicated by underline], L, J, B, and the first half of M) and *cob* (blocks E, A, T) (Fig. 3). Three blocks encode portions of a cytochrome and fragments of rRNA genes (blocks H, M and T) (Fig. 3). The replicated small portion of *coxI* at the end of sequence block H does not appear to contribute to a functional cytochrome. Six blocks (F, H, I, K, O, U) encode LSU and SSU rRNA gene fragments and genes for well-conserved apicomplexan RNAs (RNA8 and RNA10) although, it should be noted that RNA10 is split onto SBs (O and K) and the short 22 nt LSUC fragment observed in *Plasmodium falciparum* could not be detected. Currently, 4 blocks remain unannotated and have no discernable function (blocks D, N, P, R) (Fig. 3), although two have misleading BLAST hits to annotated nuclear genes where NUMTs containing portions of these SBs are located. The majority, but not all of the gene typically encountered in an apicomplexan mtDNA are present. Several RNA genes observed in other apicomplexans (Feagin et al. 2012) have not been identified (Fig. 3). Overall, 3,333 bp of the SBs are annotated as protein encoding and 1,290 bp are annotated as rDNA. Searches of the SBs and the longest *T. gondii* ME49 Nanopore read for additional RNA or protein features did not reveal any additional genes.

Full-length cytochrome genes are detected in the Nanopore reads and can be annotated when particular SBs appear in the correct order and orientation (Figs. 5, S6). To confirm that the full-length cytochrome sequences are routinely present, we mapped paired-end *T. gondii* ME49 Illumina genomic reads and paired-end RNA-seq data from *T. gondii* CZ to the full-length sequences encoding *coxI, coxIII* and *cob* (Fig. 5) and all data (including RT-PCR and RNA Nanopore, see below) support the presence of full-length cytochrome genes. The RNA-seq evidence is less strong but it is important to note that poly-A+ RNA was purified for these libraries. The encoded *T. gondii* mtDNA cytochrome proteins are comparable in length and sequence to other apicomplexan cytochrome genes (Fig. S7, Dataset S4) but there is some doubt about the exact beginning of the *cob* open reading frame as two potential starts are possible, one 9 amino acids longer than the other (Fig. S7). Block V contributes to two different cytochrome genes. All of block V is utilized near the 5’ end of *coxI* (blocks V, S, C, Q). However, only a portion of sequence block V in the reverse orientation (indicated by underline) is required to encode *coxIII* (blocks <u>V</u>, L, J, B, M) (Figs. 3 & 5). COXI and COB are well conserved but COXIII is less well conserved (Fig. S7). Each sequence begins with a methionine (using mitochondrial translation tables) and ends with a canonical stop codon. There are no spliceosomal introns located in the full-length coding sequences no indication that RNA editing is required to produce the coding sequences.

**Fig 5.**
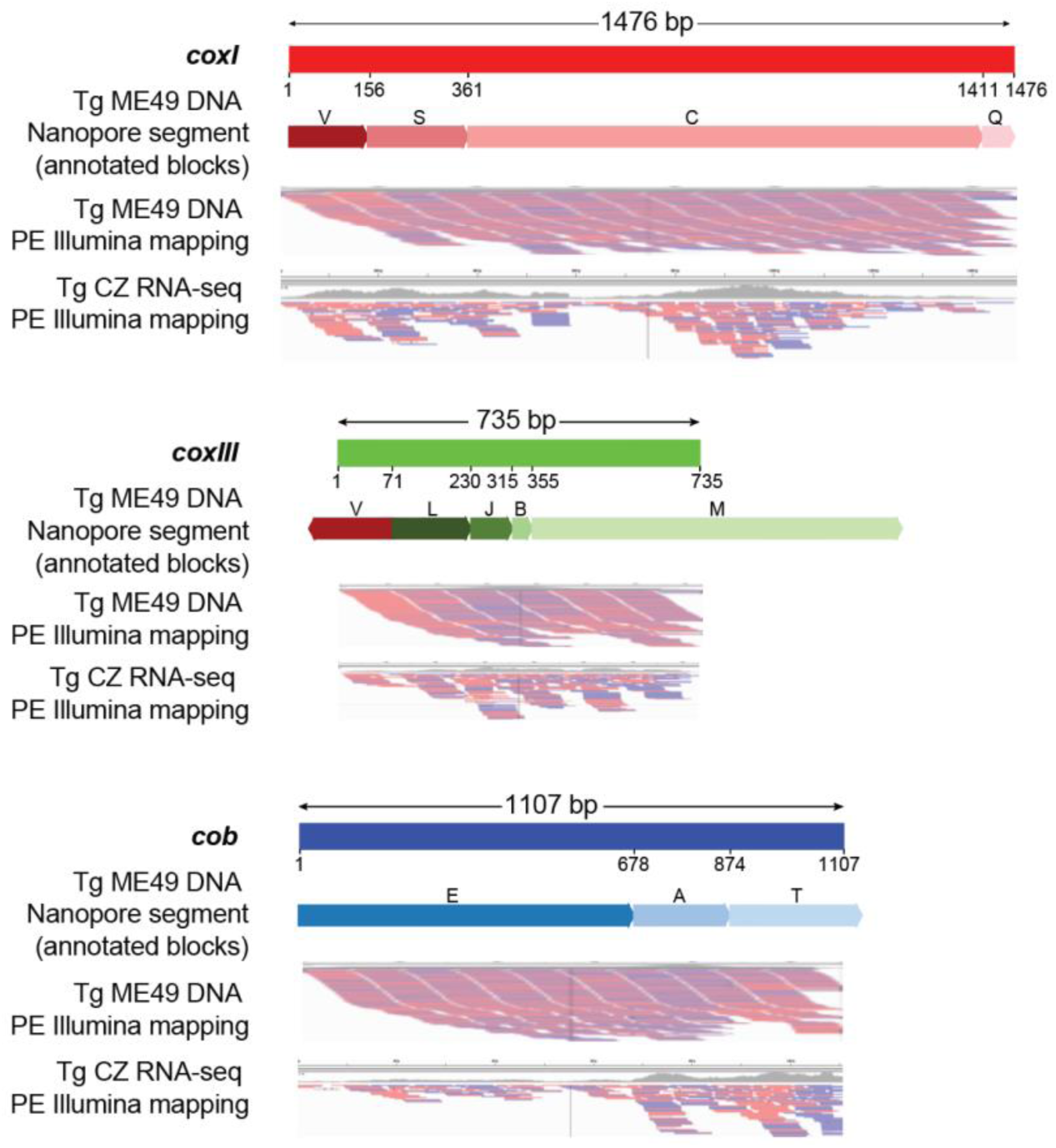
Full-length protein-encoding genes in the *T. gondii* mt genome are supported. Segments of a Nanopore read that can encode a full-length cytochrome gene are annotated with their sequence block name and shown below the schematic of each gene. Numbers below each gene schematic represent nucleotide start/stop positions of a sequence block on the gene. All blocks are in the forward orientation except block V in *coxIII*. All blocks shown participate fully in creation of the cytochrome sequence except block V where only a part of *coxIII* is used, and sequence blocks M and T where only a portion of the block contributes to encoding a cytochrome gene. MtDNA-specific paired-end Illumina DNA (SRR9200762) and RNA-seq (SRR6493545) reads were independently mapped to each of the cytochrome gene sequences and visualized using IGV. Red and blue lines below each gene indicate the mapped Illumina paired-ends. Both ends were required to map.

The cytochrome genes are not detected at similar frequencies. In order to estimate *T. gondii* ME49 relative gene copy numbers we mapped genomic Illumina paired-end data from two different Illumina data sets, one ME49 HiSeq and one RH-88 MiSeq, to assembled full-length cytochrome gene sequences and a single-copy nuclear gene, GAPDH. Both ends of a paired read were required to match the target sequence 100%. The calculated GAPDH coverage for ME49 of 1195 X and RH-88 of 89 X was set to equal 1 copy and used to normalize the coverage of *coxI, coxIII, cob* and a few other often observed sequence block arrangements. *coxI, coxIII* and *cob* were detected at 316, 423 and 297 X for ME49 relative to GAPDH and 369, 523 and 313 X for RH-88 (Table S4). For comparison, the *E. tenella* mtDNA was estimated to exist at ∼50 copies per nuclear genome (Hikosaka et al. 2011) (Fig. 1). As a negative control, sequence blocks that are never observed to occur together, *e*.*g*. A,B,D,E and K,M,V were manually concatenated and Illumina reads were mapped as above. The negative controls do show mapped read coverage ranging from 150-200 X (Table S4) but, visualization of the reads revealed they were mapping within the longer sequence blocks, rather than across the SBs with the exception of 33 reads that spanned sequence blocks K and M (data not shown). Arrangements of SBs that do not encode cytochrome genes are also observed in Nanopore reads, Sanger EST and genome sequence reads (Figs. S3-4; S6) and are supported by Illumina reads (*e*.*g*. Fig. 4 SBs EAP). In fact, about 1/3 of the observed Nanopore sequences (length 2.1-11 kb) do not encode a full-length cytochrome despite containing numerous SBs that contain portions of a cytochrome gene (Fig. S6).

Similar to other apicomplexan mitochondrial genome sequences, the rRNA genes in *T. gondii* are fragmented. We were able to identify a minimum of 14 ribosomal fragments encoded in the 21 SBs (Fig. 3) based on sequence similarity to the *P. falciparum* and *E. tenella* LSU and SSUrRNA gene fragments (Table S2) and alignment to the *E. coli* SSU and LSU secondary structures (Fig. S8). Blocks M, K and H contain multiple rRNA gene fragments. Comparison of rRNA gene fragment order in each of these three SBs to the order of the corresponding fragments in *E. tenella* or *P. falciparum* revealed that the order is not conserved (Table S2).

### Cytochrome Transcripts are Identified for all Three Cytochrome Genes

Cytochrome b is well studied in *T. gondii* and several cDNAs have been reported in the literature and GenBank (McFadden et al. 2000). A partial *coxI* transcript has been reported. However, a *coxIII* transcript is not present in the GenBank. Analysis of *T. gondii* EST Sanger sequence data identified numerous reads consisting fully of mtDNA SBs, some of which can partially encode a cytochrome suggesting cytochrome genes are indeed being transcribed. All three cytochrome genes can be assembled from multiple Sanger ESTs as well as Illumina RNA-seq data (Fig. 5, Table S5). Using RT-PCR we were able to generate a nearly full-length *cob* and *coxIII* and a partial *coxI* (Dataset S5). Ultimately, examination of available *T. gondii* PRU RNA sequence data from Oxford Nanopore (kindly provided by Stuart Ralph) revealed numerous reads capable of encoding full-length cytochrome transcripts (except for a few missing base pairs at read ends), once corrected. Of the 30 longest error-corrected RNA Nanopore molecules that consisted of only mtDNA SBs, 3 reads were capable of encoding *cob*; 6 *coxI* and 2 *coxIII* (Table S6; Dataset S6). We observe that the *coxI* reads begin at the beginning of the gene and appear to only encode *coxI* (blocks V, S, C, Q) or *coxI* followed by sequence block J (V, S, C, Q, J) whereas the other transcripts encoded numerous SBs in addition to the cytochrome coding sequence. As was observed with the Sanger ESTs, numerous transcripts that consisted of some SBs that together do not encode a full-length cytochrome as well as other non-random arrangements of SBs were detected. For example, Oxford Nanopore direct RNA strand sequences encoding sequence blocks ‘<u>I, P, A, N,</u> Fp, <u>R,</u> S, <u>V,</u> L, B, Mp, <u>U, Kp</u>’ (TgNano RNA2) and ‘N, A, T, V, D, Kp, U, <u>Mp, B, J,</u> O, <u>K</u>’ (TgNano_RNA4) were observed (Table S6). Together, these analyses reveal that both cytochrome genes as well as partial and non-cytochrome arrangements of SBs are transcribed.

The *coxI* transcripts that are not followed by a nearly full block J are polyadenylated, but the pattern is unusual. An additional 12 nucleotides of sequence, specifically the last 12 nucleotides of sequence block J in the reverse orientation (5’ – CACAATAGAACT) are present following the *coxI* TAA stop codon. These 12 nucleotides are followed by a poly(A) stretch of variable length, suggesting that sequences present in J may serve as a degenerate poly(A) signal since sequence block J contains sequence with only a single base difference from the canonical AAUAAA poly(A) signal. Variant polyadenylation signals have been observed in humans, including the variant seen here (Beaudoing et al. 2000). The remainder of sequence block J is capable of forming a prominent but imperfect, stem loop (Fig. S9) but the rest of the sequences do not appear to contain long runs of obvious hairpin structures as evidenced by the dot plots (Fig. S5). Examination of the three detected polyadenylated *coxI* transcripts showed that following the poly(A) stretch there is addition of apparently random sequence and in one transcript this random sequence is followed by another poly(A) tail (Table S6). The origin of the random sequence in the poly(A) tails is unknown. There is no evidence of polyadenylation of any other transcripts, coding or not, based on the available sequences (Table S6). Therefore, it is possible the lack of polyadenylation in the mitochondrial transcripts may be affecting the presence of mitochondrial reads in the Illumina RNA-seq data as sequencing libraries are typically generated following oligo(dT) purification (Fig. 5). Nevertheless, RT-PCR and read data from multiple sequencing technologies indicate that all three cytochromes in *T. gondii* are transcribed.

### Homoplasmy of Sequence is Maintained

Comparative analyses of the multiple copies of the 21 SBs revealed that all internally located SBs, *i*.*e*. blocks not at the end of a sequence read (where incomplete blocks are often present) are identical in sequence, barring a few SBs that contain small aberrations characteristic of rearrangement artifacts (Tables S6, S7). To further assess homoplasmy in the context of this unusual genome architecture, a recent N-ethyl-N-nitrosourea, (ENU), *cob* mutant named ELQ-316 (Alday et al. 2017) was sequenced with Oxford Nanopore technology. Parasites from an early passage of this newly created mutant were kindly provided by Stone Doggett. This strain contains a point mutation in mtDNA sequence block E which encodes a Thr222 -> Pro amino acid substitution. Sequence block E is only found flanked by sequence blocks J and E arranged as, J, E, A. (Table 2). If homoplasmy does exist, we expect this mutation to be present in all copies of block E. In order to not bias the results, only uncorrected Nanopore reads were analyzed. 779 mtDNA-only reads were identified. Of the 138 sequence block Es identified that contained the mutated region, 132 contained the point mutation, indicating homoplasmy is maintained. The 6 regions that did not show the mutation are well within the ∼5% error rate of Nanopore. A multiple sequence alignment of this region is provided in Fig. S10.

**Table 2.** Lexicon of triplet block order in *T. gondii* and *N. caninum* mtDNA.

| Sequence Block | Arrangement | Tg ME49 | Nc |
| --- | --- | --- | --- |
| A | E-A-T | Y | Y |
|  | E-A-P | Y | Y |
|  | N-A-P | Y | Y# |
|  | N-A-T | Y | Y |
| B | J-B-M | Y | Y |
|  | J-B-Mp | Y | Y |
| C | S-C-Q | Y | Y |
| D | V-D-K | Y | Not detected |
|  | V-D-Kp | Y | Not detected |
|  | Sp-D-K | Not detected | Y |
|  | Sp-D-Kp | Not detected | Y |
| E | J-E-A | Y | Y |
| F | O-F-M | Y | Y |
|  | O-F-R | Y | Y |
| Fp | N-Fp-M | Y | Y |
|  | N-Fp-R | Y | Y |
| H | P-H-Q | Y | Y |
| I | P-I-K | Y | Y |
|  | P-I-Kp | Y | Y |
| J | L-J-B | Y | Y |
|  | O-J-B | Y | Y |
|  | L-J-E | Y | Y |
|  | O-J-E | Y | Y |
|  | L-J-Q | Y | Y |
|  | O-J-Q | Y | Y |
| K | D-K-O | Y | Y |
|  | I-K-O | Y | Y |
| Kp | D-Kp-U | Y | Y |
|  | I-Kp-U | Y | Y |
| L | V-L-J | Y | Y |
| M | B-M-F | Y | Y |
|  | B-M-Fp | Y | Y |
| Mp | B-Mp-U | Y | Y |
| N | Fp-N-A | Y | Y |
| O | J-O-F | Y | Y |
|  | J-O-K | Y | Y |
| P | A-P-H | Y | Y |
|  | A-P-I | Y | Y |
| Q | C-Q-J | Y | Y |
|  | H-Q-J | Y | Y |
| R | F-R-S | Y | Y* |
|  | Fp-R-S | Y | Y* |
| S | V-S-C | Y | Y |
|  | V-S-R | Y | Y |
| Sp | V-Sp-D | Not detected | Y |
| T | A-T-V | Y | Y |
| U | Kp-U-Mp | Y | Y |
| V | L-V-D | Y | Not detected |
|  | T-V-D | Y | Not detected |
|  | L-V-S | Y | Y |
|  | L-V-Sp | Not detected | Y |
|  | T-V-S | Y | Y |
|  | T-V-Sp | Not detected | Y |

#=When block N and A occur together, the first 100 bp of A are missing and instead this junction contains 4 new bp 'CTAT' or 'ATAG'.
\*=When block R and F/Fp occur together, the last 4 bp of F/Fp are missing and instead this junction contains the 2 nt 'AC/GT' depending on orientation. Y = present.

### mtDNA Sequence Blocks are Evolutionarily Conserved in the Toxoplasmatinae but not *Eimeria*

As the mitochondrial genome sequence was also lacking for the closely related *Neospora caninum* we analyzed the available genomic and transcriptomic sequence data for this species. An older assembly of the *N. caninum* genome contained a contig comprising 20 out of the 21 SBs detected in *T. gondii*. The block missing in this contig was identified in the current *N. caninum* assembly (Table 1; Table S2). All 21 mtDNA SBs are highly conserved between the two species including sequence and sequence length (Table 1; Table S2; Dataset S7). However, *N. caninum* does contain an additional partial SB, block Sp, relative to *T. gondii*.

Analysis of *N. caninum* Sanger genomic and EST reads using the 21 *N. caninum* SBs revealed the presence of non-random arrangements similar to those identified in *T. gondii* (Figs. S3-S4). We generated and analyzed low-coverage Oxford Nanopore genome data for *N. caninum* strain Nc-1 and observed single-molecule reads (0.374-15.6 kb) consisting entirely of concatenated mtDNA SBs, some of which are capable of encoding cytochromes and some of which cannot (Fig. S11; Table S3; Dataset S8), as was observed in *T. gondii* ME49 (Fig. S6). *N. caninum* ESTs encoding mtDNA SBs, which if assembled could encode each of the cytochromes were also detected (Table S5, Dataset S9).

Examination of the available sequence data from another closely-related Toxoplasmatinae, *Hammondia* revealed the presence of all 21 mtDNA SBs. It should be noted that the best matches for a number of blocks were to NUMTs and the ends of a few blocks could not be detected likely due to the sparsity of sequence data available for this species (Table 1, Table S2). However, we were not able to conclusively determine the nature of the mtDNA in the more distantly-related *Sarcocystis neurona* a member of the Toxoplasmatinae separated by ∼250 million years (Su et al. 2003). While mining publicly available *Sarcocystis* sequence data we identified hits to a number of the 21 mtDNA SBs but the block boundaries were rarely conserved (Table S2) and we were unable to detect similar permutation patterns. However, this genome sequence still exists in a very large number of contigs and a mitochondrial genome sequence has not been published. An examination of the more distant coccidian *E. tenella*, (Su et al. 2003) (∼500 million years of divergence) for which a published mtDNA genome sequence exists (Hikosaka et al. 2011), did not reveal fragmented blocks as are observed in *T. gondii* (Fig. 1). Examination of the *S. neurona* sequences using the assembled mtDNA and annotated genes of *E. tenella*, does not strongly support an *E. tenella*-like mtDNA either. Together, the findings suggest that the unusual nature of the mtDNA is a derived state shared among the closest relatives of *T. gondii*.

### The Order, Orientation and Number of the 21 Sequence Blocks is Highly Variable but not Random

To determine if any repeating patterns of larger-scale identity could be identified in the *T. gondii* ME49 mtDNA, dot plot analyses of Nanopore reads with a variety of window sizes were performed. At a window size of 100 nt, everything hits everything (data not shown). At a window size of 1,000 nt, the dot plot is still dense (Fig. 6A). At a window size of 2000 nt, lots of hits are seen (Fig. 6B) but at a window size of 3000 nt, nearly all hits vanish (Fig 6C). In other words, no significant stretches of sequence identity, longer than 3 kb, other than the few stretches indicated are observed. The analysis was repeated between *T. gondii* and *N. caninum* Nanopore reads (Fig. 6D). Again, no significant sequence identity larger than 3 kb is observed. Each Oxford Nanopore sequence read >4 kb is unique.

**Fig. 6.**
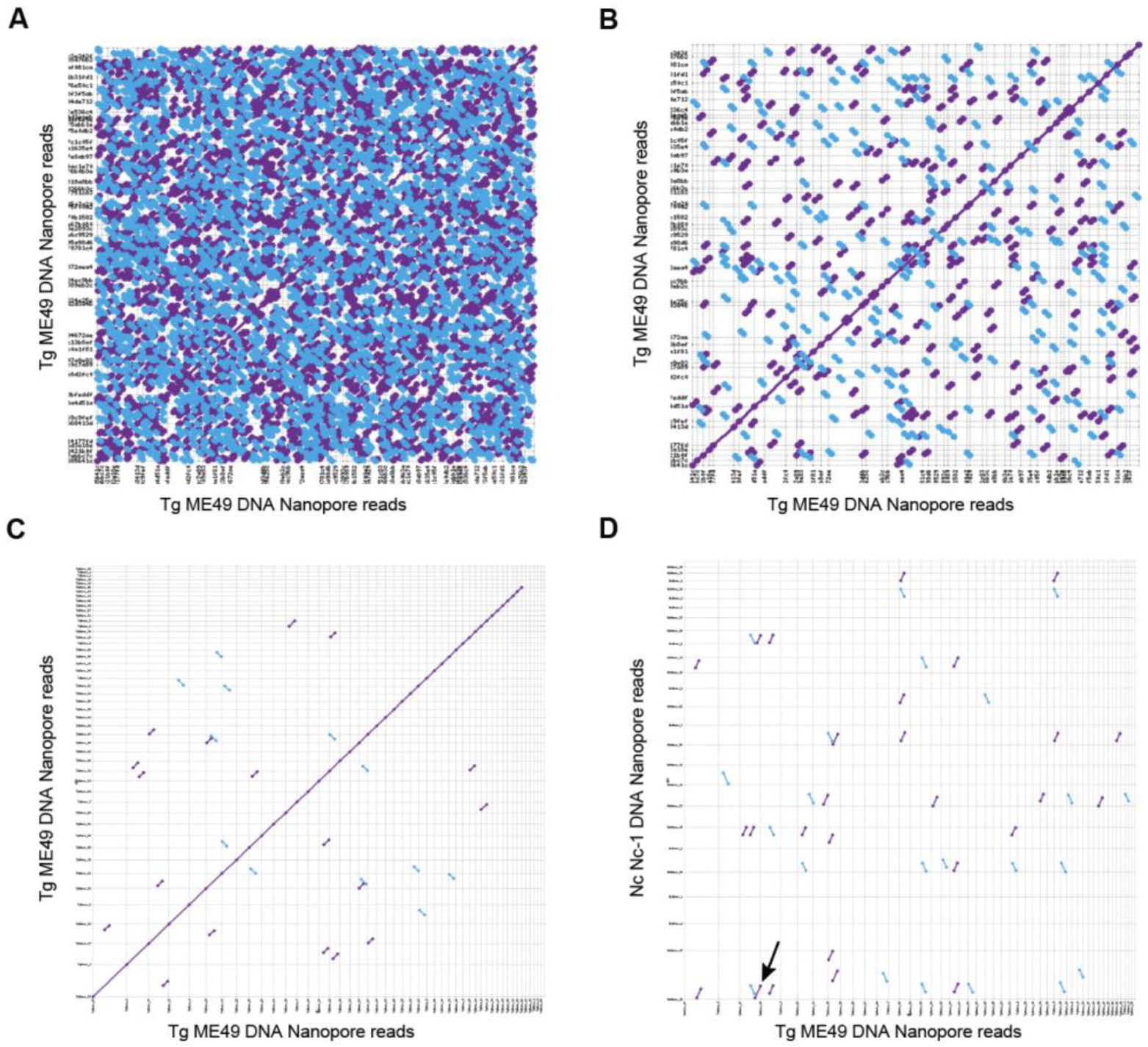
Dot plot comparisons of long *T. gondii* ME49 Nanopore reads against themselves and *N. caninum* Nc-1 Nanopore reads. (A-C) 48 mtDNA-specific *T. gondii* ME49 Nanopore reads (Fig. S6; Dataset S3) were aligned against each other and visualized as a dot plot (D) 48 mtDNA-specific *T. gondii* ME49 Nanopore reads were compared to 25 mtDNA specific *N. caninum* Nc-1 Nanopore reads (Fig. S11; Dataset S8) and visualized as a dot plot. Matches in the forward and reverse orientation are indicated as purple and blue lines respectively. Matches with a window size of 1,000 nt are shown in (A); a window size of 2,000 bp (B and D); a window size of 3,000 nt (C). The arrow in (D) points to the only match observed to be longer than 3000 bp.

To better analyze sequence block orders, a lexicon of all triplet block orders was constructed from the Nanopore mtDNA sequences for both *T. gondii* and *N. caninum* (Table 2). This analysis revealed that the sequence orders are not random and that most SBs are only associated with a few other SBs. The lexicon also revealed that *T. gondii* and *N. caninum* share the majority of their triplets but not all. *T. gondii* ME49 contains the following triplets that *N. caninum* does not: V-D-K, V-D-Kp, L-V-D and T-V-D while *N. caninum* contains Sp-D-K, Sp-D-Kp, V-Sp-D, L-V-Sp and T-V-Sp to the exclusion of *T. gondii* (Table 2; Figs. S6, S11). Comparison of these triplets demonstrates that this difference between *T. gondii* and *N. caninum* is due to the acquired presence of block Sp in *N. caninum* between blocks D and V. Closer examination of the SBs Sp and D in *N. caninum* revealed a 8 bp overlap between the 3’ end of Sp and the 5’ end of D suggesting microhomology as a possible mechanism for the creation of this arrangement. Additionally, *N. caninum* also contains an altered version of SBs N-A and F/Fp-R junctions. Specifically, at the N-A junction, the first 100 bp at the 5’ end of A are missing and instead contains the sequence ‘CTAT’ or ‘ATAG’ depending on the orientation. In the F/Fp and R arrangement, the last 4 bp of F/Fp are replaced by the 2 nucleotides ‘AC’ or ‘GT’. These differences in A and F/Fp are detected only when they occur next to N and R respectively and is observed in all instances when they co-occur.

To visualize the pattern of sequence block relationships a matrix was created in which the identity of the element directly at the 5’ and 3’ end of each sequence block was identified and counted in the Nanopore mtDNA reads in both *T. gondii* ME49 and *N. caninum* (Fig. S12 A-B). Certain SBs are always observed upstream or downstream of specific SBs (*e*.*g*. block L) whereas other blocks (*e*.*g*. block J) are observed to be flanked by several different blocks (Fig S12 A-B; Table 2). The analysis was also performed with *T. gondii* PRU Nanopore direct RNA strand sequencing and *N. caninum* Sanger EST reads to verify the occurrence of the same block relationships in transcripts (Fig. S12 C-D). The patterns observed in the RNA Nanopore reads and ESTs are highly similar for each species except for the differences associated with the acquisition of Sp in *N. caninum*. Thus, the basic units of mtDNA secondary structure are evolutionarily conserved.

### The Topology of the Mitochondrial Genome Remains an Enigma

Oxford Nanopore single-molecule sequencing of *T. gondii* ME49 revealed molecules as long as 23 kb consisting fully of mtDNA SBs although most Nanopore reads were shorter (Figs. 7A, S6). To assess whether or not the Nanopore reads were informative with respect to the size(s) of the naturally occurring mtDNA molecules (a read cannot be longer than the naturally occurring mtDNA molecule), we plotted the size distribution of Oxford Nanopore read from the same run separated into mtDNA reads and nuclear reads (Fig. 7A). As the number of reads was quite different in each of the three Nanopore runs (Table S3) the nuclear reads could not be plotted on the same graph. Nanopore read sizes for *Toxoplasma* mtDNA reads decay more quickly and do not reach the lengths observed in the nuclear reads. The observed mtDNA Nanopore read lengths are similar to the range of sizes observed in a Southern of a CHEF blot probed with a 1,021 bp section of the *cob* gene (Fig. 7B) but the abundance of Nanopore reads in the 1-4kb range to not match the intense smear of (∼5-16 kb) observed on the blot (Fig. 7A-B). However, both views clearly illustrate a wide variation in mtDNA size that ranges from considerably smaller than other observed apicomplexan mtDNA (Fig. 1) to considerably larger. A similar Southern smear was observed in *P. falciparum* (Preiser et al. 1996) where rolling circle replication creates tandem concatemers (Wilson and Williamson 1997). Physically, if the *Toxoplasma* mtDNA exists as a linear concatemer of tandemly repeating units the dot plots (Fig. 6) should have revealed this topology. To test if a tandemly repeating sequence structure is present, restriction digestion with an enzyme that has a single cut site within the unit should result in one major band and a trailing smear of smaller fragments, as is seen in *Plasmodium* (Preiser et al. 1996). However, when *T. gondii* genomic DNA was digested with *XhoI*, which uniquely cuts in the *cob* gene and probed with a *coxI* probe, the probe identified two faint bands of ∼1.35 and 1.65 kb in addition to a large smear (Fig. S13) but a pattern characteristic of a tandem repeat as was not observed. These experiments did not provide any clear indication of the topology of the *T. gondii* mitochondrial genome.

**Fig. 7.**
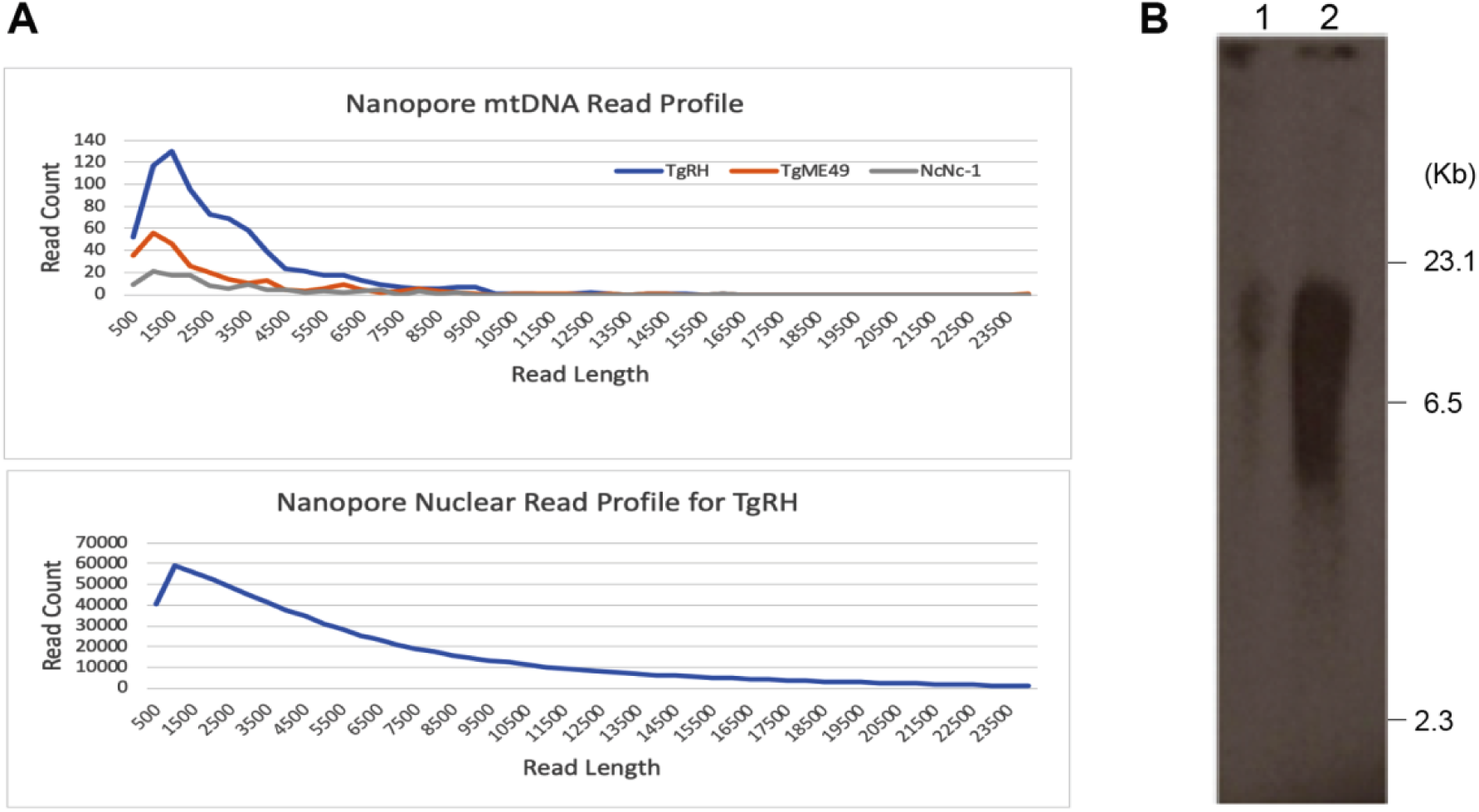
Oxford Nanopore Technology read length distribution and CHEF electrophoresis of DNA from mitochondrial enriched *T. gondii* cell fractions. (A) Profile of read counts plotted by length for mtDNA (top) and nuclear (bottom). Species and strains are as indicated in the legend. (B) Southern of a Contour-clamped homogeneous electric field gel electrophoresis of *T. gondii* total DNA (Lane 1) and mitochondrial-enriched DNA (Lane 2) probed with a 1012 bp section of the *cob* gene. DNA ladder is as indicated. A plot of the ME49 and Nc-1 nuclear reads is located at the bottom of Table S3 since they could not be plotted here due to difference in read count scale.

The secondary structure of the mtDNA appears to be dynamic. There was a time span of several years during which *T. gondii* was being passaged in between the generation of *T. gondii* ME49 DNA for Illumina reads and when DNA was isolated for Oxford Nanopore sequence generation, which came later. We note that when Illumina paired-end reads are mapped to the Nanopore reads of the same strain, passaged in the same laboratory, the mapping is not perfect (Figs. 4, 8). The mapping further degenerates when paired-end Illumina reads from a different *T. gondii* strain, RH, are mapped (Fig. 8) and the degeneration of mtDNA sequence block order is even stronger when *N. caninum* paired-end Illumina sequence reads are mapped even though a lower mapping stringency was allowed (Fig. 8). A similar trend was observed when *N. caninum* and *T. gondii* Illumina reads were mapped to *N. caninum* Nanopore reads (Fig. S13). Given the fact that the SBs (Table 1) and lexicon (Table 2) are highly-conserved, it seems likely that the long mtDNA molecules evolve rapidly and apparently, uniquely.

**Fig. 8.**
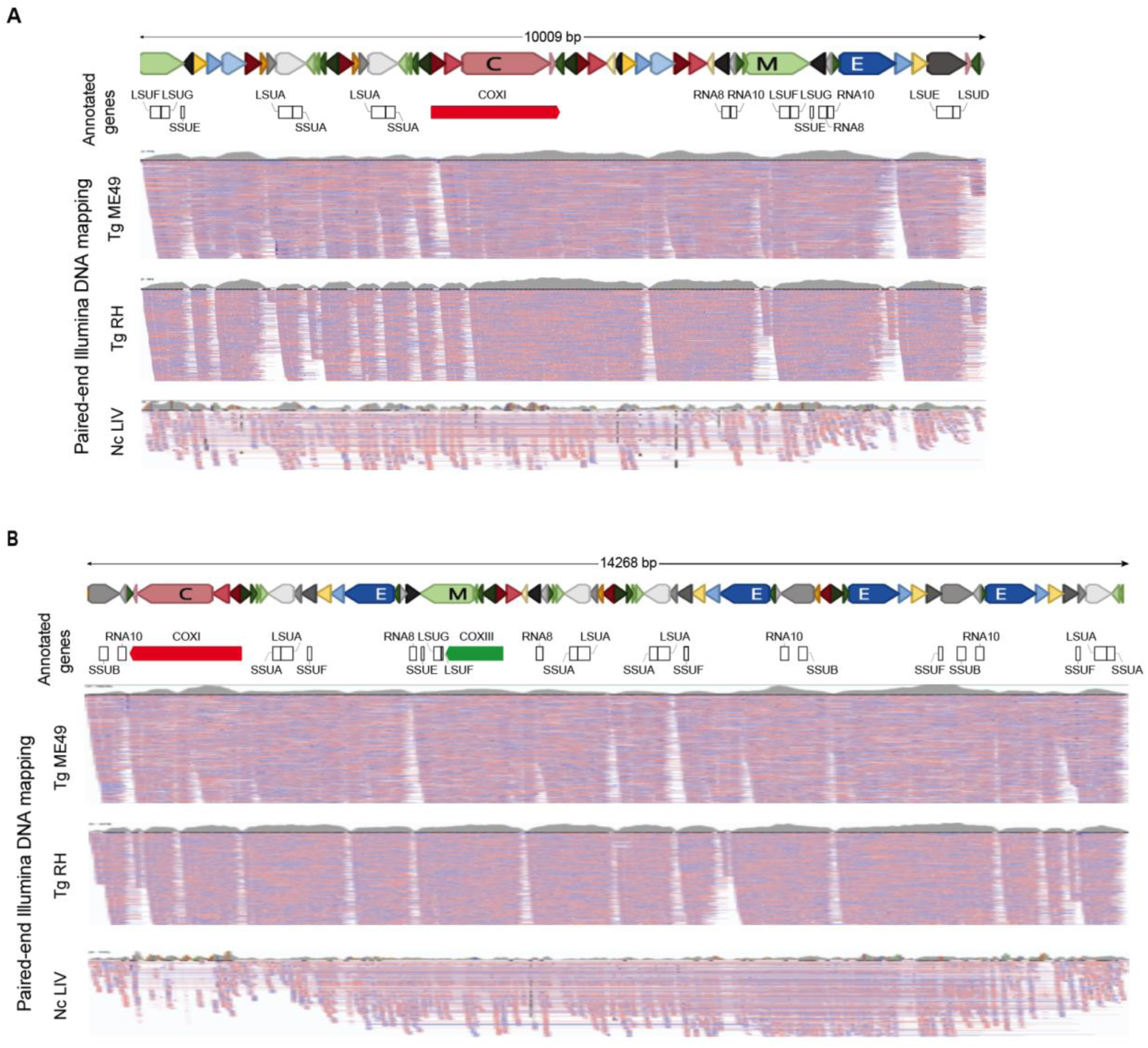
Nanopore and Illumina comparisons reveal sequence block order variability and decay with evolutionary time. (A-B) Two *T. gondii* ME49 Nanopore sequence reads were annotated with the gene sequences they contain. Reads and SBs are drawn to scale. The blocks are colored as shown in Table 1. ‘Annotated genes’ track represents the annotation of the cytochrome genes and rRNA gene fragments on the Nanopore reads. The three ‘Paired-end Illumina DNA mapping’ tracks show paired-end read mapping of *T. gondii* Tg ME49 (SRR9200762), Tg RH (SRR521957) and *N. caninum* (Nc) LIV (ERR012900) mtDNA-specific reads. Mapping of Tg ME49 and Tg RH reads required 100% nucleotide identity whereas 1% mismatch was allowed for mapping Nc LIV reads. Reads were independently mapped to each of the Nanopore mtDNA reads and visualized using IGV. Red and blue lines below each read indicate the mapped Illumina paired-end reads.

The mapped paired-end Illumina reads also reveal the presence of different populations of molecules. There are some paired-end reads that span certain junctions and others that do not as evidenced by the gaps (white space) in paired-end mapping below each Nanopore read (Figs. 4, 8). Likewise, there are regions of Nanopore reads where only half or three quarters of the Illumina reads map to a region as seen in the right-hand edge of Fig. 8A, or the beginning of the TgRH track in Fig. 8B.

The single-molecule Nanopore reads are devoid of obvious telomeres at their ends and while riddled with inverted SBs throughout, they do not appear to be located at the ends of the molecules with any obvious pattern (Smith and Keeling 2013). A few Nanopore reads have portions of the same sequence block on their ends, a feature characteristic of rolling-circle replication (Fig. 9A), but most do not (Fig. 9C). Attempts to circularize the Nanopore reads using the same program that was able to detect and circularize the *P. falciparum* mtDNA, Circlator (Hunt et al. 2015), revealed many potential circular chromosome candidates. The candidates range in size from 1.6-5.6 kb some of which are shown in Figs. 9 and S15, but the putative circular molecules (Fig. 9B, D, S15) do not contain any measurable degree of support from other Nanopore reads (Figs 9A, 9C top panels). The putative circular molecules do have significant support over some regions from mapped paired-end Illumina reads (Figs. 9A, C bottom panels and Fig. S15), but this support is not uniform in many of the putative candidates. A few of the putative circular chromosomes encode full-length cytochrome genes, for example *cob* is encoded by sequence blocks E, A, T (Figs. 9A, B) and the other cytochrome genes are detected (Fig. S15). Finally, to address the possibility that there is a population of very small circular chromosomes, we identified all Nanopore reads 0.6–2.0 kb in length from the 779 mtDNA Nanopore reads identified in the *T. gondii* RH ELQ-316 mutant strain and asked if they could be clustered into significantly overlapping populations. We did not find any support for populations of reads that were likely derived from one or more identical short templates.

**Fig. 9.**
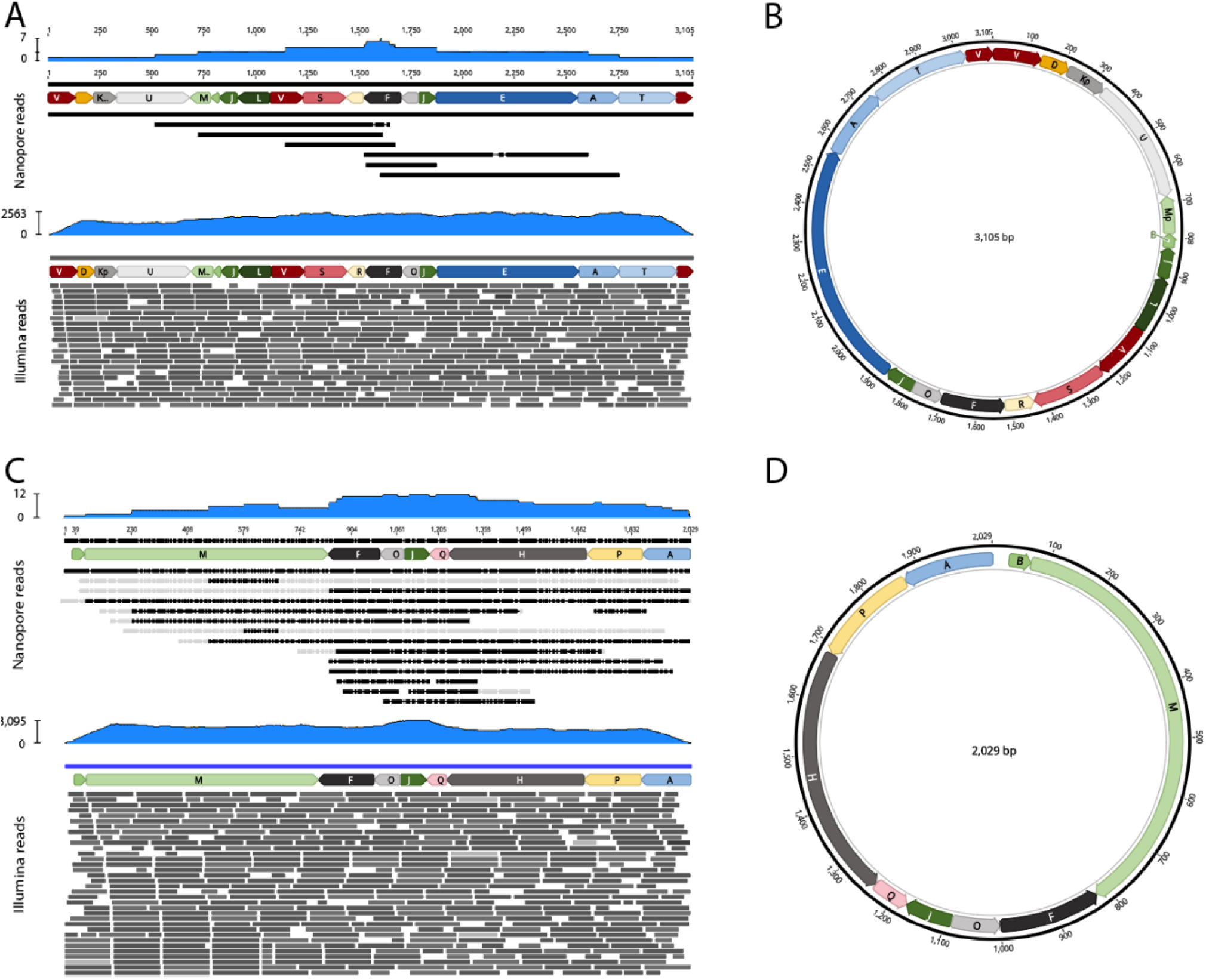
Nanopore and Illumina support for possible circular structures. (A, C) Mapped tracks of TgME49 Nanopore (top panels) and TgME49 Illumina reads (bottom panels) to single Nanopore reads identified as possibly existing as circular molecules according to Circlator predictions (B, C). Note the presence of the *cob* gene, sequence blocks E, A, T in panel B. Paired-end Illumina mapping was not required. Not all mapped reads are shown. The depth of read mapping is indicated above each panel in blue. The portions of Nanopore reads that are greyed out in the top panels do not map to the annotated Nanopore read above. Additional predictions some of which encode the remaining cytochrome genes are provided in Fig. S15.

## Discussion

The mtDNA of the apicomplexan pathogen *Toxoplasma gondii* and its closest relatives is novel and highly fragmented. Minimally, the mtDNA consists of 21 non-redundant sequence blocks named A thru V, none of which individually encodes a full gene, for a total of 5,909 bp of sequence in *T. gondii* and 5,908 in *N. caninum*. The evidence suggests that molecules of mtDNA became shattered and rearranged by some unknown event, prior to the divergence of the tissue coccidia. Single mtDNA molecules of 0.15-23 kb comprised exclusively of SBs, in non-random arrangements, are observed. Single-molecule Nanopore reads longer than 3 kb are unique. Despite the novel genome sequence(s) which include many redundant fragments of protein coding and rRNA genes, full-length *coxI, coxIII* and *cob* genes and their transcripts are detected.

The fragmented nature of the *T. gondii* mtDNA is supported by multiple lines of evidence: Individual unassembled Sanger reads emerging from community genome and EST projects; PCR from DNA isolated from mitochondria enriched cell fractions; Single-molecule Nanopore long read sequences of DNA and RNA and finally, evolutionary conservation of nearly identical sequence blocks in the related tissue coccidia, *Hammondia* and *Neospora*. Despite the high degree of mtDNA fragmentation, Nanopore DNA and RNA reads confirm the presence of some full-length *coxI, coxIII* and *cob* genes with uninterrupted open reading frames. Other expected mtDNA features are also present including 12 of the SSU and LSU rRNA gene fragments and 2 RNA genes observed in other apicomplexans. Additional RNA and protein features were not detected. The coding sequences of *coxI* and *cob* are well conserved with other apicomplexans, but *coxIII* which is reported here for the first time is less well conserved. Low conservation was also reported for the *coxIII* of a free-living ancestor of the Apicomplexa, *Chromera*. The authors report it was nearly overlooked (Obornik and Lukes 2015). *Toxoplasma gondii*, like several species in the alveolata has evolved overlapping coding sequences. Sequence block V is required to form the full open reading frame (using opposite DNA strands), for both *coxI* and *coxIII*, reminiscent of the fused *coxI* and *coxIII* in chromerids (Flegontov et al. 2015; Obornik and Lukes 2015; Gagat et al. 2017), albeit a different fusion.

Evidence for the expression of every single sequence block was seen in the Nanopore direct RNA strand sequence dataset including SBs for which we have no annotation (blocks D, N, P and R). Transcripts capable of encoding full-length *coxI, coxIII* and *cob* were also observed in the Nanopore RNA strand sequence data. Not all transcripts contain full genes. As was observed in the dinoflagellates, partial cytochrome genes are expressed. The mtDNA transcripts are short ≤ 2.8 kb. As was observed in the dinoflagellates, *coxI* does not have an ATG start codon, but *coxIII* and *cob* do. All genes have stop codons. There is no evidence of trans-splicing or RNA-editing. There is no evidence of self-splicing introns, a feature often observed in mtDNA (Lang et al. 2007). Nanopore direct RNA transcripts revealed poly(A) tails on a subset of *coxI* transcripts only. In the dinoflagellates, the cytochrome transcripts are polyadenylated but this occurs upstream of the stop codon (Jackson et al. 2007). Apicomplexans do display polyadenylation of mtDNA transcripts and they have been well characterized in *P. falciparum* where even the rRNA fragments have been shown to be polyadenylated with tails of up to 25 nt (Feagin et al. 2012). Dinoflagellates display strong stem loop and multiple hairpin features (Jackson et al. 2007) in non-coding regions however, the dot plot analyses performed here did not find strong evidence for long, nearly perfect hairpins other than the structure in sequence block J. Examination of *coxI* transcripts that are flanked by sequence block J reveals that sequence block J may encode both a cryptic poly(A) signal and a prominent stem loop structure.

Fragmented mtDNA sequences have been observed in other organisms, most notably the sister phylum to the Apicomplexa, the dinoflagellates, which also share fragmented rRNA genes and a reduced mtDNA gene content of only *coxI, coxIII* & *cob*. What is fascinating from an evolutionary perspective is the fact that the dinoflagellates *Crypthecodinium* and *Karlodinium*, and a basal dinoflagellate, *Hematodinium*, have also fragmented their cytochrome genes and contain both full-length and fragmented copies (Jackson et al. 2007). An examination of several reported dinoflagellate cytochrome gene fragments in *Hematodinium* reveals that the sequences have been shattered in different locations relative to *T. gondii* and that there are a greater number of different, often overlapping fragments in the dinoflagellates. *Toxoplasma* has a minimal set of 21 non-redundant mtDNA fragments and only two fragments, C and H overlap for 50 bp, a region corresponding to the very end of *coxI*. Thus, this phenomenon has arisen independently in both the dinoflagellates and the Apicomplexa.

Concatenated mtDNA is not unusual as an observed mitochondrial structure. It is a frequent observation in the Apicomplexa and numerous other organisms including yeast and even humans under certain circumstances (Ling and Shibata 2004; Bedoya et al. 2009). However, *Toxoplasma* concatemers are different from previously described mtDNAs because molecules >3 kb appear to be unique. This observation contrasts with concatemers of a single repetitive sequence as is observed in rolling-circle replication. Theoretically, the mtDNA of *Toxoplasma* and its relatives could be either circular, linear or, some combination of both. Circular chromosomes that yield linear concatemers and linear chromosomes are observed in the Apicomplexa. Likewise, all mtDNA could be contained on a single chromosome as is observed in all Apicomplexa to date (Fig. 1), or distributed onto several distinct chromosomes, either circular or linear as has been observed in many organisms (Burger et al. 2003; Flegontov and Lukes 2012; Dong et al. 2014; Yahalomi et al. 2017). The fact that *Toxoplasma* long mtDNA sequences are often unique, but also non-random, is quite informative and sheds light on the various possibilities.

The data presented here do not support a single circular topology that replicates via rolling-circle replication as the observed concatemers are not constructed of a single repeating sequence unit as evidenced by read annotation, the dot plot comparisons or Southern blot analysis of restricted DNA. The formal possibility exists that each of the 21 SBs is contained on its own chromosome and that copies of the chromosomes become concatenated, non-randomly via some undiscovered process that guides particular ligations. This too is unlikely because if the SBs existed as linear chromosomes, they would need inverted repeat at their ends for replication and if they existed as circular chromosomes they would replicate via rolling-circle replication and we should detect concatemers of single SBs. We do not see them, even at very deep sequencing coverage. We do however detect support for the hypothesis that when the mtDNA shattered into smaller pieces, it shattered redundantly into many different units of sequence, each of which is capable of being annotated with more than one of the 21 SBs. While it is impossible to estimate an average mtDNA genome size, there are clearly more mtDNA mitochondrial reads relative to the nuclear genome equivalent than there are nuclear reads. Estimated copy numbers for full-length cytochrome genes relative to a single copy nuclear gene are several hundred copies to 1 based on Illumina read mapping.

We see that each of the cytochrome genes can exist as 3-5 fragments. We also observe the presence of one or more full-length cytochrome sequences on some of the Nanopore reads in addition to the cytochrome gene fragments. The data suggest that both whole genes and fragments exist, simultaneously. Certain short arrangements of SBs totaling 1-3 kb of sequence are observed to occur multiple times within the Nanopore reads (Fig. 6). The lexicon of sequence block orders (Table 2) reveals a restrictive vocabulary of triplets. Indeed, some SBs, *e*.*g*. blocks C and H occur in only two contexts, S, C, Q, and P, H, Q. Both of these longer annotated sequence stretches involve sequence block Q, so Q must exist in at least two distinct molecules. The differences between *T. gondii* and *N. caninum* are also supportive of longer stretches of sequence. When a sequence block is altered as in the creation of Sp or when SBs N and A occur next to each other in *N. caninum*, each occurrence is always identical, *i*.*e*. V, Sp, D or N, A when together but not when each sequence block is found in other arrangements (Table 2). Thus, these stretches of sequence are likely physically encoded together as a single larger unit. The lexicon, the dot plots and the analyses of upstream and downstream SBs in genomic and transcriptomic data suggest that units as small as 3 SBs are quite well conserved. They data support a model in which parts of the mitochondrial genome, including at least one full-length copy of each cytochrome gene are redundantly present on different chromosomes. It is much less risky to the organism if it maintains at least one full-length copy of each gene or rRNA gene fragment.

The situation is more complex when considering stretches of DNA longer than the triplets in the lexicon. The dot plot analysis confirms that sequences longer than 3 kb are unique but it also reveals that longer sequences share numerous stretches of sequence 1-3 kb in length. It is probable and parsimonious to hypothesize that larger sequences are constructed, albeit uniquely, from smaller sequences. The likely mechanism is homologous recombination. If the mtDNA of *T. gondii* exists as a population of circular molecules, each of which contains multiple portions of the ancestral mtDNA redundantly, then homologous recombination between two different circular chromosomes sharing a homologous region (or stretch of microhomology) will create a novel larger molecule. With 21 different SBs identified, recombination can happen at many different places between however many different circular chromosomes contain a homologous sequence. Recombination among such a population would give rise to the variety of sequences that are detected by Nanopore. It would explain some of the errors we see in some of the Nanopore reads (Tables S6-7) where we see rearranged portions of a sequence block. We have highlighted these regions in yellow and annotated with a repeat of the SB, *e*.*g*. KK, CC, FF or MM but there are others. Recombination has been observed in *P. falciparum* during mtDNA replication (Preiser et al. 1996) and it is a widely reported phenomenon within and between mitochondria across eukaryotes (Sandhu et al. 2007; Chen 2013; Gualberto and Newton 2017).

In Figs. 4, 8 and S14 we mapped paired-end Illumina reads from the same strain, same species or related species to Nanopore reads to assess the strength with which each Nanopore read is supported. The DNA for the Nanopore and Illumina sequences in the same species were generated years apart and the effects of time are evident. The Nanopore reads have good support for many regions, but rarely strong support for the entire molecule and this support decays with evolutionary distance as evidenced by the increasing white space and dips in the grey histograms that indicate areas where reads don’t map. We required both ends of a paired read to map precisely to assess contiguity of the reference sequence. This mapping experiment also revealed the presence of sub-populations of Illumina reads that provide partial support for some regions. These observations are consistent with the above hypothesis that the larger molecules sequenced by Nanopore resulted from unique recombination between two or more smaller molecules, likely but not proven, to be circular. Since recombination is likely occurring, is variation present within a single mitochondrion? The data presented here cannot address that question as DNA was extracted from millions of parasites and as of yet, individual mitochondrial organelle sequencing is not yet possible. However, the *T. gondii* RH ENU mutant strain that we sequenced with Nanopore is quite informative. DNA from the *T. gondii* ELQ-316 mutant was from a recently-derived clone presumably originating from a single resistant parasite. The Nanopore reads from this mutant also yielded a myriad of long, unique, sequence block concatemers, suggesting that they either already existed in the parent mitochondrion or are easily regenerated. The ENU-induced point mutation in this clone exists in sequence block E of the *cob* gene. When we analyzed the uncorrected Nanopore reads that contained the mutated region of sequence block E. We observed that the point mutation was present in nearly all copies of sequence block E. This finding suggests that *T. gondii* mtDNA either uses a template copying mechanism or has the machinery needed to maintain identity, *e*.*g*. homoplasmy between sequence blocks if they physically occur on different chromosomes, likely including a DNA mismatch repair protein MSH1, a MutS homolog that targets to the mitochondrion in *Toxoplasma* (Garbuz and Arrizabalaga 2017). This finding is exciting for those planning experiments targeting the mtDNA. The mitochondrion is an important drug target and although the mtDNA appears to exist as a population of molecules with high complexity, the prospect exists for engineered alterations that can propagate such as this point mutation.

The size of the mtDNA has been minimally determined at 5909 bp based on the SBs. In *P. falciparum* a Southern Blot smear ranging from (6-23 kb) was attributed to rolling-circle replication intermediates (Weissig and Rowe 1999). Smears of 6-10 kb or more were observed in dinoflagellates (Jackson et al. 2007; Nash et al. 2007; Flegontov and Lukes 2012). *T. gondii* showed a smear ranging from 2-16 kb with greatest signal above 5 kb (Fig. 7B). This size range overlaps with the sizes of the Nanopore reads detected but Nanopore detected more molecules less than 5 kb, perhaps due to DNA breakage.

Examination of sequence data from the closely-related (∼12 my divergence) parasites, *N. caninum* (Reid et al. 2012) and *H. hammondi* (Walzer et al. 2013) indicated that these parasites also share fragmented mtDNA as is observed in *T. gondii*. This characteristic provides another synapomorphy to link these taxa to the exclusion of *Sarcocystis* (Tenter et al. 2002). The *Sarcocystis* mtDNA remains uncharacterized but based on our searches of the limited available data, its structure differs from both *Eimeria* and *Toxoplasma*. Importantly, all mtDNA sequences generated to date reveal rearrangements of gene order and orientation (Fig. 1). Clearly, the capacity to both generate and survive structural rearrangements within such a small genome is a prominent feature in both the Apicomplexa and dinoflagellates.

Regulation of mtDNA gene expression in this highly polymorphic mtDNA context is unknown. Based on the data available we have no clear promotor candidates. All SBs are expressed. *coxI* transcripts seem to begin at sequence block V, the beginning of the coding sequence, but *coxIII* and *cob* are found in transcripts that have additional SBs located upstream of the coding sequences.

Elucidation of the mtDNA of *T. gondii* might have happened years ago were it not for NUMTs The presence of NUMTs in the *T. gondii* nuclear genome (initially referred to as ‘REP’ elements) was first reported by Ossorio *et al*., (Ossorio et al. 1991). A probe made from the region of sequence downstream of the single-copy nuclear gene, *rop1*, hybridized to multiple regions of the nuclear genome resulting in smeared signals in Southern hybridization assays. This region of their probe contained NUMTs of the, *coxI* and *cob* genes that we can now annotate as mtDNA SBs typically found in apicomplexan mtDNA (Table S7). *Toxoplasma gondii* contains >9000 NUMTs in its nuclear genome (Namasivayam 2015) thwarting molecular and sequence assembly based approaches. The NUMTs together with the unique architecture presented here generated a difficult puzzle that required new technology for a breakthrough. Single-molecule Nanopore sequencing of both DNA and RNA proved to be essential tools. In the absence of this data, the individual Sanger sequences and hybridization data pointed in the direction revealed by Nanopore but they could not determine and validate the findings presented here. If we had begun this study with only Nanopore reads and then annotated mtDNA features on these molecules, the 21 SBs that are used to architect these long molecules would not have been evident and likely would have been missed. Single-molecule long-read technology should be equally useful for understanding the dinoflagellate mtDNA. The presence of mtDNA SBs and the fact that their non-random concatenation creates unique molecules is fascinating biology.

### Conclusion

Evolutionarily, the mtDNA of *T. gondii* seems more similar to that of the dinoflagellates than apicomplexans, yet their fragmented mtDNA evolved independently. Our inability to assemble a single full-length mitochondrial genome sequence and the presence of sequence block order and orientation variants suggests that the *T. gondii* mtDNA could exist as number of distinct molecules as is seen in dinoflagellates. We find a similar fragmented mtDNA pattern in *N. caninum* proving that this is a conserved phenomenon. Despite the long unique concatemers of SBs that are observed, full-length *coxI, coxIII* and *cob* genes and transcripts are detected. We still do not know the exact topology of the mitochondrial genome or how it replicates and segregates during daughter cell formation. While the lack of a single mtDNA sequence raises difficulties for those targeting the *T. gondii* mitochondrion, the presence of a mechanism for maintaining homoplasmy at the level of a sequence block was observed. This finding opens the door to further exploration of the *Toxoplasma* mitochondrion. The need to be able to purify mitochondria in *Toxoplasma* remains strong. Nanopore technology offers the opportunity for re-analysis of the dinoflagellate mtDNA and further exploration of mitochondrial biology.

## Supporting information

Data-Set-Files

## Materials and Methods

### Species and Strains Utilized in this Study

*Toxoplasma gondii* RH and ME49 (kindly provided by David Sibley) tachyzoites were maintained in confluent monolayers of human fibroblast reverse transcriptase (hTERT) cells in DMEM supplemented with 1% heat inactivated fetal bovine serum, 2 mM glutamine and 1% Penicillin/Streptomycin (Weiss and Kim 2007). 2X PBS washed extracellular tachyzoite stages were used for mitochondrial enrichment or DNA extraction. *Neospora caninum* Nc-1 (Dubey et al 1988) were cultured on Vero cells using RPMI 1640 supplemented with 5 % normal horse serum, purified by centrifugation through 30 % isotonic Percoll and washed twice in PBS. Pellets of purified tachyzoites were stored at -20°C.

### Enriched Mitochondrial Fractions

*Toxoplasma gondii* RH tachyzoite parasites were propagated in hTERT cells as described previously (Weiss and Kim 2007). Parasites (approximately 2-4 × 10_9_) were purified by passing them through a cotton mesh followed by filtration through 8 and 5 micron nucleopore membranes as published (Miranda et al, 2010). The purified suspension was centrifuged and the pellet resuspended in lysis buffer followed by centrifugation at 700 g. The wet pellet was mixed with silicon carbide and this mix was transferred to a mortar and manually ground for 3-4 15 sec periods. The cell-silicon carbide mix was resuspended in approximately 20 ml of lysis buffer, decanted and clarified by three centrifugations at 50 X g (5 min) and 150 X g (twice) (10 min each). The supernatant was centrifuged at 1000 X g or 1200 X g for 10 min to collect the mitochondrial-enriched pellet.

### Nucleic acid Extraction, PCR, RT-PCR and Sequencing

DNA was isolated through standard proteinase K and RNase A digestion followed by phenol/chloroform extraction. Primer pairs were designed utilizing publicly available cDNA sequences of expressed mitochondrial genes and based on the sequences of these PCR amplicons (Table S1). All primers have a T_m_ higher than 57°C. The high-fidelity Platinum Taq polymerase (Invitrogen, Waltham MA, USA) was used for PCR. PCR was performed as follows: initial denaturation at 95°C for 3 minutes; followed by 35 cycles of amplification at 95°C for 60 seconds, 54°C for 45 seconds, and 72°C for 60 seconds. The PCR products were visualized using agarose gel electrophoresis. PCR products, if not pure, were gel extracted prior to sequencing using a PCR purification kit (Qiagen, Hilden, Germany). If the yield of the gel extraction was low, extracted DNA was used as template for a second PCR amplification prior to sequencing.

RNA was extracted from *T. gondii* RH Δ *ku80* using RNeasy Mini Kit (Qiagen) following manufacturer’s Quick-Start protocol with the DNase digestion step. RT-PCR was performed using the Superscript III first-strand synthesis kit (Invitrogen) and cytochrome gene-specific primers (Table S1, *coxI* V2, *coxIII* M9, *cob* E5) to identify the 3 cytochrome transcripts. PCR products of the cDNA were sequenced using primers C9, M9 and T3 (Table S1) to verify the transcript sequences for *coxI, coxIII* and *cob* respectively. DNA for Illumina (San Diego, CA, USA) and Oxford Nanopore (Oxford, United Kingdom) sequencing was extracted from *T. gondii* ME49 parasites using DNeasy Blood and Tissue Kit (Qiagen) following manufacturer’s cultured cells protocol. DNA from frozen *N. caninum* cells was purified using DNeasy Blood and Tissue Kit (Qiagen) according to the manufacturer’s protocol. The DNA was eluted from the spin columns using 50 µl Buffer AE supplied with the kit. The DNA was concentrated to approximately half volume and a final concentration of 20 µg/ml using speedvac. Concentration was measured on Qubit.

Illumina sequencing was performed using the MiSeq platform (PE 150) at the Georgia Genomics and Bioinformatics Core. Long read sequencing on the MinION platform was performed using Rapid Sequencing Kit, SQK-RAD004, and 400 ng of *T. gondii* ME49, *T. gondii* strain RH Δ*uprt* ELQ-316 mutant (DNA kindly provided by J Stone Doggett) or *N. caninum* Nc-1 genomic DNA on a Nanopore flowcell 9.4.1, all according to the standard procedure described by the manufacturer. Data for each sequencing run are provided in Table S3 and deposited in the GenBank SRA as detailed. Base calling was performed directly in the Oxford Nanopore MinKNOW program, version 18.03.1. Nanopore Direct RNA sequence data generated from Poly-A+ purified RNA was provided by Stuart Ralph. All data generated as a part of this project have been deposited in the NCBI GenBank or provided in supplementary data set files.

### DNA Electrophoresis and Southern Analysis

Restriction digestion of *T. gondii* RH genomic DNA was performed with XhoI (New England BioLabs, Ipswich, MA) at 37°C and run on a 0.8% agarose (Sigma, St. Louis, Mo) gel with a 1 kb ladder (NEB, Ipswich MA) and positive control. The gel was then soaked in 0.25M HCl for 15min to depurinate and then denatured with 0.5M NaOH, 1M NaCl for 30min and neutralized with 1.5M Tris HCL, 3MNaCl, pH 7.4, twice, 15 min each, followed by over-night transfer to a nylon membrane and UV fixation. Pre- and post-staining with ethidium bromide was used to examine successful transfer of DNA to the membrane. Blots were pre-hybridized in 30 ml hybridization solution (50% deionized formamide, 5X Denhardt’s solution, 5X SSC, 0.1% SDS, 0.1 mg/ml boiled salmon sperm DNA). A radioactive *coxI* probe (367 bp sequence block C) was boiled for 5min and immediately chilled on ice prior to hybridization at 43°C or 58°C. The membranes were washed several times, each time for 15 min with increasing stringency and finally in 0.1X SSC, 0.1% SDS at 65°C.

### Contour-clamped Homogeneous Electric Field (CHEF) Electrophoresis and Southern Analysis

*T. gondii* RH tachyzoite parasites were isolated and suspended in 1.0% low melting agarose plugs and digested with 0.5M EDTA, 1% lauroylsarcosine, 2mg/ml proteinase K and 0.2mg/ml RNaseA at pH 8.0, 50°C for 48 hours. Electrophoresis was performed using a CHEF Mapper (Bio-Rad Laboratories, Hercules, CA) in 1% agarose gel (Sigma, St. Louis, MO, USA) with the following conditions: 6V/cm, linear 11s-22s, 15hours and 4°C with a switching angle of 120°. A radioactive *cob* probe 1012 bp in length was synthesized through PCR with an annealing temperature of 53°C, using gel-extracted DNA as template and Taq polymerase (New England BioLabs). The probe product was cleaned using mini Quick spin columns (Roche, Basel, Switzerland). Radioactive alpha-32P dATP (specific activity 3000 Ci/mmol) was used. Southern analysis was performed as described above.

### Initial Sequence Analysis and Data Mining Strategies

The sequenced PCR amplicons generated from *T. gondii* mitochondrial-enriched DNA fractions, publicly available unassembled Sanger mtDNA-like EST and genome sequence reads obtained from NCBI EST and Short Reads Archive (SRA) databases were compared with each other using BLASTN (E-value: e_-10_, nucleotide identity ≥98%). 21 mitochondrial SBs were determined based on their reproducible occurrence, sequence boundaries and arrangements in the sequenced PCR fragments as well as publicly available Sanger reads. *T. gondii* genomic sequences (including the unassembled contigs) were obtained from www.toxodb.org release version 11 (Gajria et al. 2008). These contigs and all unassembled *T. gondii* Sanger genomic and EST reads from NCBI were screened via BLASTN (E-value: 1e_-10_, nucleotide identity ≥ 98%) for any additional mtDNA sequences using the determined mtDNA SBs as queries. A contig/read was classified as mitochondrial if no chromosomal sequence was found. All reads and contigs classified as mitochondrial were analyzed for length, sequence boundaries and arrangements using BLASTN.

### Identification of NUMTs

RepeatMasker version 4.0.5 ((http://www.repeatmasker.org) with a repeat library composed of the 21 *T. gondii* DNA SBs and search engine ‘crossmatch’ was used to identify nuclear sequences of mitochondrial origin (NUMTs) in the nuclear genome sequences of *T. gondii* and *N. caninum* based on known *T. gondii* NUMTs (Namasivayam 2015). This step was performed to facilitate distinguishing nuclear-encoded mtDNA fragments from the true organellar mtDNA.

### Analysis of Oxford Nanopore Reads

To obtain possible mitochondrial reads from the Nanopore datasets two strategies were used. First, the Nanopore reads were screened for the 21 mtDNA SBs using BLASTN. Reads with an E-value 1e_-10_, nucleotide identity ≥ 60% and ≤ 10% alignment length mismatch were classified as possibly mitochondrial. Second, the Nanopore reads were aligned against the 21 mtDNA SBs using Exonerate v2.4.0 (Slater and Birney 2005) and the best 2000 alignments and reads with an alignment score ≥ 60% were classified as possibly mitochondrial. To remove Nanopore reads that are nuclear in origin but contain NUMTs, the potential mitochondrial reads identified above were aligned using BWA v0.7.17 (Li and Durbin 2009) against *T. gondii* ME49 chromosomal sequences masked for NUMTs. All reads that did not align to chromosomal sequences were considered putative mitochondrial reads. Finally, error-correction of the mitochondrial Nanopore reads was performed using proovread (Hackl et al. 2014) and *T. gondii* ME49 Illumina reads that mapped to the 21 mtDNA SBs (see below). These error-corrected reads (Dataset S6) were annotated with mtDNA SBs using BLASTN as previously described. All unannotated sequence bits in each Nanopore read, which typically consisted of approximately 50-100 bp at the beginning of a read were examined and compared to each other for any repeating patterns to identify any additional mtDNA sequence. However, no such potential sequence was identified. To check if any of the Nanopore mtDNA reads circularize, Circlator v1.5.3 with minimus2 (Hunt et al. 2015), previously used to identify different circular genomes, including *P. falciparum* 3D7 mitochondrial genome, was used.

### Analysis of Genomic Illumina Reads

Paired-end Illumina reads generated from *T. gondii* ME49 DNA were mapped, requiring 100% identity to the 21 mtDNA SBs using BWA v0.7.17 to identify mtDNA-containing reads. These reads were then further examined to detect any previously unidentified mtDNA sequence that might be present. In order to assess if the Illumina dataset supported the Nanopore read sequences *T. gondii* ME49 (Accession ID: SRR9200762, this study) as well as *T. gondii* RH-88 (Accession ID: SRR521957) mtDNA reads were independently mapped to a few of the corrected *T. gondii* ME49 Nanopore reads described above using BWA v0.7.17 (parameters: local alignment, require both pairs to match at 100% identity) and visualized using Integrated Genomics Viewer (IGV) (Robinson et al. 2011) (Figs. 4, 8 and S14). If different parameters were used for mapping of the Illumina reads, they are indicated in the associated figure legend. When estimating relative copy numbers for mtDNA and single-copy nuclear genes (Table S4), negative controls included mapping to manually created SB fusions of block orders not observed in our lexicon.

### Gene Prediction and Annotation

Mitochondrial genes and rRNA gene fragments were annotated based on similarity to the cytochrome genes and rRNA fragments of *P. falciparum* and *E. tenella* (Feagin 1992; Feagin et al. 1997; Hikosaka et al. 2011; Feagin et al. 2012). TBLASTN and ORF prediction (utilizing the NCBI ORF finder and the genetic code for protozoan mitochondria) was used to identify and annotate the protein coding regions in the sequenced DNA blocks. Open reading frames >50 amino acids from the SBs and the longest *T. gondii* Nanopore read were searched with InterProScan V5.31-70.0 to identify any additional protein features. The identity of the predicted cytochrome sequences was confirmed via comparison to partial or full-length cDNA sequences and full-length genes detected in the Nanopore reads reported in this study, alignments to publicly available incomplete *T. gondii* mitochondrial mRNA sequences and EST reads and through examination of the alignment of the predicted sequences (nucleotide and amino acid) to other apicomplexan mitochondrial proteins using MUSCLE (Edgar 2004)(Fig. S7). Paired-end *T. gondii* RNA-seq reads obtained from NCBI (Accession ID: SRR6493545) were mapped to mtDNA SBs as described above to identify mtDNA specific reads. These reads were then mapped to the predicted cytochrome gene sequences using BWA v0.7.17 (parameters: local alignment, require both pairs to map at 100% identity). *T. gondii* PRU Δ*ku80* Nanopore direct RNA strand reads (Accession ID: SRR9200760) (Dataset S7) were processed similar to the genomic Nanopore reads and annotated to identify the presence of full-length cytochrome sequences. rRNA genes that did not align well with the rRNA gene fragments of *P. falciparum* were identified using conserved nucleotides, manually folded and compared with secondary structures of the *E. coli* rRNA and their *Plasmodium* counterparts. All SBs and the longest Nanopore read were used to search RFAM v14.2 to identify any additional RNA genes or RNA features like catalytic RNAs or self-splicing introns that may be present.

### Mitochondrial Genome Assembly Approaches

MtDNA reads were identified in all datasets as described above. Efforts to assemble the various sequences are detailed as follows: PCR amplicons and unassembled mtDNA Sanger reads were assembled using CAP3 and default parameters since read coverage was low (Huang and Madan 1999). MtDNA PE Illumina reads were assembled with SPADES 3.12.0 (Bankevich et al. 2012) which automatically tests various k-mers sizes and selects the best for the assembly. Corrected mtDNA Nanopore reads were assembled using several assembly programs, such as CANU v1.9 (Koren et al. 2017), Flye v2.6 (Kolmogorov et al. 2019), CAP3 and Geneious Prime 2019.1.3 (Kearse et al. 2012). All efforts were unsuccessful.

### Identification of mtDNA Sequences in *N. canium, Hammondia and S. neurona*

*N. caninum* genomic sequence was obtained from ftp://ftp.sanger.ac.uk/pub/pathogens/Neospora/caninum/NEOS.contigs.072303. A single contig containing mtDNA sequences was identified using the 21 *T. gondii* mtDNA SBs via BLASTN (E-value: e_-10_). This contig was not present in later genome assemblies. *N. caninum* mtDNA SBs were annotated from this contig and the SBs were used to identify additional mtDNA sequence contigs. *N. caninum* Nanopore data was queried similar to *T. gondii* Nanopore reads to identify mtDNA reads and these reads were error-corrected using *N. caninum* paired-end Illumina genomic reads (Accession ID: ERR012900). The error-corrected mtDNA Nanopore reads were annotated using the *N. caninum* SBs and the SBs were further curated based on this annotation (Dataset S8). The mitochondrial genes of *N. caninum* were annotated using *T. gondii* mtDNA genes as well as genomic and expression data. Similarly, the genome sequence of *Hammondia hammondi* H.H.34 (toxodb.org version 2014-06-03) as well as sequences of other *Hammondia* species available in NCBI were mined for mtDNA sequences via BLASTN. *Sarcocystis neurona* SN3 and SN1 genome sequences (versions 2015-04-13 and 2015-07-23 respectively) obtained from toxodb.org and sequence data in NCBI were mined for mtDNA using *T*.*gondii* mtDNA SBs and the mitochondrial genome sequence of *E. tenella* (AB564272.1) using BLASTN and TBLASTX searches (to search for protein coding sequences).

### Other Bioinformatic Analyses and Data Visualization

Various arrangements and permutations of SBs in the different datasets were parsed from BLASTN results using custom scripts and manual inspection. An GFF file containing the annotation of genes and SBs for each read and PCR amplicon was generated using BLASTN (E-value: 1e-10, nucleotide identity ≥ 98%) and a Perl script (bp_search2gff.pl) available from APPRIS (Rodriguez et al. 2018) and visualized using Geneious Prime v.2019 (Kearse et al. 2012). Homology and micro-homology analyses of the SBs and corrected Nanopore reads were performed using nucmer and plotted using mummerplot (Kurtz et al. 2004). The GenBank accession numbers are: *P. falciparum* (AAC63390.2, AAC63389.2, AAC63391.1) and *E. tenella* (BAJ25753.1, BAJ25754.1, BAJ25752.1) and their associated CDSs. Alignment of Illumina reads to SBs, Nanopore and cytochrome sequences was visualized using IGV and coverage was determined using BEDTools 2.21.0 genomeCoverageBed (Quinlan and Hall 2010). Specific parameters used for various analyses if they differ from those listed here, are indicated in associated figure legends and tables.

## Acknowledgements

We would like to acknowledge Silvia Moreno, University of Georgia for providing mitochondrial-enriched fractions of *T. gondii*; Stuart Ralph for sharing his *T. gondii* PRU Δ*ku80* direct RNA strand sequence data; David Sibley for providing *T. gondii* ME49 strain used in the genome project and Tobias Lilja and Faruk Dube for assistance with Oxford Nanopore sequencing. Diego Huet and Charles Delwiche provided useful discussion and Ujwal Bagal and Jeremy DeBarry provided critical feedback on the manuscript. This work benefited from helpful discussions with members of the Kissinger research group both past and present. This work was supported in part through NIH funding (NIH R01 AI068908). This study was supported in part by resources and technical expertise from the Georgia Advanced Computing Resource Center, a partnership between the University of Georgia’s Office of the Vice President for Research and Office of the Vice President for Information Technology.

## Author contributions

SN and JCK designed research; SN, JCK, WX and EMH performed research; KT and JSD contributed new reagents / analytic tools; SN, RPB and JCK analyzed data; SN, RPB, KT and JCK wrote the paper.

## Data deposition

Data deposition: All sequences reported in this paper have been deposited in the National Center for Biotechnology Information under accession numbers:

*T. gondii* ME49 PE-150 Illumina, SRA SRR6793863;

*T. gondii* ME49 Oxford Nanopore mtDNA reads, SRA SRR9200762;

*T. gondii* strain RH Δ*uprt* ELQ-316 mutant Oxford Nanopore whole genome reads SRR9961591;

*N. caninum* Nc-1 Oxford Nanopore mtDNA reads, SRA SRR9200761;

*T. gondii* PRU Δ*ku80* Oxford Nanopore mtRNA reads, SRA SRR9200760;

Tg and Nc *coxI, coxIII* and *cob* nucleotide sequences MN077082-MN077087

*T. gondii* 21 full and 3 partial sequence blocks, MN077088-MN077111;

*N. caninum* 21 full and 3 partial sequence blocks, MN077112-MN077136;

*T. gondii* cDNA *for coxIII* MN267020.

## Supplementary Information

Supplementary figures S1-S15

Supplementary tables S1-S8

Dataset_S1 *Toxoplasma gondii* RH PCR products

Dataset_S2 *Toxoplasma gondii* 21 mtDNA sequence blocks

Dataset_S3 *Toxoplasma gondii* ME49 corrected Nanopore mtDNA reads

Dataset_S4 *Toxoplasma gondii coxI, coxIII* & *cob* nucleotide and amino acid sequences

*Dataset_S5 Toxoplasma gondii RH RTPCR prod*ucts

Dataset_S6 *Toxoplasma gondii* PRU Δ*ku80* corrected Nanopore mtRNA reads

Dataset_S7 *Neospora caninum* 21 mtDNA sequence blocks

Dataset_S8 *Neospora caninum* Nc-1 corrected Nanopore mtDNA reads

Dataset_S9 *Neospora caninum coxI, coxIII* & *cob* nucleotide and amino acid sequences

## Competing interests

None.

## Notes

### Competing Interest Statement

The authors have declared no competing interest.

