## Supplemental Figures for "A novel fragmented mitochondrial genome in the protist pathogen *Toxoplasma gondii* and related tissue coccidia"

ORCID: 0000-0002-6413-1101

**This PDF file includes:**

**Supplementary figures S1-S15**

**Supplementary tables S1- S8**

**Supplementary datasets S1-S9 (Fasta formatted files of sequences) are in a separate text file

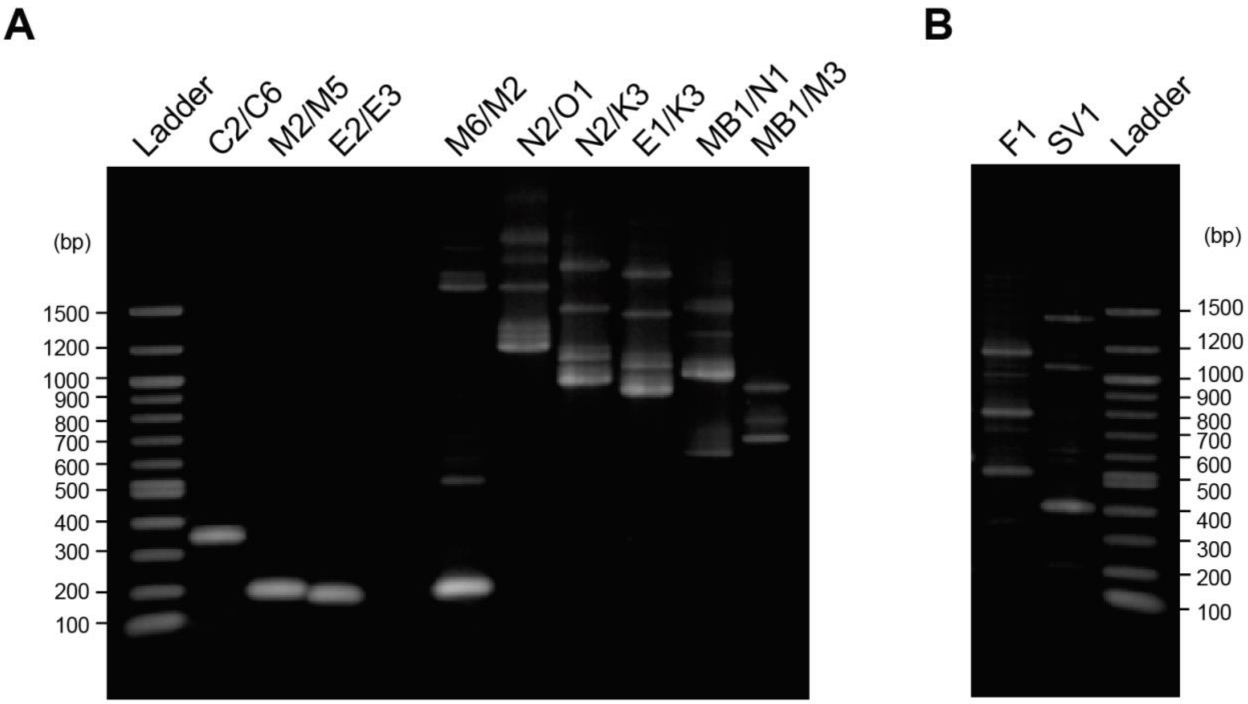


**Fig. S1 Electrophoretic analysis of *T. gondii* mtDNA PCR products.** (A-B) Genomic DNA from mitochondrial-enriched fragments was assayed with different primer pairs. (A) The labels across the top indicate primers used and primer pairs are separated by '/'. The primers are named with block names and primer sequences are provided in Table S1. Lanes 2-4 represent amplicons from cytochrome-specific primers that avoid NUMTs but amplify a small fragment of each cytochrome gene; (B) In some cases, just one primer produced multiple amplicons.

**Fig. S2 – Annotated *Toxoplasma gondii* RH sequenced mtDNA PCR products.** Fasta format sequences for these products can be found in Dataset_S1.

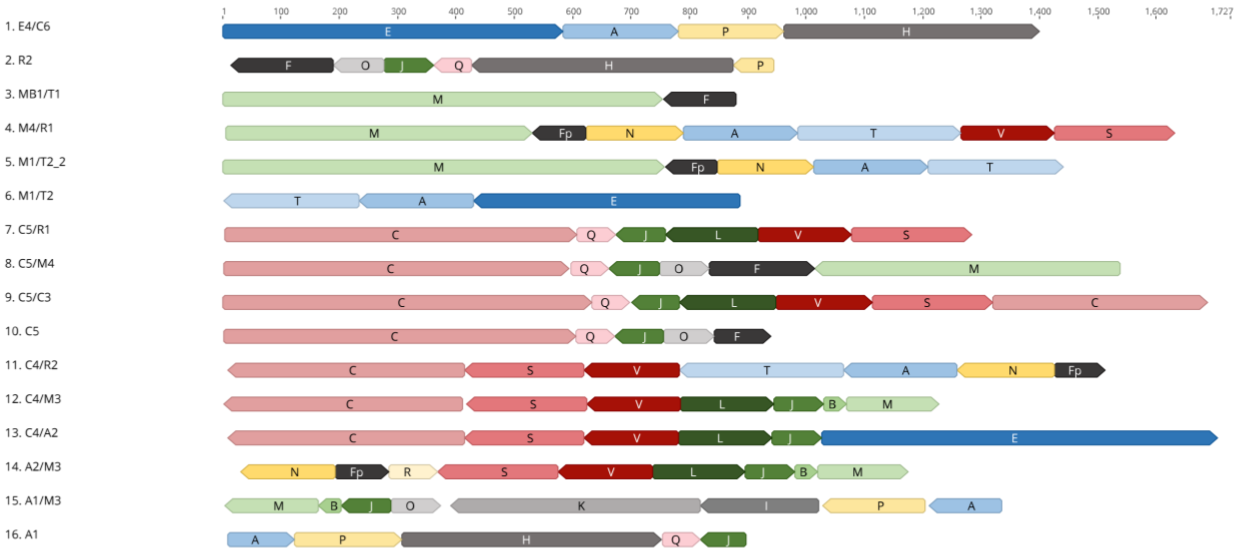

**Fig. S3 - Annotation of select A) *T. gondii* and B) *N. caninum* genomic Sanger reads from NCBI SRA database using the 21 sequence blocks from the respective organism**

***A) Toxoplasma gondii***

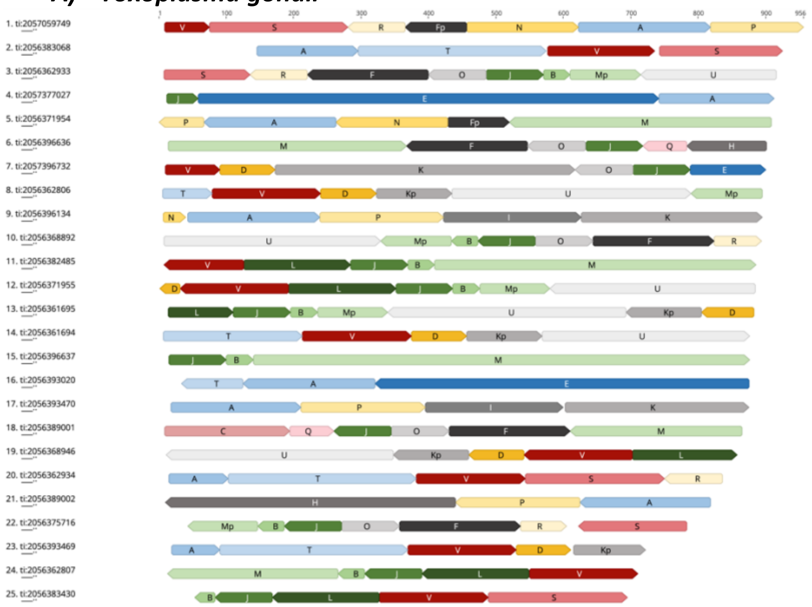


***B) Neospora caninum***

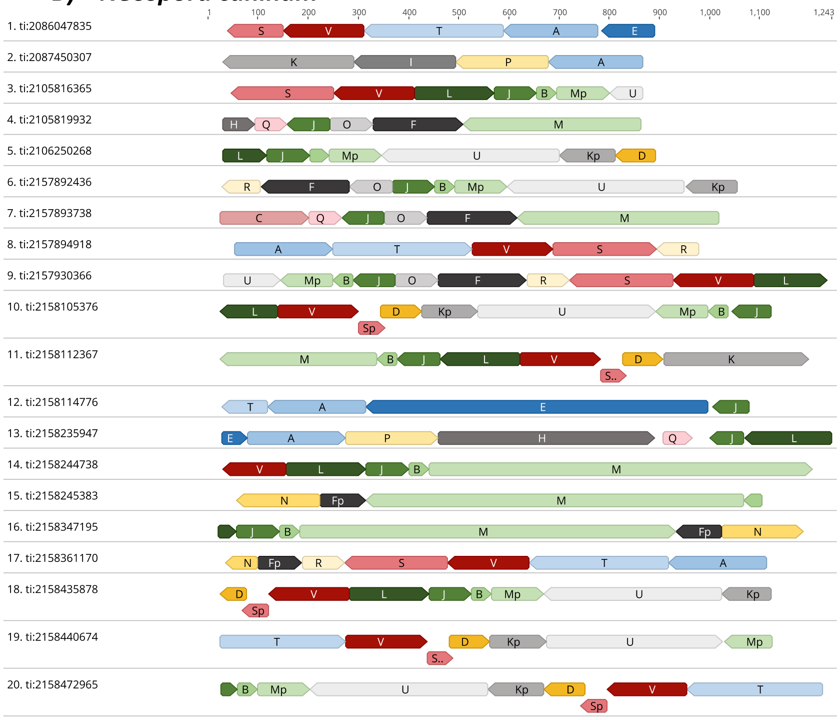


*When block R and F/Fp occur next to each other in *Neospora canin*um, the last 4 bp of F/Fp are missing and instead this junction contains the 2 nt AC/GT depending on orientation

**Fig. S4 - Annotation of select A) *T. gondii* and B) *N. caninum* Sanger EST reads from the NCBI EST database using the 21 sequence blocks from the respective organism.**

### A) Annotated *Toxoplasma gondii* EST reads

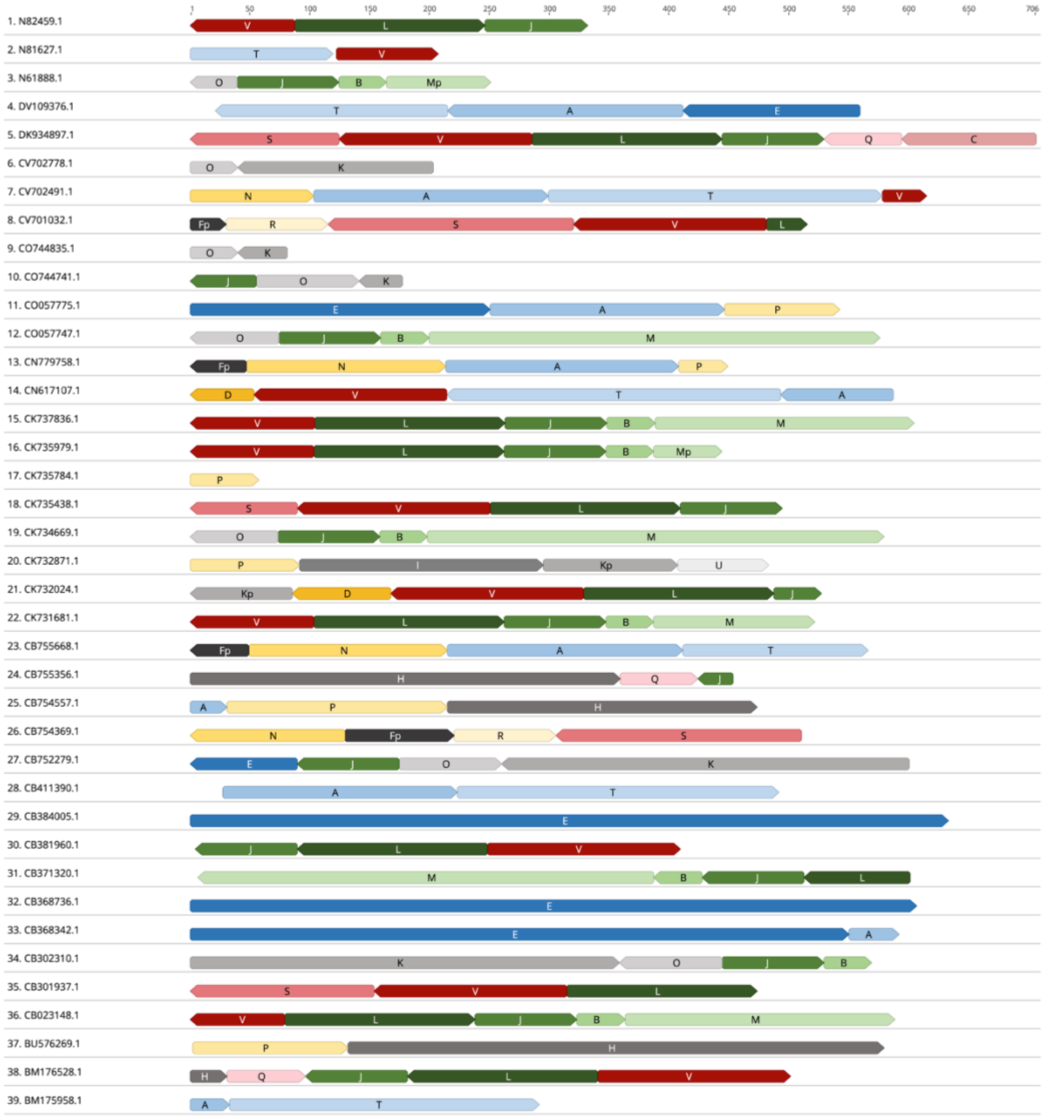

### B) Annotated *Neospora caninum* Sanger EST reads

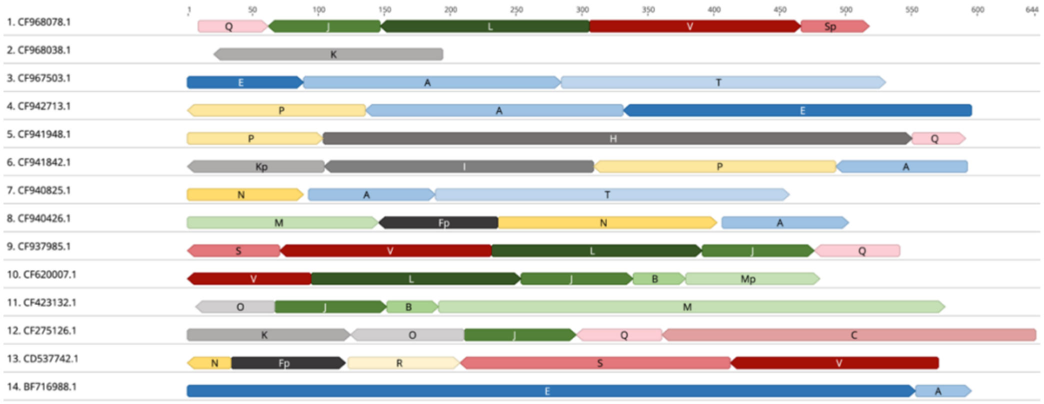

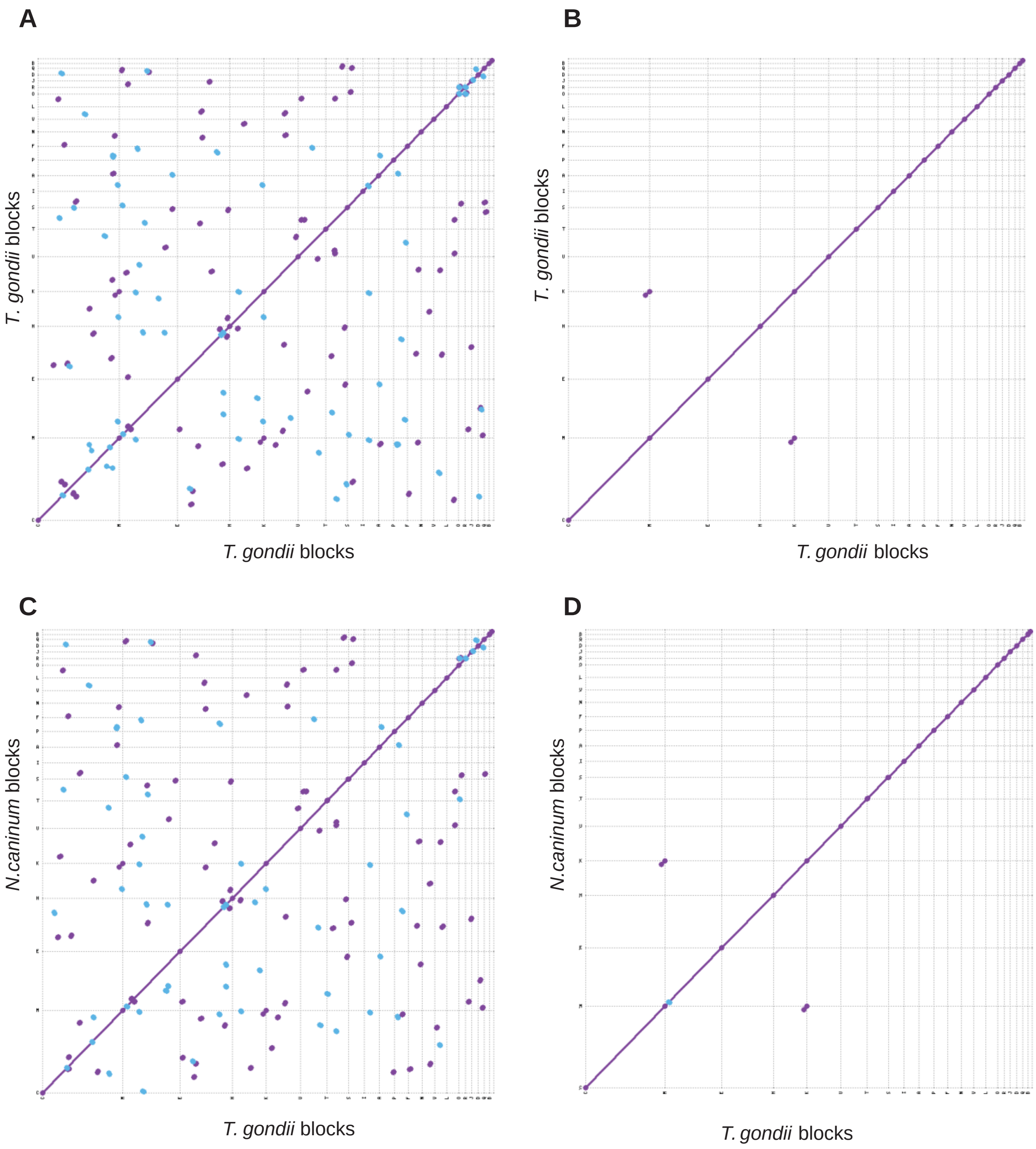


**Fig. S5 Dot plot comparisons of *T. gondii* ME49 and *N. caninum* Nc-1 sequence blocks to detect microhomology.** (A-B) The 21 *T. gondii* mtDNA SBs were aligned against each other or (C-D) to the 21 mtDNA blocks from *N. caninum*. Matches detected with window sizes of 10 bp in (A, C) and 15 bp in (B, D) are displayed in blue or purple for forward and reverse matches respectively.

### Fig. S6 – Annotated *Toxoplasma gondii* ME49 mtDNA Nanopore reads arranged by length.

#### a) *Toxoplasma gondii* ME49 Nanopore reads 4-23 kb

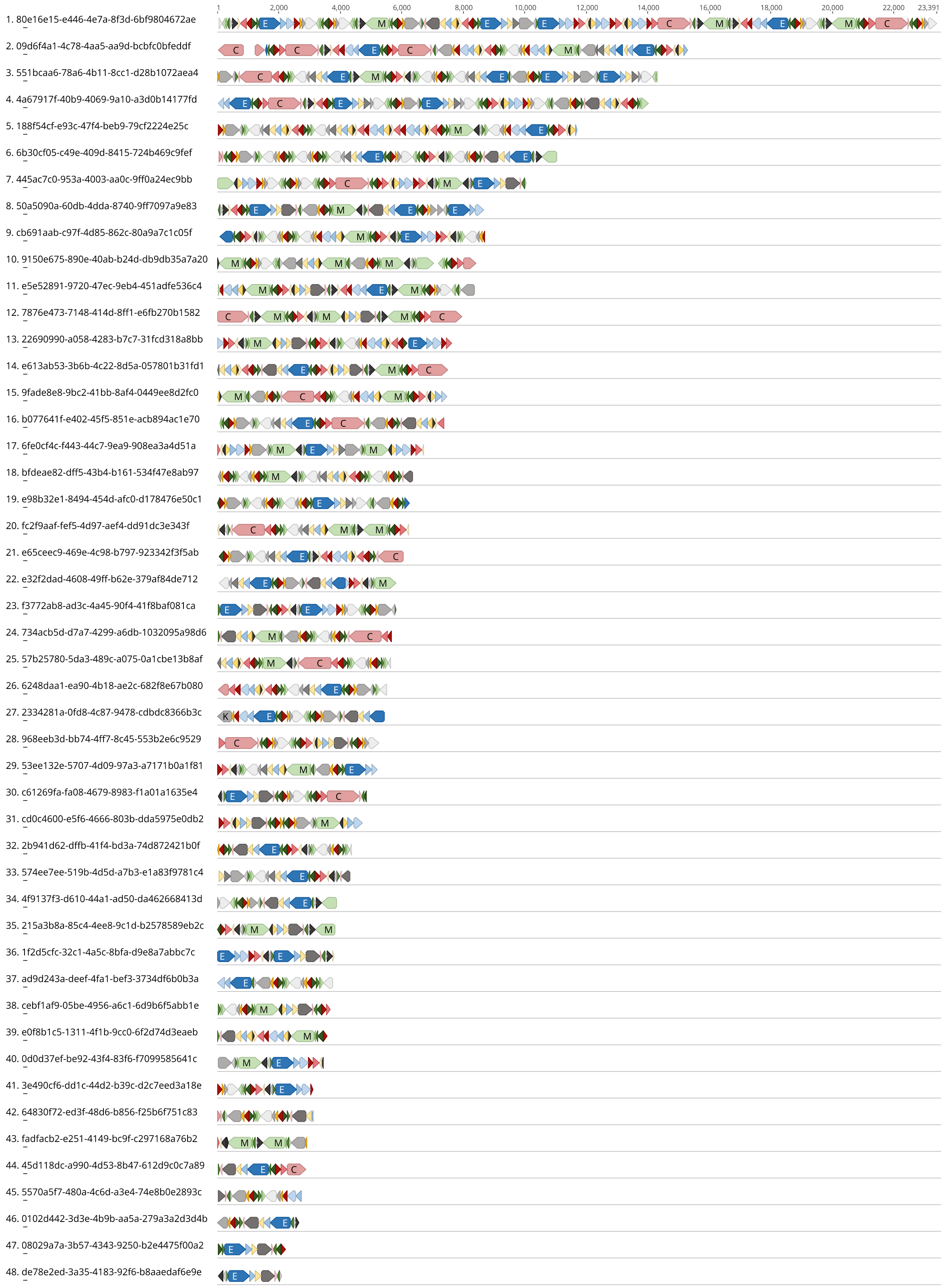

**b)** ***Toxoplasma gondii*** **ME49 Nanopore reads 1-4 kb**

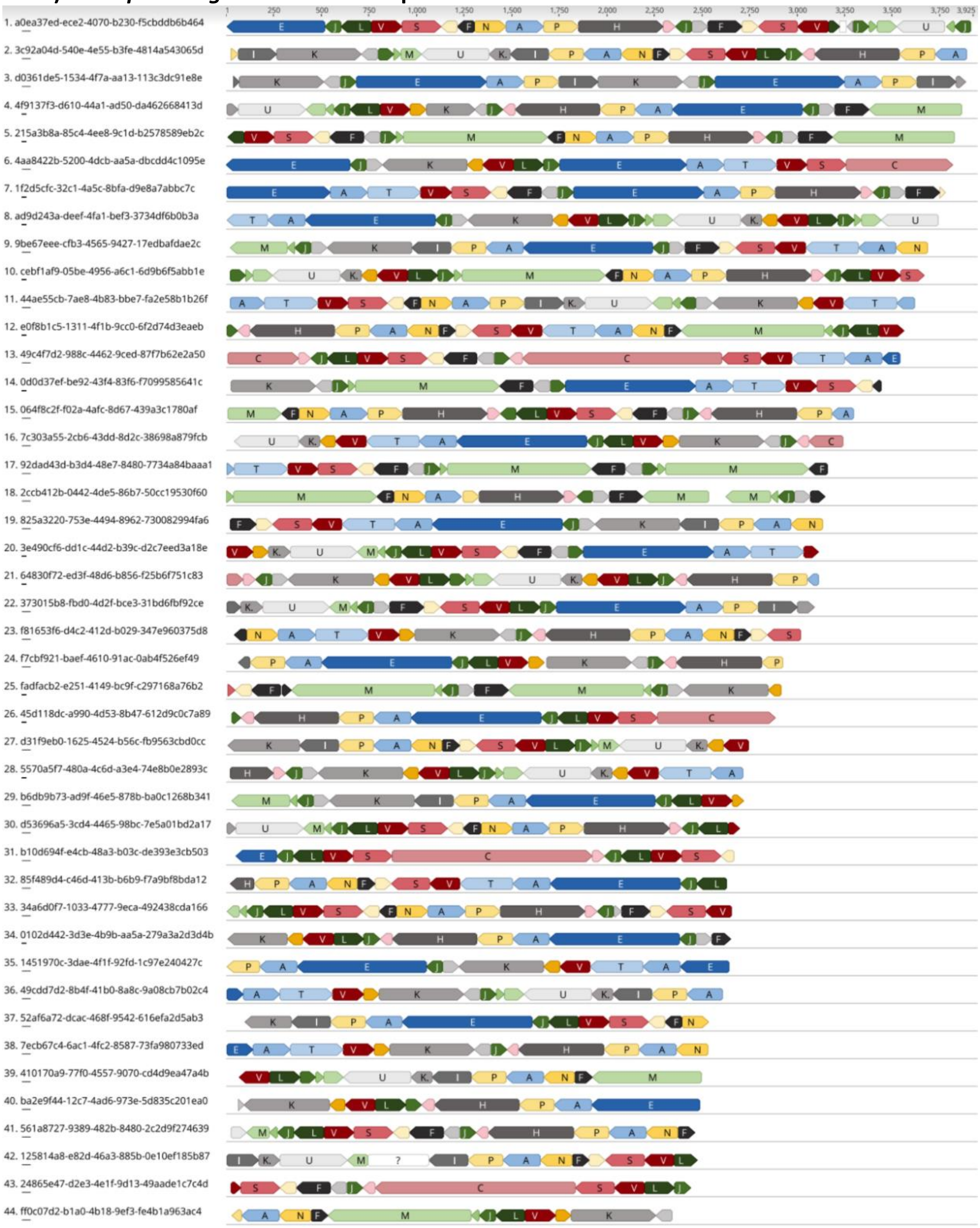

### *Toxoplasma gondii* ME49 Nanopore reads 1-4 kb (continued)

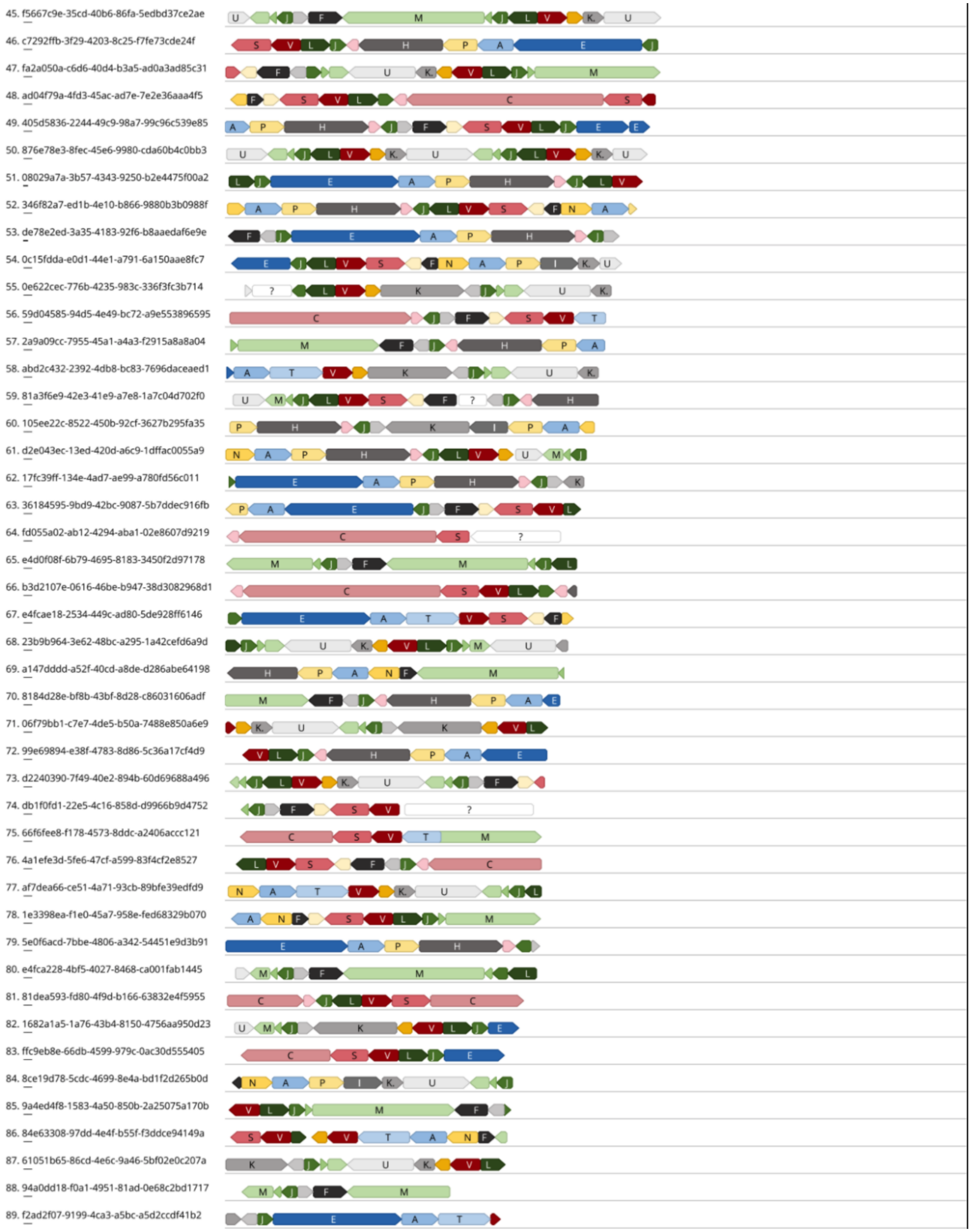

### *Toxoplasma gondii* ME49 Nanopore reads 1-4 kb (continued)

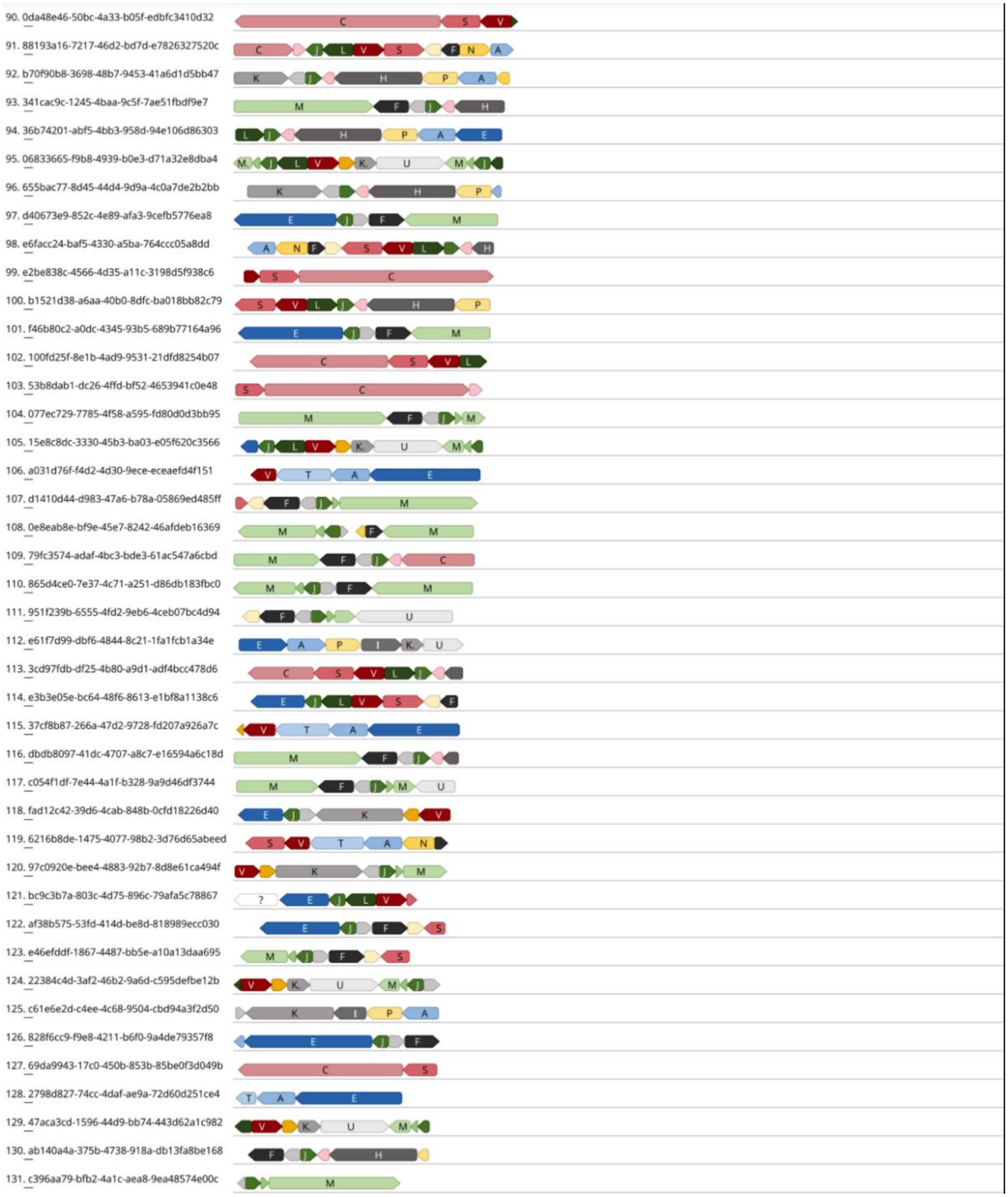

**Fig. S7. Multiple sequence alignments of mitochondrial cytochrome proteins.** Multiple sequence alignment of COXI, COXIII and COB proteins (A-C) and genes (D-F) from the apicomplexan parasites *P. falciparum*, *E. tenella*, *T. gondii* and *N. caninum*. *T. gondii* and *N. caninum* cytochrome sequences were identified as described in the methods and results. Codon 'ATA' was annotated as the start Methionine for the COXI protein in *T. gondii* and *N. caninum*. Amino acid and corresponding CDS nucleotide sequences were downloaded from NCBI for *P. falciparum* (AAC63390.2, AAC63389.2, AAC63391.1) and *E. tenella* (BAJ25753.1, BAJ25754.1, BAJ25752.1).

### A) Cytochrome B

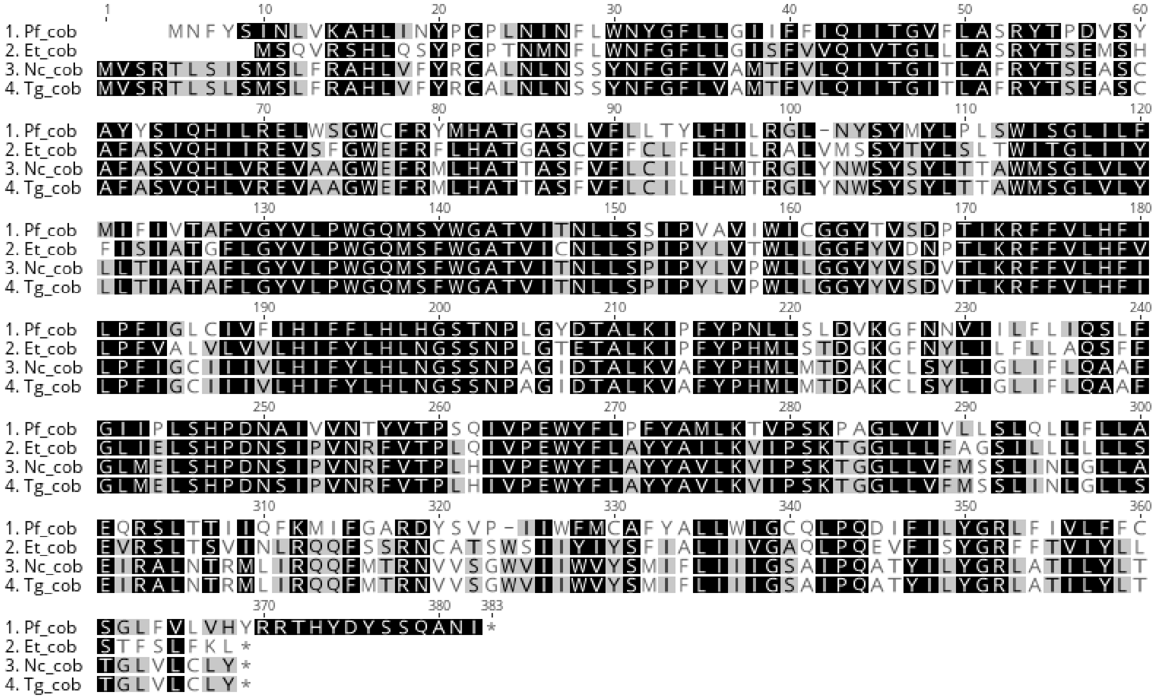

### B) Cytochrome oxidase I

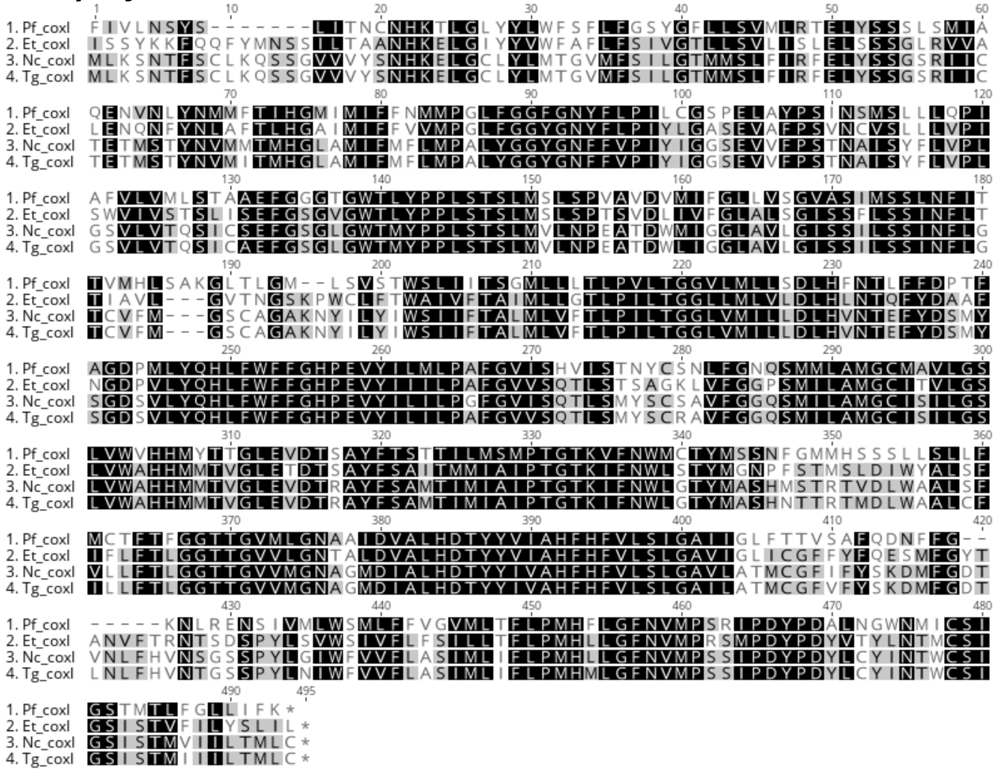


### C) Cytochrome oxidase III

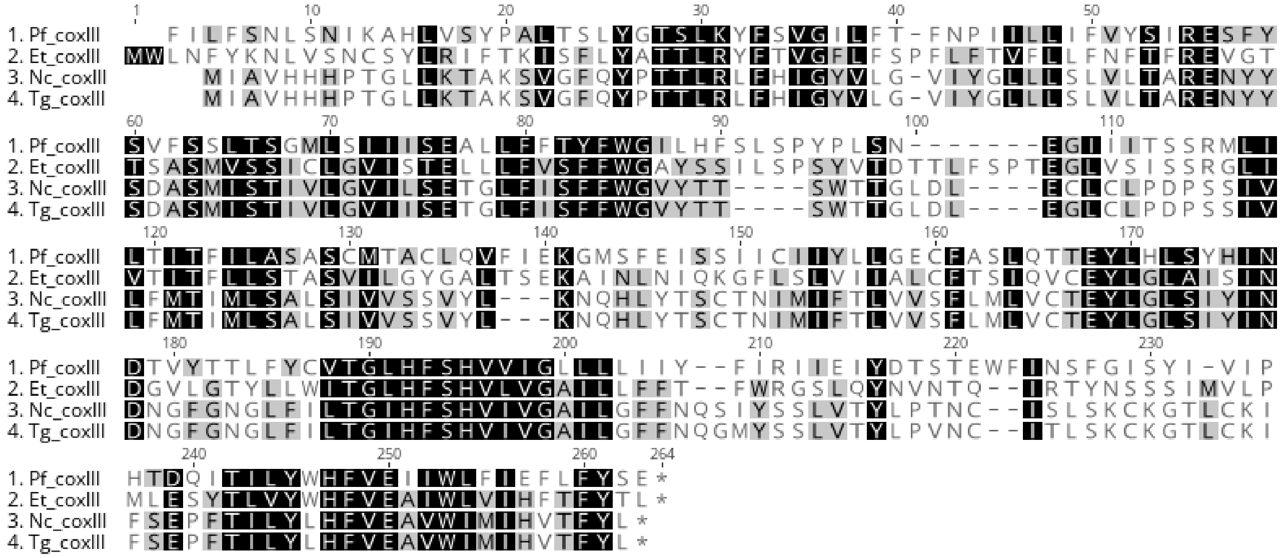

### D) Cytochrome B

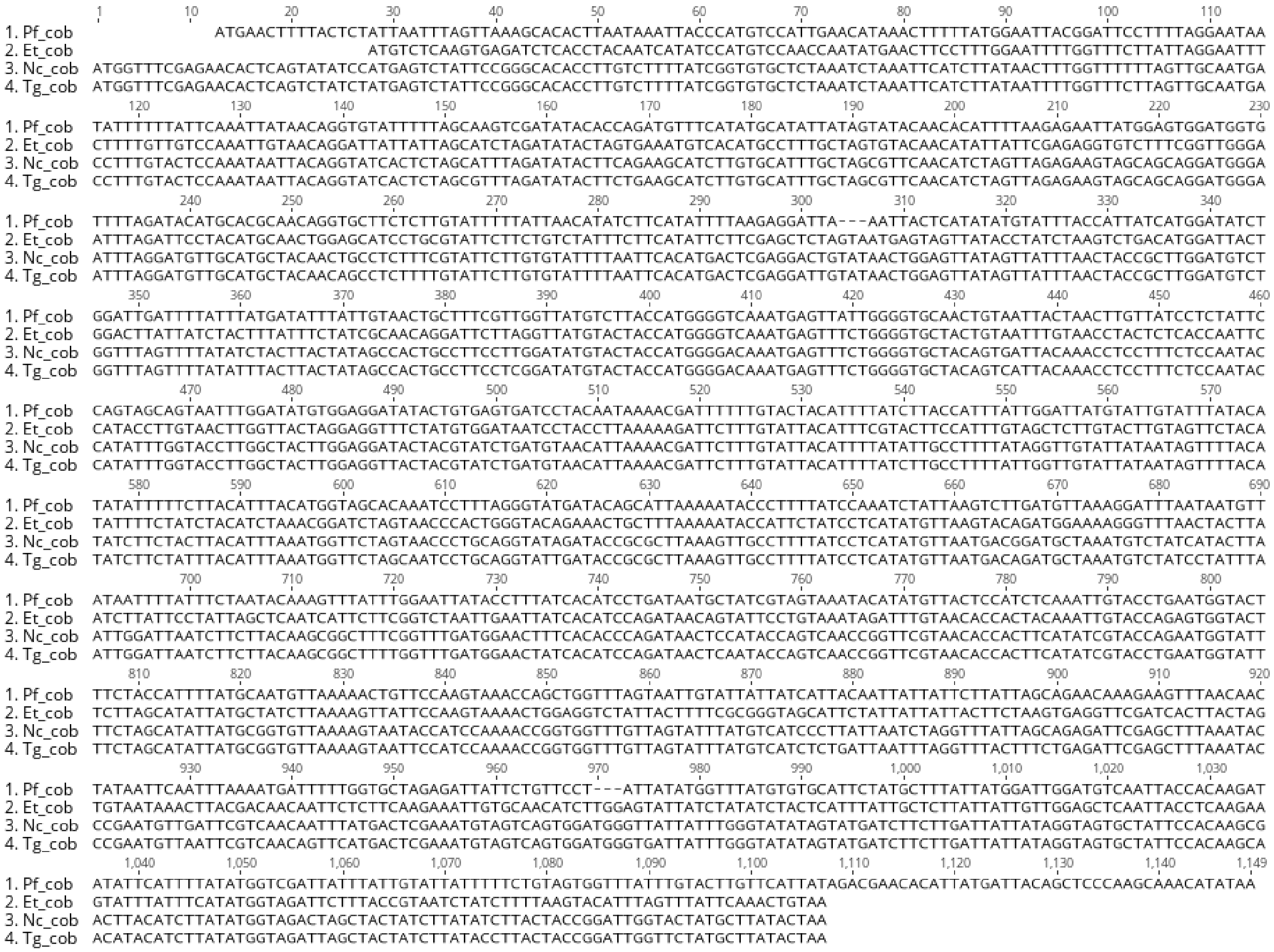

### E) Cytochrome oxidase I

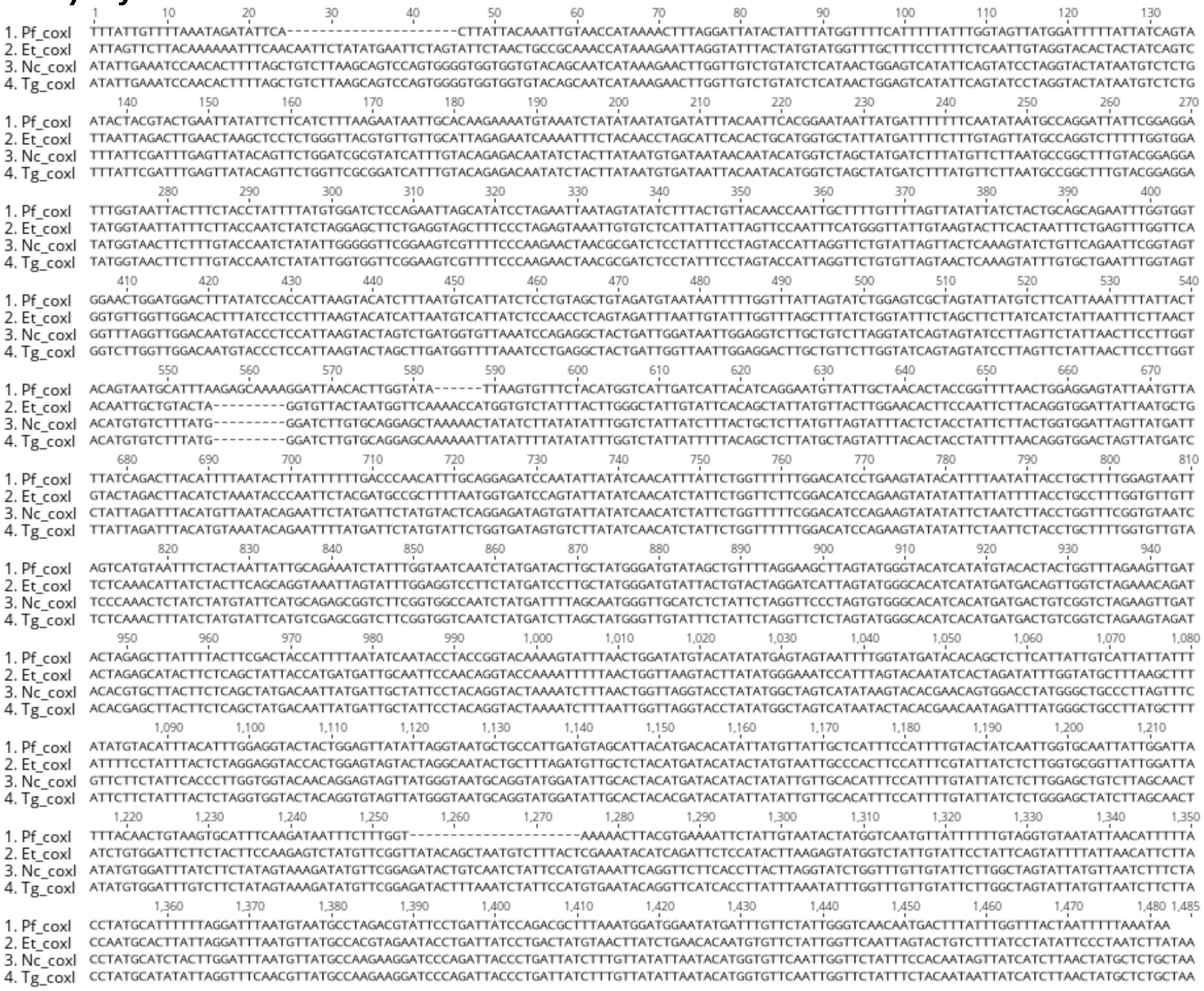

### F) Cytochrome oxidase III

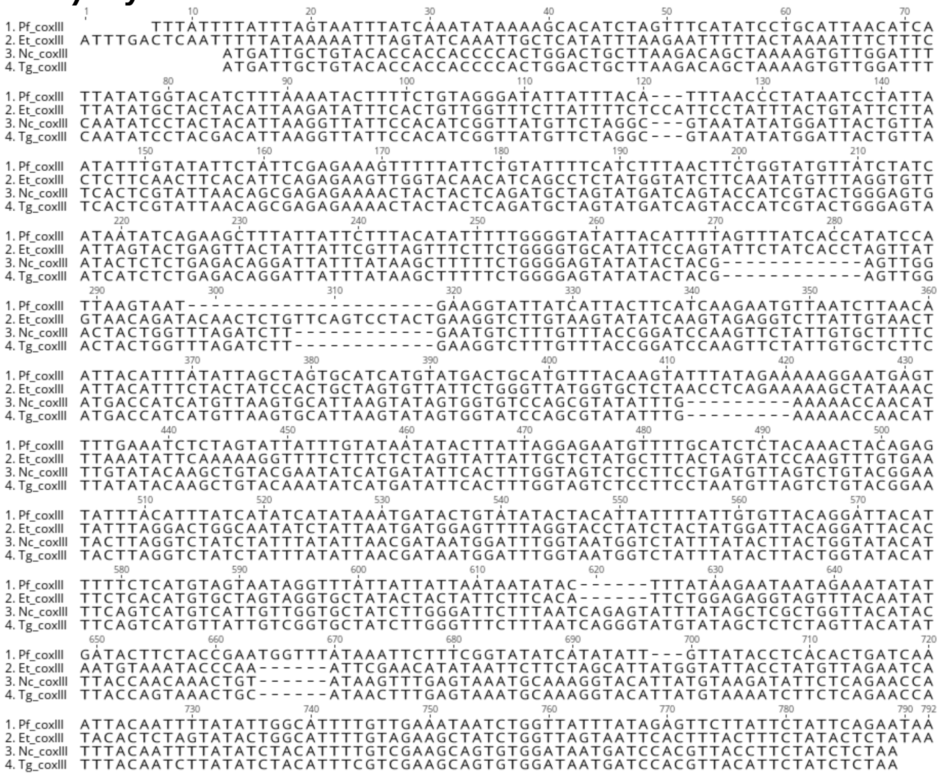

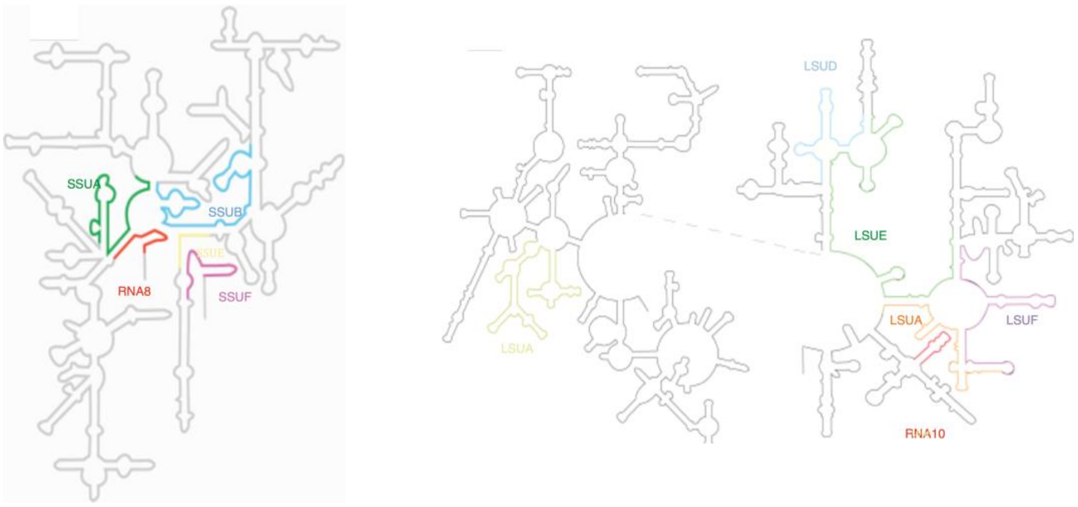


**Fig. S8. Predicted mtDNA rRNA fragments in *T. gondii* folded onto *E. coli* rRNA secondary structures.** (A) Detected *T. gondii* mtDNA small and large subunit rRNA fragments are manually folded and mapped on to conserved *E. coli* SSU and (B) LSU secondary structures. Grey lines represent *E. coli* secondary structures. *T. gondii* mtDNA rRNA fragments are shown using different colors and are labeled.

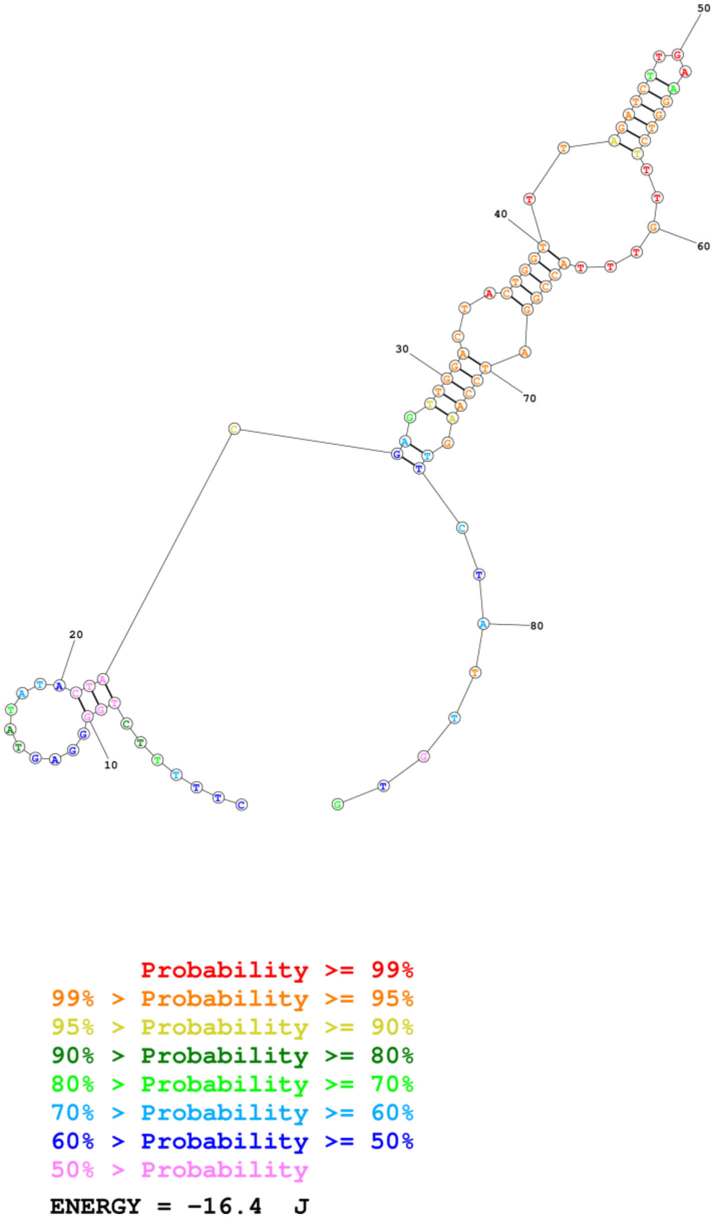


**Fig. S9 Secondary structure of sequence block J.** Structure analysis was performed with RNA structure Web Servers for RNA Secondary Structure Prediction. The Mathews Group. https://rna.urmc.rochester.edu/RNAstructureWeb/Servers/Predict1/Predict1.html

Settings: MaxExpect partition.pfs MaxExpect.ct --gamma 1 --percent 10 --structures 20 --window 3

**Fig. S10. Multiple sequence alignment of a subset of uncorrected Nanopore reads from the ENU mutant strain ELQ-316.** The wt sequence is on the first row. The mutant sequence is on the second row. The mutation , which is an A -> C, is located at position 23.

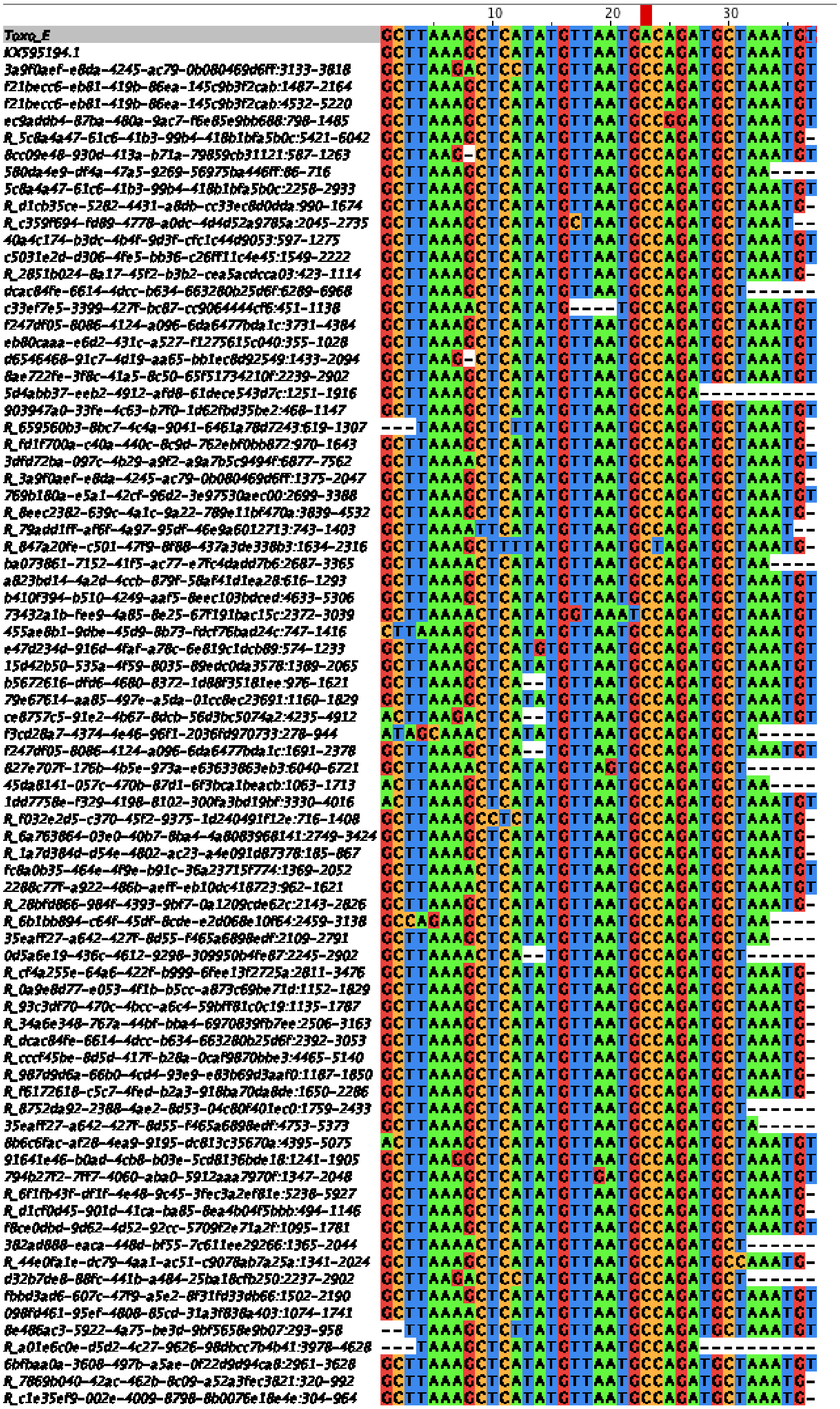

**Fig. S11 – Annotated *Neospora caninum* mtDNA Nanopore reads (2-15 kb) arranged by length**

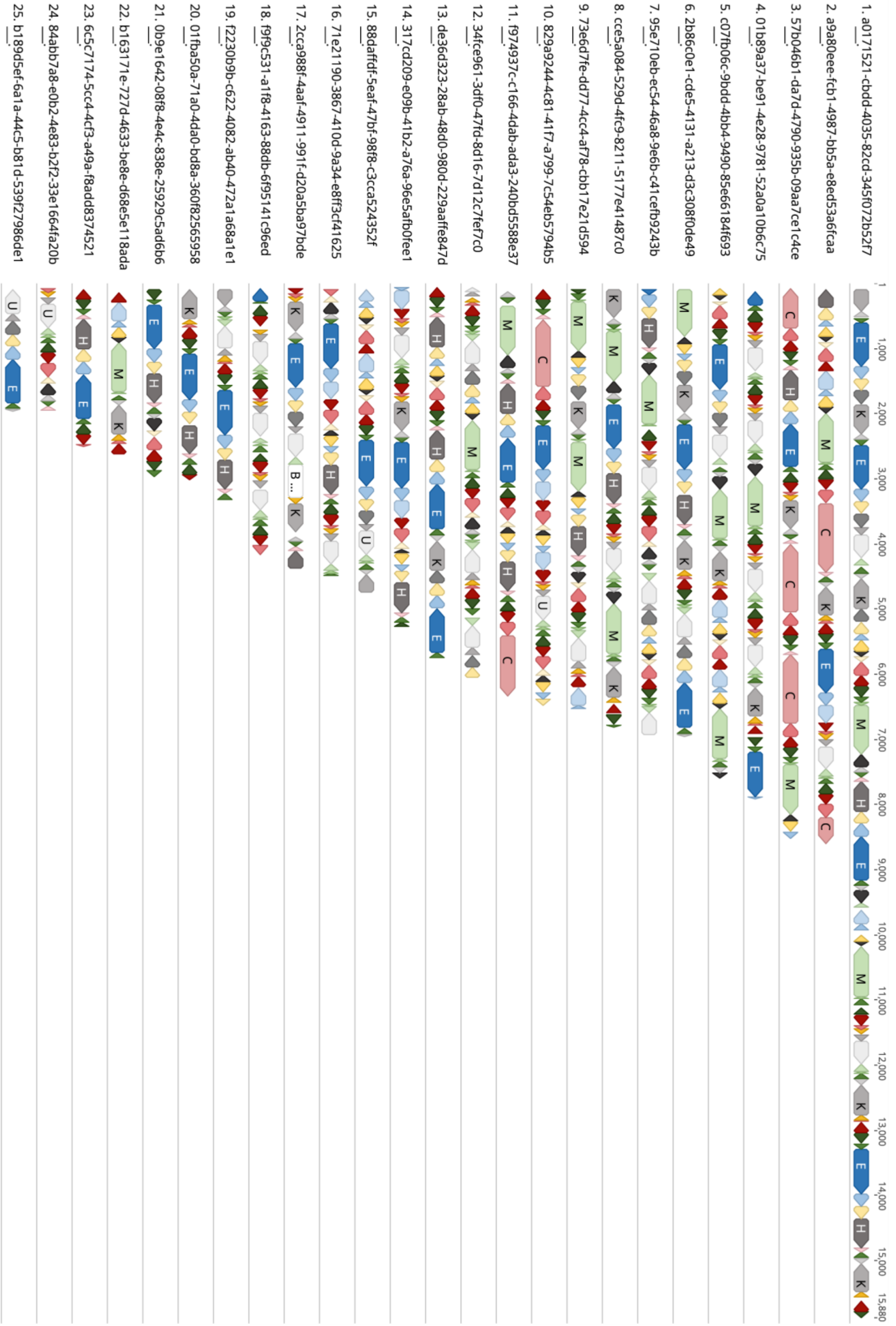

**A**

| | A | B | C | D | E | F | Fp | H | I | J | K | Kp | L | M | Mp | N | O | P | Q | R | S | T | U | V |
|---|---|---|---|---|---|---|---|---|---|---|---|---|---|---|---|---|---|---|---|---|---|---|---|---|
| A | | | | | 50 | | | | | | | | | | | 43 | | 55 | | | | 38 | | |
| B | | | | | | | | | | 98 | | | | 40 | 58 | | | | | | | | | |
| C | | | | | | | | | | | | | | | | | | | 19 | | 19 | | | |
| D | | | | | | | | | | | 31 | 39 | | | | | | | | | | | | 70 |
| E | 50 | | | | | | | | | 48 | | | | | | | | | | | | | | |
| F | | | | | | | | | | | | | | 21 | | | 48 | | | 24 | | | | |
| Fp | | | | | | | | | | | | | | 19 | | 44 | | | | 29 | | | | |
| H | | | | | | | | | | | | | | | | | | 30 | 33 | | | | | |
| I | | | | | | | | | | | 6 | 17 | | | | | | 25 | | | | | | |
| J | | 98 | | | 48 | | | | | | | | 106 | | | | 89 | | 50 | | | | | |
| K | | | | 31 | | | | | 6 | | | | | | | | 37 | | | | | | | |
| Kp | | | | 39 | | | | | 17 | | | | | | | | | | | | | | 55 | |
| L | | | | | | | | | | 106 | | | | | | | | | | | | | | 108 |
| M | | 40 | | | | 21 | 19 | | | | | | | | | | | | | | | | | |
| Mp | | 58 | | | | | | | | | | | | | | | | | | | | | 58 | |
| N | 43 | | | | | | 44 | | | | | | | | | | | | | | | | | |
| O | | | | | | 48 | | | | 89 | 37 | | | | | | | | | | | | | |
| P | 55 | | | | | | | 30 | 25 | | | | | | | | | | | | | | | |
| Q | | | 19 | | | | | 33 | | 50 | | | | | | | | | | | | | | |
| R | | | | | | 24 | 29 | | | | | | | | | | | | | | 54 | | | |
| S | | | 19 | | | | | | | | | | | | | | | | | 54 | | | | 72 |
| T | 38 | | | | | | | | | | | | | | | | | | | | | | | 34 |
| U | | | | | | | | | | | | 55 | | | 58 | | | | | | | | | |
| V | | | | 70 | | | | | | | | | 108 | | | | | | | | 72 | 34 | | |

**B**

| | A | B | C | D | E | F | Fp | H | I | J | K | Kp | L | M | Mp | N | O | P | Q | R | S | Sp | T | U | V |
|---|---|---|---|---|---|---|---|---|---|---|---|---|---|---|---|---|---|---|---|---|---|---|---|---|---|
| A | | | | | 25 | | | | | | | | | | | 20 | | 32 | | | | | 13 | | |
| B | | | | | | | | | | 39 | | | | 15 | 24 | | | | | | | | | | |
| C | | | | | | | | | | | | | | | | | | | 4 | | 7 | | | | |
| D | | | | | | | | | | | 13 | 19 | | | | | | | | | | 22 | | | |
| E | 25 | | | | | | | | | 26 | | | | | | | | | | | | | | | |
| F | | | | | | | | | | | | | | 9 | | | 15 | | | 6 | | | | | |
| Fp | | | | | | | | | | | | | | 8 | | 19 | | | | 12 | | | | | |
| H | | | | | | | | | | | | | | | | | | 19 | 19 | | | | | | |
| I | | | | | | | | | | | 5 | 9 | | | | | | 14 | | | | | | | |
| J | | 39 | | | 26 | | | | | | | | 49 | | | | 39 | | 23 | | | | | | |
| K | | | | 13 | | | | | 5 | | | | | | | | 23 | | | | | | | | |
| Kp | | | | 19 | | | | | 9 | | | | | | | | | | | | | | | 28 | |
| L | | | | | | | | | | 49 | | | | | | | | | | | | | | | 47 |
| M | | 15 | | | | 9 | 8 | | | | | | | | | | | | | | | | | | |
| Mp | | 24 | | | | | | | | | | | | | | | | | | | | | | 26 | |
| N | 20 | | | | | | 19 | | | | | | | | | | | | | | | | | | |
| O | | | | | | 15 | | | | 29 | 23 | | | | | | | | | | | | | | |
| P | 32 | | | | | | | 18 | 14 | | | | | | | | | | | | | | | | |
| Q | | | 4 | | | | | 19 | | 23 | | | | | | | | | | | | | | | |
| R | | | | | | 6 | 12 | | | | | | | | | | | | | | 19 | | | | |
| S | | | 7 | | | | | | | | | | | | | | | | | 19 | | | | | 27 |
| Sp | | | | 32 | | | | | | | | | | | | | | | | | | | | | 31 |
| T | 13 | | | | | | | | | | | | | | | | | | | | | | | | 12 |
| U | | | | | | | | | | | | 28 | | | 26 | | | | | | | | | | |
| V | | | | | | | | | | | | | 47 | | | | | | | | 27 | 31 | 12 | | |

**C**

| | A | B | C | D | E | F | Fp | H | I | J | K | Kp | L | M | Mp | N | O | P | Q | R | S | T | U | V |
|---|---|---|---|---|---|---|---|---|---|---|---|---|---|---|---|---|---|---|---|---|---|---|---|---|
| A | | | | | 59 | | | | | | | | | | | 23 | | 16 | | | | 86 | | |
| B | | | | | | | | | | 89 | | | | 72 | 23 | | | | | | | | | |
| C | | | | | | | | | | | | | | | | | | | 21 | | 34 | | | |
| D | | | | | | | | | | | 7 | 23 | | | | | | | | | | | | 30 |
| E | 59 | | | | | | | | | 15 | | | | | | | | | | | | | | |
| F | | | | | | | | | | | | | | 5 | | | 19 | | | 8 | | | | |
| Fp | | | | | | | | | | | | | | 4 | | 18 | | | | 16 | | | | |
| H | | | | | | | | | | | | | | | | | | 13 | 11 | | | | | |
| I | | | | | | | | | | | 11 | 12 | | | | | | 25 | | | | | | |
| J | | 89 | | | 15 | | | | | | | | 73 | | | | 62 | | | | | | | |
| K | | | | | | | | | 9 | | | | | | | | 25 | | | | | | | |
| Kp | | | | 25 | | | | | 12 | | | | | | | | | | | | | | 22 | |
| L | | | | | | | | | | 73 | | | | | | | | | | | | | | 58 |
| M | | 73 | | | | 5 | 6 | | | | | | | | | | | | | | | | | |
| Mp | | 23 | | | | | | | | | | | | | | | | | | | | | 28 | |
| N | 63 | | | | | | 18 | | | | | | | | | | | | | | | | | |
| O | | | | | | 19 | | | | 26 | 25 | | | | | | | | | | | | | |
| P | 25 | | | | | | | 13 | 25 | | | | | | | | | | | | | | | |
| Q | | | 21 | | | | | 11 | | 61 | | | | | | | | | | | | | | |
| R | | | | | | 4 | 16 | | | | | | | | | | | | | | 23 | | | |
| S | | | 34 | | | | | | | | | | | | | | | | | 23 | | | | 59 |
| T | 86 | | | | | | | | | | | | | | | | | | | | | | | 17 |
| U | | | | | | | | | | | | 22 | | | 28 | | | | | | | | | |
| V | | | | 31 | | | | | | | | | 58 | | | | | | | | 59 | 17 | | |

**D**

| | A | B | C | D | E | F | Fp | H | I | J | K | Kp | L | M | Mp | N | O | P | Q | R | S | Sp | T | U | V |
|---|---|---|---|---|---|---|---|---|---|---|---|---|---|---|---|---|---|---|---|---|---|---|---|---|---|
| A | | | | | 3 | | | | | | | | | | | 9 | | 5 | | | | | 5 | | |
| B | | | | | | | | | | 3 | | | | | 1 | | | | | | | | | | |
| C | | | | | | | | | | | | | | | | | | | | | 1 | | | | |
| D | | | | | | | | | | | 1 | | | | | | | | | | | 1 | | | |
| E | 3 | | | | | | | | | | | | | | | | | | | | | | | | |
| F | | | | | | | | | | | | | | | | | | | | | | | | | |
| Fp | | | | | | | | | | | | | | | | | | | | | | | | | |
| H | | | | | | | | | | | | | | | | | | | 3 | | | | | | |
| I | | | | | | | | | | | | 2 | | | | | | 3 | | | | | | | |
| J | | 3 | | | 1 | | | | | | | | | | | | 3 | | 4 | | | | | | |
| K | | | | 1 | | | | | | | | | 3 | | | | 1 | | | | | | | | |
| Kp | | | | | | | | | 2 | | | | | | | | | | | | | | | | |
| L | | | | | | | | | | 3 | | | | | | | | | | | | | | | 4 |
| M | | | | | | | | | | | | | | | | | | | | | | | | | |
| Mp | | 1 | | | | | | | | | | | | | | | | | | | | | | | |
| N | 9 | | | | | | | | | | | | | | | | | | | | | | | | |
| O | | | | | | | | | | 3 | | | | | | | | | | | | | | | |
| P | 5 | | | | | | | | 3 | | | | | | | | | | | | | | | | |
| Q | | | | | | | | 3 | | 4 | | | | | | | | | | | | | | | |
| R | | | | | | | | | | | | | | | | | | | | | | | | | |
| S | | | 1 | | | | | | | | | | | | | | | | | | | | | | 4 |
| Sp | | | | 1 | | | | | | | | | | | | | | | | | | | | | 1 |
| T | 5 | | | | | | | | | | | | | | | | | | | | | | | | |
| U | | | | | | | | | | | | | | | | | | | | | | | | | |
| V | | | | | | | | | | | | | 4 | | | | | | | 2 | 4 | 1 | | | |

**Fig. S12. Upstream and downstream sequence blocks in *T. gondii* ME49 and *N. caninum* Nanopore reads.** Data are based on 48 mtDNA-specific Nanopore reads provided in Fig. S6, Dataset S3. The identity of blocks located immediately at the 5′ and 3′ end of each of the 21 sequences blocks and 3 partials were counted, and the counts are shown as a matrix. The number of times a block on the y-axis is present at the 5′ or 3′ end in reference to the block on the x-axis is indicated in blue or red respectively. If two blocks were not found next to each other that corresponding square is blank. Only upstream or downstream block occurrences appearing >3 were counted to avoid anomalies (Fig. S6; Table S6). (A) Data derived from *T. gondii* Nanopore DNA read (B) *N. caninum* Nanopore DNA reads, (C) *T. gondii* Nanopore RNA reads and (D) *N. caninum* Sanger EST reads. Only upstream or downstream block occurrences appearing >3 times were counted to avoid anomalies (Tables S6 & S11). Please note the absence of a block relationships in (D) is likely due to missing data since the dataset was sparse.

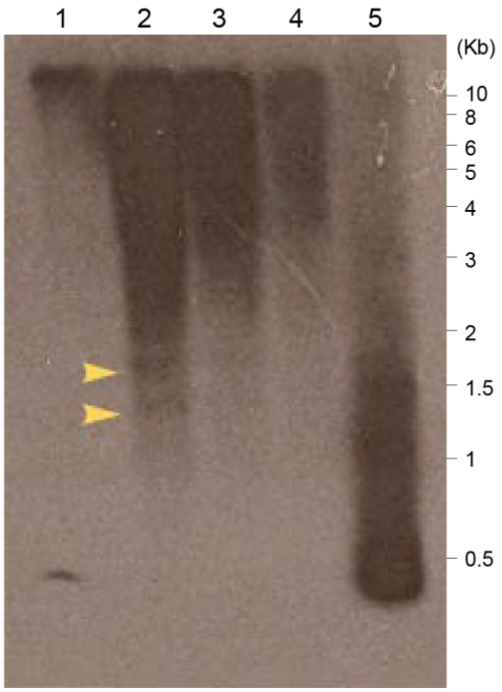


**Fig. S13. *T. gondii* mtDNA is variable in size and does not exist as a tandem repeat unit.** Southern blot analysis. Total DNA from the *T. gondii* RH strain was hybridized to a 350 bp *coxI* probe under very stringent conditions. Lane 1: Undigested 2 µg genomic DNA; 2: *XhoI* Digested 15 µg genomic DNA; 3: *XhoI* Digested 5 µg genomic DNA; 4: *XhoI* Digested 2 µg genomic DNA; 5: positive control 350 bp probe fragment. *XhoI* cuts only once in the mitochondrial SBs, it cuts inside sequence block E of the *cob* gene. The yellow arrows point to distinct bands (~1.65 and 1.35 kb) on the blot. Size markers are indicated.

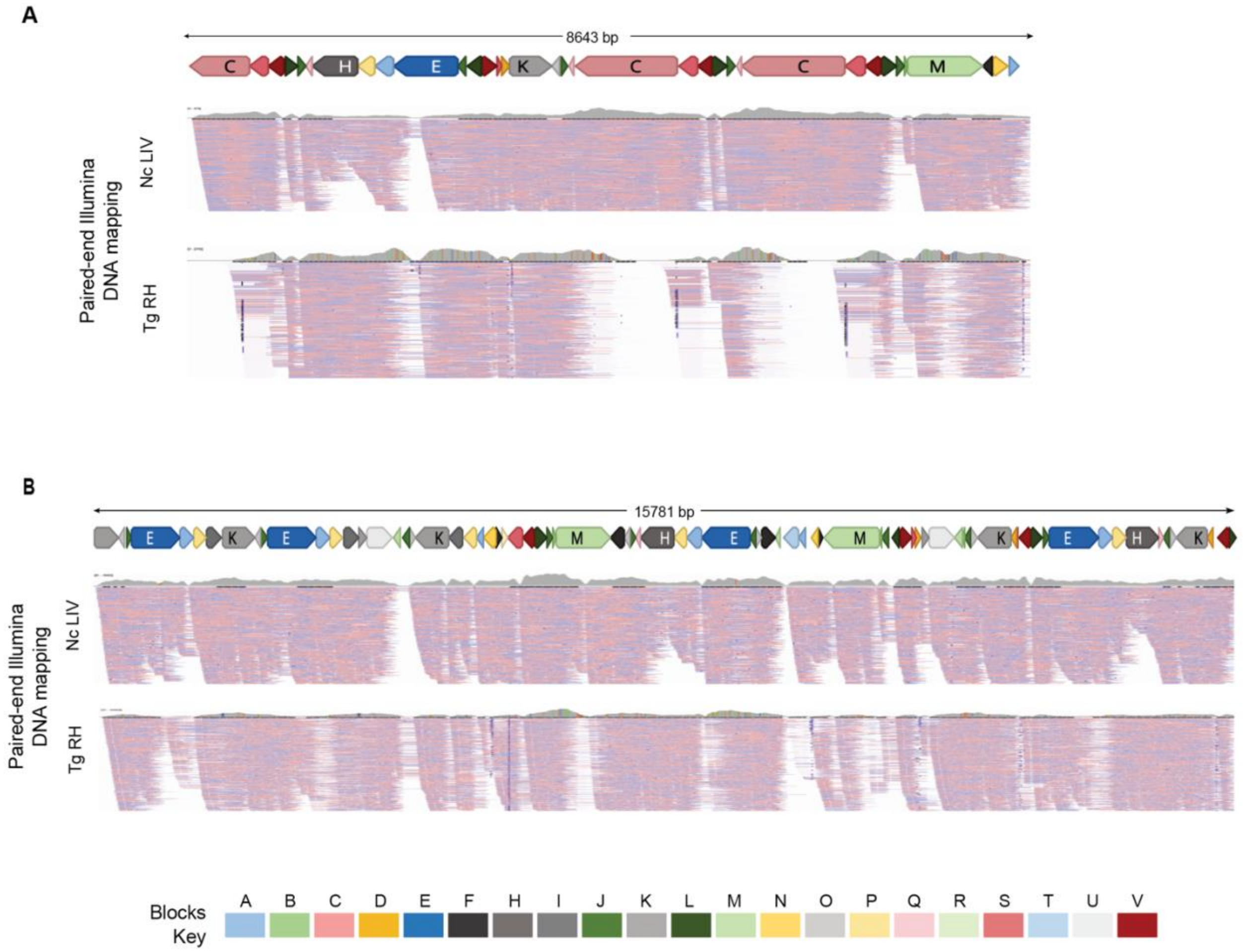


**Fig. S14. *N. caninum* nanopore sequence comparison and degeneration.** (A-B) SBs were annotated on two *N. caninum* Nc-1 nanopore sequences reads at scale. The blocks are colored as shown in the key at the bottom of the figure. The two 'Paired-end Illumina DNA mapping' tracks show paired-end read mapping of *N. caninum* (Nc) LIV (ERR012900) and *T. gondii* (Tg) RH (SRR521957) mtDNA-specific reads. Mapping of Nc LIV reads required 100% nucleotide identity whereas 1% mismatch was allowed for mapping *T. gondii* RH reads. Reads were independently mapped to each of the Nanopore mtDNA reads and visualized using IGV. Red and blue lines below each read indicate the mapped Illumina paired-ends.

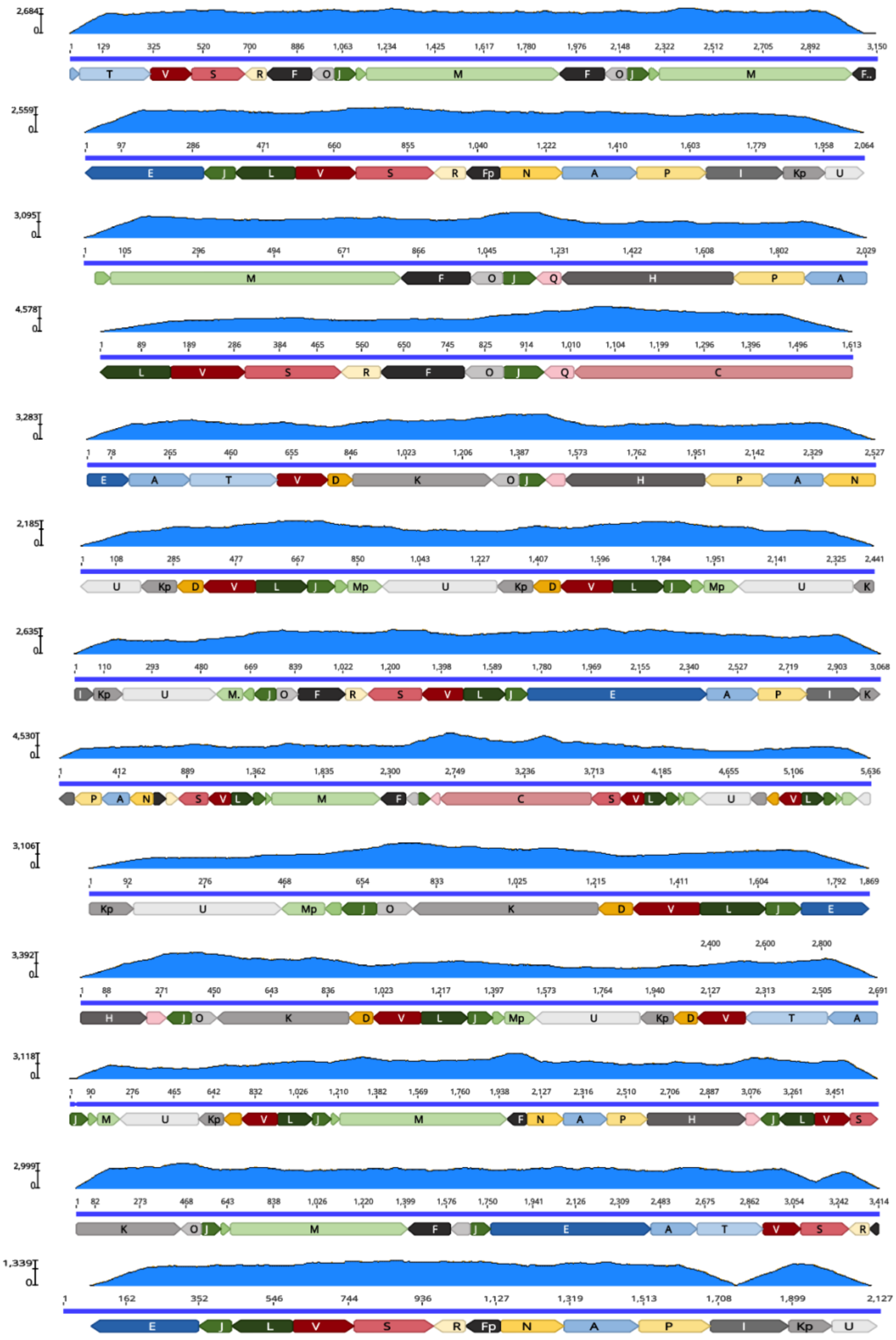

**Fig. S15 Nanopore reads predicted to circularize.** *T. gondii* ME49 Illumina mapping frequency support for predicted circular sequences. Similar to Fig. 9, mapped read counts are displayed above each Nanopore read in blue. Some reads show fairly uniform coverage and others do not. Scale is as indicated for each read. Full-length cytochrome genes are observed.

### Table S1. Primer sequences and corresponding *Toxoplasma gondii* sequence blocks

| Primer Name (Based on sequence block) | Sequence | Sequence Block Co-ordinates |
|---|---|---|
| A1 | 5'-GTT TGA TGG AAC TAT CAC ATC CAG ATA A-3' | 153-126 |
| A2 | 5'-CAA AAG CCG CTT GTA AGA AGA TTA ATC-3' | 154-180 |
| BM1 | 5'-AGT GCA TTA AGT ATA GTG GTA TCC AGC GTA-3' | B (19-1) M (1-11) |
| C1 | 5'-GGG TTC TGT GTT AGT AAC TCA AAG TA-3' | 1-24 |
| C2 | 5'-TGA TTG GTT AAT TGG AGG ACT TGC TGT-3' | 110-136 |
| C3 | 5'-AAG ACA CTA TCA CCA GAA TAC ATA GAA TC-3' | 367-339 |
| C4 | 5'-AGA TAC AAC ACC AAA AGC AGG TAG AA-3' | 417-442 |
| C5 | 5'-TTC TAC CTG CTT TTG GTG TTG TAT CT-3' | 442-417 |
| C6 | 5'-GTT TGA GAT ACA ACA CCA AAA GCA GG-3' | 447-422 |
| C7 | 5'-ACA TGA TGA CTG TCG GTC TAG AAG TAG A-3' | 546-573 |
| C8 or H1 | 5'-GAT AAT CAG GGT AAT CTG GGA TCC-3' | 1040-1017 |
| C9 | 5'-ATC AGG GTA ATC TGG GAT CCT T-3' | 1037-1016 |
| D1 | 5'-TTC AGG AGC ATA CCG TTA TAT TCG ATG AT-3' | 37-65 |
| D2 | 5'-ATC ATC GAA TAT AAC GGT ATG CTC CTG AA-3' | 65-37 |
| D3 | 5'-TAA TGT GAA CAC ATA AGA TCA TCG AAT ATA AC-3' | 83-51 |
| E1 | 5'-TTG ATA CCG CGC TTA AAG TTG CCT-3' | 59-36 |
| E2 | 5'-CGT AGT AAC CTC CAA GTA GCC AAG G-3' | 183-207 |
| E3 | 5'-AAC TAC CGC TTG GAT GTC TGG TTT AG-3' | 352-327 |
| E4 | 5'-TGG TTT CTT AGT TGC AAT GAC CTT TGT-3' | 583-557 |
| E5 | 5'-TCT TTT ATC GGT GTG CTC TAA ATC T-3' | 58-82 |
| F1 | 5'-GGG GAC AAA AAG ACA TCA CGA T-3' | 179-162 |
| K1 | 5'-CTG AGT ACG TAA GGA AAA GGA AAG G-3' | 1-25 |
| K2 | 5'-AAC GTA ACA AAC CTC GAG GCA AAG A-3' | 200-176 |
| K3 | 5'-TCT TTG CCT GGA GGT TTG TTA CGT T-3' | 176-200 |
| MB1 | 5'-TAC GCT GGA TAC CAC TAT ACT TAA TGC ACT-3' | M (11-1) B (1-19) |
| M1 | 5'-TAT CCA GCG TAT ATT TGA AAA ACC AAC ATT-3' | 1-30 |
| M2 | 5'-TCA TGT TAT TGT CGG TGC TAT CTT GG-3' | 179-204 |
| M3 | 5'-CCG ACA ATA ACA TGA CTG AAA TGT ATA C-3' | 193-166 |
| M4 | 5'-GCT ATC TTG GGT TTC TTT AAT CAG GG-3' | 195-220 |
| M5 | 5'-GAT CAT TAT CCA CAC TGC TTC GAC GA-3' | 359-334 |
| M6 | 5'-ATC CAC ACT GCT TCG ACG AAA TGT AGA-3' | 352-326 |
| M7 | 5'-TAC GTG ACG AGC GGT GTG TTT AA-3' | 754-732 |
| M9 | 5'-AGA GAT AGA ATG TAA CGT GGA TCA T-3' | 378-354 |
| N1 | 5'-ATA GCT GTG ATG TTA AGG AGC ATA GGA A-3' | 1-24 |
| N2 | 5'-GAA GTT ATG GTT TTG GGC TCG TGA GT-3' | 121-96 |
| O1 | 5'-TGT GTA ACA GGG AGT CTA GCT TCA GTT-3' | 36-62 |
| R1 | 5'-ATT TAG CTC ACT GCG TAC TTA GGA TC-3' | 44-69 |
| R2 | 5'-AGA TAC AAG GAA CTT GAC AAG CAT TAC-3' | 43-17 |
| T1 | 5'-GAT TAG CTA CTA TCT TAT ACC TTA CTA CC-3' | 180-208 |
| T2 | 5'-GTT AGT ATA AGC ATA GAA CCA ATC CG-3' | 232-208 |
| T3 | 5'-AAG CAT AGA ACC AAT CCG GTA G-3' | 226-205 |
| SV1 | 5'-AGT TAT ACA GTT CTG GTT CGC GGA TCA T-3' | S (9-28) V (154-161) |
| SV2 | 5'-ATG ATC CGC GAA CCA GAA CTG TAT AAC T-3' | S (20-1) V (161-154) |
| V1 | 5'-TGG ACT GCT TAA GAC AGC TAA AAG T-3' | 45-21 |
| V2 | 5'-TGG TTG TCT GTA TCT CAT AAC TGG A -3' | 0 |

**Table S2. BLAST comparison of *Toxoplasma gondii* mtDNA sequence blocks to other publicly available apicomplexan mtDNA or genome sequences**

| *P. falciparum* | | | | *E. tenella* | | | | *T. gondii* | | *N. caninum* | | | | | *Hammondia* | | | | | *S. neurona* | | | | |
|---|---|---|---|---|---|---|---|---|---|---|---|---|---|---|---|---|---|---|---|---|---|---|---|---|
| mtDNA | | Tg Block | | mtDNA | | Tg Block | | | | Contig | | | Tg Block | | oxoDB.org Contig/NCBI sequenc | | | Tg Block | | ToxoDB.org Contig/NCBI sequence | | | Tg Block | |
| Start | Stop | Start | Stop | Start | Stop | Start | Stop | Sequence Block | Length | Contig | Start | Stop | Start | Stop | Contig/ Sequence ID | Start | Stop | Start | Stop | Contig/Sequence ID | Start | Stop | Start | Stop |
| 4164 | 4364 | 10 | 192 | 727 | 912 | 4 | 189 | A* | 196 | Contig8315 | 9607 | 9802 | 1 | 196 | KL545313 | 167 | 362 | 1 | 196 | KT207462.1 | 640 | 741 | 1 | 102 |
| not found | | | | not found | | | | B* | 40 | Contig8315 | 2644 | 2683 | 1 | 40 | KL544033 | 659216 | 659255 | 1 | 40 | not found | | | | |
| 2383 | 2571 | 6 | 194 | | | | | | | | | | | | | | | | | | | | | |
| 2614 | 3048 | 234 | 668 | 1518 | 2561 | 6 | 1049 | C* | 1050 | Contig8315 | 7173 | 6124 | 1 | 1050 | JX473251.1 | 1493 | 2542 | 1 | 1050 | SnSN1_scaffold00666 | 5 | 973 | 81 | 1049 |
| 3079 | 3273 | 699 | 893 | | | | | | | | | | | | | | | | | | | | | |
| 3253 | 3402 | 894 | 1043 | | | | | | | | | | | | | | | | | | | | | |
| not found | | | | not found | | | | D | 82 | Contig8315 | 1992 | 2073 | 1 | 82 | KL544028 | 581003 | 581083 | 2 | 82 | SnSN1_scaffold01461 | 474 | 504 | 15 | 45 |
| 3504 | 3782 | 30 | 308 | 73 | 720 | 33 | 680 | E* | 683 | Contig8315 | 8924 | 9606 | 1 | 683 | JX473260.1 | 377 | 1059 | 1 | 683 | KT207462.1 | 1 | 639 | 45 | 683 |
| 3783 | 4151 | 312 | 680 | | | | | | | | | | | | | | | | | | | | | |
| 5864 | 5911 | 19 | 65 | 3799 | 3879 | 19 | 99 | F^ | 179 | Contig8315 | 2855 | 3033 | 1 | 179 | KL544045 | 676029 | 675851 | 1 | 179 | SnSN1_scaffold01461 | 593 | 508 | 19 | 104 |
| 5577 | 5772 | 54 | 245 | 3002 | 2965 | 12 | 49 | H^ | 447 | Contig8315 | 9987 | 10433 | 1 | 447 | JX473270.1 | 1068 | 631 | 1 | 438 | sneu_contig02982 | 705 | 376 | 1 | 330 |
| | | | | 5593 | 5780 | 57 | 243 | | | | | | | | | | | | | | | | | |
| | | | | 6072 | 6094 | 25 | 47 | I^ | 204 | CADU01000139 | 1565 | 1362 | 1 | 204 | JX473262.1 | 265 | 468 | 1 | 204 | SnSN1_scaffold01461 | 256 | 360 | 1 | 101 |
| not found | | | | 3160 | 3224 | 92 | 156 | | | | | | | | | | | | | | 801 | 905 | 98 | 202 |
| 5877 | 5845 | 13 | 45 | 4418 | 4441 | 2 | 25 | J* | 85 | Contig8315 | 1627 | 1543 | 1 | 85 | KL544027 | 3083220 | 3083136 | 1 | 85 | sneu_scaffold00016 | 462040 | 461960 | 5 | 85 |
| 395 | 505 | 156 | 271 | 3599 | 3566 | 81 | 114 | | | | | | | | | | | | | | | | | |
| | | | | 3520 | 3480 | 105 | 145 | K^ | 445 | Contig8315 | 5443 | 5887 | 1 | 445 | JX473261.1 | 469 | 913 | 1 | 445 | SnSN1_scaffold01922 | 324 | 205 | 155 | 274 |
| 629 | 662 | 410 | 443 | 3295 | 3264 | 146 | 275 | | | | | | | | | | | | | | | | | |
| | | | | 5371 | 5499 | 412 | 443 | | | | | | | | | | | | | | | | | |
| 1333 | 1256 | 77 | 154 | 4328 | 4414 | 71 | 157 | L* | 159 | Contig8315 | 1786 | 1628 | 1 | 159 | KL554933 | 156 | 314 | 1 | 159 | SnSN1_scaffold02568 | 348 | 467 | 38 | 157 |
| 1060 | 911 | 54 | 203 | 4625 | 4786 | 51 | 212 | | | | | | | | | 1056505 | 1056602 | 1 | 104 | SnSn1_scaffold00839 | 522 | 680 | 60 | 218 |
| 1520 | 1613 | 384 | 477 | 2701 | 2898 | 397 | 593 | | | | | | | | | 293595 | 294050 | 104 | 560 | | 1134260 | 1134101 | 427 | 586 |
| 382 | 285 | 491 | 589 | 3447 | 3379 | 627 | 695 | M*^ | 754 | Contig8315 | 3787 | 3034 | 1 | 754 | KL544043 | 881359 | 881102 | 478 | 742 | sneu_scaffold00012 | 1134010 | 1134052 | 629 | 671 |
| 1659 | 1675 | 736 | 752 | 5937 | 5892 | 708 | 752 | | | | | | | | | 275663 | 275757 | 660 | 754 | | 1134789 | 1134750 | 715 | 754 |
| not found | | | | not found | | | | N | 166 | Contig8315 | 4612 | 4777 | 1 | 166 | JX473266.1 | 484 | 649 | 1 | 166 | sneu_contig02982 | 296 | 239 | 61 | 118 |
| low evalue- small match | | | | not found | | | | O^ | 86 | Contig8315 | 2769 | 2854 | 1 | 86 | KL558884 | 1350452 | 1350537 | 1 | 86 | low evalue- small match | | | | |
| low evalue- bad match | | | | not found | | | | P | 184 | Contig8315 | 9803 | 9986 | 1 | 184 | JX473270.1 | 447 | 630 | 1 | 184 | SnSN1_scaffold00004 | 5274306 | 5274250 | 128 | 184 |
| 2526 | 2564 | 2 | 40 | low evalue- bad match | | | | Q* | 65 | Contig8315 | 1478 | 1542 | 1 | 65 | KL544030 | 303803 | 303867 | 1 | 65 | low evalue- small match | | | | |
| low evalue- small match | | | | not found | | | | R | 85 | Contig8315 | 4522 | 4438 | 1 | 85 | KL544067 | 90510 | 90594 | 1 | 85 | SnSN1_scaffold00050 | 183223 | 183141 | 2 | 84 |
| 2173 | 2376 | 1 | 204 | 1308 | 1511 | 1 | 204 | S* | 205 | Contig8315 | 922 | 1126 | 1 | 205 | JX473255.1 | 733 | 937 | 1 | 205 | SnSN1_scaffold02197 | 247 | 450 | 1 | 204 |
| 4446 | 4619 | 99 | 272 | 922 | 1122 | 3 | 203 | T* | 279 | Contig8315 | 4878 | 5155 | 1 | 278 | JX473250.1 | 848 | 1123 | 1 | 276 | SnSN1_scaffold01547 | 717 | 487 | 3 | 233 |
| 594 | 623 | 7 | 35 | 3557 | 3524 | 6 | 38 | | | | | | | | | 169532 | 169467 | 1 | 67 | | | | | |
| 5141 | 5034 | 93 | 200 | 6164 | 6211 | 238 | 285 | U^ | 354 | Contig8315 | 2186 | 2539 | 1 | 354 | KL544069 | | | | | SnSN1_scaffold01547 | 451 | 182 | 44 | 313 |
| 1996 | 1921 | 234 | 309 | 1 | 52 | 288 | 339 | | | | | | | | | 24962 | 24634 | 26 | 354 | | | | | |
| 2062 | 2172 | 51 | 161 | 1170 | 1304 | 24 | 158 | V* | 161 | Contig8315 | 761 | 921 | 1 | 161 | KL544045 | 675154 | 675314 | 1 | 161 | SnSN1_scaffold02197 | 139 | 243 | 54 | 158 |

* Denotes seqeunce blocks that part of a cytochrome gene; ^ Denotes sequence block contains rRNA sequences; low evalue: above 1E-07; small match: 14-17 bp

[green] Contig from *N. caninum* genome assembly (ftp://ftp.sanger.ac.uk/pub/pathogens/Neospora/caninum/NEOS.contigs.072303)

[red] NCBI accesion numbers

[lavender] Scaffold/Contig from t oxodb.org version 44 (tblastx used for sequence blocks that are part of a gene to identify matches in *S. neurona*)

[light blue] *E. tenella* mtDNA sequence AB564272.1 (tblastx used for sequence blocks that are part of a CDS)

[blue] *P. falciparum* mtDNA sequence M76611.1 (tblastx used for sequence blocks that are part of a CDS)

*N. canium* : Best matches were mostly to Contig8315

*Hammondia*: The latest genome assembly in Toxodb.org and NCBI were searched and the better of the two hits are reported. The NCBI matches are to reported gene sequences

For *N. caninum* and *Hammondia* only blastn results are reported

### Table S3 – Summary of Nanopore and Illumina sequences generated

| *T. gondii* ME49 Nanopore statistics | | | | | | |
|---|---|---|---|---|---|---|
| | **Genome size** | **# reads** | **Size Range (nt)** | **Total Length (nt)** | **Coverage** | **SRA** |
| Nuclear reads | 67 MB | 43,392 | 48-90,222 | 313,953,499 | 4.68 X | SRR9200762 |
| mtDNA reads | 5,909 (minimum) | 271 | 219-23,619 | 711,833 | | |

| *T. gondii* RH *Δuprt* Nanopore statistics | | | | | | |
|---|---|---|---|---|---|---|
| | **Genome size** | **# reads** | **Size Range (nt)** | **Total Length (nt)** | **Coverage** | **SRA** |
| Nuclear reads | 67 MB | 765,200 | 56-94,214 | 4,309,669,177 | 64.32 X | SRR9961591 |
| mtDNA reads | 5,909 (minimum) | 779 | 146-15,914 | 2,057,412 | | |

| *N. caninum* Nc-1 Nanopore statistics | | | | | | |
|---|---|---|---|---|---|---|
| | **Genome size** | **# reads** | **Size Range (nt)** | **Total Length (nt)** | **Coverage** | **SRA** |
| Nuclear reads | 60 MB | 8748 | 45-51,875 | 17,024,894 | 0.284X | SRR9200761 |
| mtDNA reads | 5,908 (minimum) | 117 | 374-15,580 | 315,061 | | |

| *T. gondii* ME49 Illumina (PE-150) statistics | | | | | | |
|---|---|---|---|---|---|---|
| | **Genome size** | **# reads** | | **Total Length (nt)** | **Coverage** | **SRA** |
| Total DNA reads | 67 MB | 327,219,513 | | 98,400,570,288 | 1,468 X | SRR6793863 |

### Continuation of Figure 7 – Plot of Nuclear Nanopore read length for *T. gondii* ME49 and *N. caninum* Nc-1.

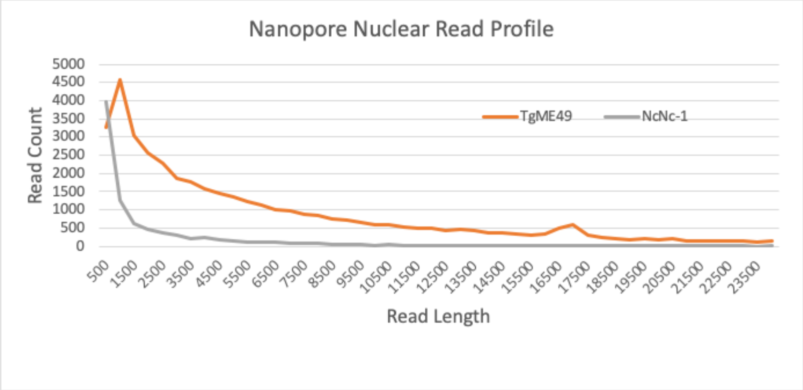

**Table S4. Mapping coverage depth to ascertain relative copy number.** *T. gondii* Illumina genomic reads from two strains were mapped to mt genes and several SB arrangements and their relative abundance with respect to the single copy nuclear gene, *gapdh* was calculated for each data set. Mapping or reads to the SBs required paired-ends at 100% identity. *The SBs A,B,D,E and K,M,V do not occur naturally and were manually constructed as negative controls. Mapping is still detected to the negative controls. Visualization of the mappings (data not shown) revealed paired-end reads mapping within single larger SBs, *e.g.* A, E, K and M. The exception to this rule was 33 paired-end reads that spanned the SBs K and M. Average coverage for paired-end reads mapping within the adjacent sequence block, M, was over 16,000 reads.

| Gene/Arrangement | ***T.gondii*** **ME49 SRR6793863** | ***T.gondii*** **RH-88 SRR521957** | **ME49** | **RH-88** |
|---|---|---|---|---|
| | **Average coverage** | | **Estimated # of copies** | |
| *gapdh* | 1195 | 90 | 1 | 1 |
| *coxI* | 377871 | 33066 | 316 | 369 |
| *coxIII* | 505607 | 46923 | 423 | 523 |
| *cob* | 355488 | 28027 | 297 | 313 |
| KOJE | 177968 | 14709 | 149 | 164 |
| VSRFOJ | 324164 | 24310 | 271 | 271 |
| CQJLVS | 237509 | 17999 | 199 | 201 |
| PIKpU | 260399 | 29081 | 218 | 324 |
| JLVDK | 191764 | 20383 | 160 | 227 |
| JQH | 122322 | 13516 | 102 | 151 |
| ABDE (neg control)* | 224646 | 18905 | 188 | 211 |
| KMV (neg control)* | 180577 | 16157 | 151 | 180 |

**Table S5. Evidence of *T. gondii* and *N. caninum* Sanger genomic and EST reads encoding Partial cytochromes genes**

| ***Toxoplasma gondii*** | | | |
|---|---|---|---|
| **Gene** | **Sequence blocks in EST read** | **GenBank Accession ID of EST** | **Trace Archive ID of a genomic read aligning with the EST read** |
| | A, T, V.D | CN617107.1 | ti:2057042422 |
| *coxI* | C, Q, J, L, V, S | DK934897.1 | ti:2056951560 |
| | V, S, R, F | CV701032.1 | ti:2057358855 |
| *coxIII* | V, L, J, B, M | CV654565.1 | ti:2057374420 |
| | B, M | DV110375.1 | ti:2064971134 |
| | A, T, V | CV653007.1 | ti:2057140800 |
| *cob* | A, T, E | DV109376.1 | ti:2057357849 |
| | E, A | CB384005.1 | ti:2057332917 |
| | K, O, J, E | CB752279.1 | ti:2057396732 |

*T. gondii* EST reads encoding portions of cytochromes and mtDNA SBs

| ***Neospora caninum*** | | | |
|---|---|---|---|
| **Gene** | **Sequence blocks in EST read** | **GenBank Accession ID of EST** | **Trace Archive ID of a genomic read aligning with the EST read** |
| | K, O, J, Q, C | CF275126.1 | ti:2158337065 |
| *coxI* | Q, J, L, V, S | CF937985.1 | ti:2158436519 |
| | C, S, V, L | CA857116.1 | ti:2158472021 |
| | V, S, R, F, N | CD537742.1 | ti:2158469753 |
| | Q, J, L, V, S | CF937985.1 | ti:2158436519 |
| *coxIII* | V, L, J, B, M | CF620007.1 | ti:2222083982 |
| | O, J, B, M | CF423132.1 | ti:2158457520 |
| | T, A, N | CF940825.1 | ti:2158465069 |
| *cob* | E, A, T | CF967503.1 | ti:2158382251 |
| | E, A | BF716988.1 | ti:2158435877 |

**Table S6. Annotation of mtDNA-specific *T. gondii* PRU RNA nanopore reads using the 21 sequence blocks.** Red= *coxI;* Green = *coxIII*; Blue = *cob* and Yellow indicates an anomaly in the sequence block. Red text = reverse orientation. * Full-length cytochrome transcript; # Cytochrome transcript at > 90% of gene length due to missing ends. Corrected sequences are available in Supplementary Dataset S8.

| TgNano_RNA_names | Tg_nanopore_RNAseq | Length | Annotation | Poly A |
|---|---|---|---|---|
| TgNano_RNA1 | e9333c9e-4041-4114-80bb-92524fcf2396 | 2880 | I K K O J B Mp U Kp I P A E | |
| TgNano_RNA2 | 023b1824-1bb4-4b96-9cda-63bdc8581de2 | 2339 | I P A N Fp R S V L B Mp U Kp | |
| TgNano_RNA3 | 968cba32-4c68-45df-abb3-2b2f4098deda | 2010 | U Mp B J L S C Q J | |
| TgNano_RNA4 | 4f149e5c-bbad-4179-87c9-ecb0a93716f6 | 1876 | N A T V D Kp U Mp B J O K | |
| TgNano_RNA5 | abeed298-7200-4999-ad78-56b85af0ac06 | 1729 | C S V L J | |
| TgNano_RNA6 | d7b561c2-da97-4eb2-8cd1-3eda76e8a39a | 1701 | M B J L V S R F | |
| TgNano_RNA7 | 5a8e1300-fa98-431a-8b43-69b9123c029e | 1671 | S V L J E A | |
| TgNano_RNA8 | 118c73fe-4caf-44c8-a547-1238d541b903# | 1616 | V S C Q | A |
| TgNano_RNA9 | a20cf716-5019-4803-a284-44d719afd29e | 1616 | C Q J L S C C | |
| TgNano_RNA10 | c566d00b-5be0-41a0-8e86-9de1ece7d3c1# | 1615 | O J E A T | B |
| TgNano_RNA11 | 7b66ffd3-a547-4dea-a733-26f681496e2c | 1570 | S C Q J | |
| TgNano_RNA12 | f6f4b33c-60e7-4f18-9f62-a9730d16cce1* | 1567 | V S C Q J | No |
| TgNano_RNA13 | 82ef02fe-dbbc-48a8-b84f-15f3ee08aa52 | 1563 | M B J O K I P | |
| TgNano_RNA14 | b500a94f-b453-41c9-bd3e-5775b5fd67de | 1563 | V S C | |
| TgNano_RNA15 | dda06ce5-a22c-4f08-8c1f-85d41848f90a# | 1527 | V S C Q J | No |
| TgNano_RNA16 | ce147cce-b308-4c26-a39c-33a35343a8dc# | 1523 | V S C Q | C |
| TgNano_RNA17 | d95c47c8-8409-491d-ab87-371a58f70548 | 1518 | V S C Q | |
| TgNano_RNA18 | eb151d53-ef7e-42b0-a038-c704b77d4a81# | 1511 | V S C Q | D |
| TgNano_RNA19 | 71371d1f-b682-4c97-a289-6718182d0198 | 1503 | C S V L J B | |
| TgNano_RNA20 | 7bb0da96-0483-4825-be30-24d67df50108 | 1503 | C S V T A | |
| TgNano_RNA21 | d5365e75-ab6d-4dfb-8936-58d5142ba9f7# | 1496 | V S C Q | E |
| TgNano_RNA22 | 9c45e8cb-0881-449e-905e-d49b4b447297# | 1493 | K J E A T | F |
| TgNano_RNA23 | a75c227b-9708-4aa9-b71f-46e8df45dd7b | 1465 | U Kp D V L J E A | |
| TgNano_RNA24 | d9cee345-5ec7-4652-85b5-d58cb2e1c300 | 1461 | H P A N Fp R S V | |
| TgNano_RNA25 | a1f14a9d-7c60-4779-8154-e2d15ca6c920 | 1452 | Kp I P A E | |
| TgNano_RNA26 | fc6a6310-d3a3-4edc-be55-e2976f642a31* | 1449 | O K D V L J B M | No |
| TgNano_RNA27 | 9508b00a-f738-481e-a614-723378e57b90 | 1376 | C S V L M | |
| TgNano_RNA28 | 1b6f35e2-9a58-4885-8821-fb90e13fe21c# | 1374 | K D V L J B M | G |
| TgNano_RNA29 | cf4c1453-4a29-41af-b2f5-e57c0150ebae* | 1356 | O J E A T | |
| TgNano_RNA30 | b844257b-7e20-4fff-a423-0863f9c25b99 | 1344 | S C Q | |

**A** -After TAA - **CACAATAGAACT**AAAAAAAAAAAAAAAAAAAAAAAAAAAAAAAAAAAAAAAAAATCCCTCCATCCATCTACTCTCATTCCATCATCTCAACTATTTCCATTTATTTC CTTTTCCTCTTCATTCCTAACCTACCATTCCCTCATCAAAACAAAAAAAAAAAAAAAAAAAAAAAAAAAAAAAAAAAAAAAAAAAAACCAG

**B -** Sequence block T not till *cob* end

**C** -After TAA - **CACAATAGAACT**AAAAAAAAAAACCCATCCCACATCTCATCACTCCCTTATCATCACTTATTAACTTCCATTTCAACCAACCCTACAACCTCAAATCCCACA

**D** - Ends 11 nt early, then TAGATCGGAAGAGCACACGTCTGAACTCCAGTCCTTACCCAACTTTCTCATAAACATAAATCTATATCAACCTGCCCTATCCTTAGAATCCCAACT

**E** - After TAA - CACAATAGAACTAAAAAAAAAAAAACCCACCCTCATCCCATACTTTGCAATGCCTATATCTATCCCTAGATA

**F** - Sequence block T not complete to *cob* end

**G** - Sequence block M not complete to end of *coxIII*

**Table S7. Examples of *T. gondii* DNA Nanopore reads with anomalies in sequence blocks.** Region with anomalies are highlighted in yellow Red text = reverse orientation. These are a subset of reads shown in Fig. S6.

| ***T. gondii* Nanopore Read ID** | **Annotation of Sequence Blocks** | | | | | | | | | | | | | | | | | | | | | | | | | | |
|---|---|---|---|---|---|---|---|---|---|---|---|---|---|---|---|---|---|---|---|---|---|---|---|---|---|---|---|
| e32f2dad-4608-49ff-b62e-379af84de712 | U | Kp | I | P | A | E | J | L | V | D | K | O | J | Q | H | P | A | E | T | V | S | R | F | O | J | B | M |
| 4f9137f3-d610-44a1-ad50-da462668413d | U | Mp | B | J | L | V | D | K | J | Q | H | P | A | E | J | O | F | M | | | | | | | | | |
| fadfacb2-e251-4149-bc9f-c297168a76b2 | S | R | F | F | M | B | J | O | F | M | M | B | J | O | K | D | | | | | | | | | | | |
| 2b941d62-dffb-41f4-bd3a-74d872421b0f | V | L | J | Q | H | P | A | E | J | L | V | S | R | F | O | J | B | Mp | U | Kp | D | V | L | J | B | Mp | U |
| c61269fa-fa08-4679-8983-f1a01a1635e4 | F | O | J | E | A | P | H | Q | J | L | V | D | Kp | U | M | B | J | L | Q | C | S | V | J | L | | | |
| 734acb5d-d7a7-4299-a6db-1032095a98d6 | Q | H | P | A | N | Fp | M | B | J | O | K | D | V | L | J | B | M | U | Kp | D | V | L | J | Q | C | S | V |

### Table S8. Annotation of REP elements with *T. gondii* sequence blocks

| REP element | GenBank ID | Annotation of the read using the 21 *T. gondii* mtDNA sequence blocks | | | | | | | | | | | |
|---|---|---|---|---|---|---|---|---|---|---|---|---|---|
| REP1 | X60240.1 | J (48-1) | L | V | S | R | F | O | J (3-48) | | | | |
| REP2 | X60241.1 | J (47-1) | L | V | D | Kp | U | Mp | B | J (85-43) | | | |
| REP3 | X60242.1 | J (48-1) | L | V | S | R | Fp | N | A | P | H | Q | J (85-43) |

Red font SBs in reverse orientation

Parentheses indicate the base pairs covered on each end (the SBs are incomplete)
